# Distributed harmonic patterns of structure-function dependence orchestrate human consciousness

**DOI:** 10.1101/2020.08.10.244459

**Authors:** Andrea I. Luppi, Jakub Vohryzek, Morten L. Kringelbach, Pedro A.M. Mediano, Michael M. Craig, Ram Adapa, Robin L. Carhart-Harris, Leor Roseman, Ioannis Pappas, Alexander R.D. Peattie, Anne E. Manktelow, Barbara J. Sahakian, Paola Finoia, Guy B. Williams, Judith Allanson, John D. Pickard, David K. Menon, Selen Atasoy, Emmanuel A. Stamatakis

## Abstract

A central question in neuroscience is how consciousness arises from the dynamic interplay of brain structure and function. Departing from the predominant location-centric view in neuroimaging, here we provide an alternative perspective on the neural signatures of human consciousness: one that is intrinsically centered on how the distributed network architecture of the human structural connectome shapes functional activation across scales. We decompose cortical dynamics of resting-state functional MRI into fundamental distributed patterns of structure-function association: the harmonic modes of the human structural connectome. We contrast wakefulness with a wide spectrum of states of consciousness, spanning chronic disorders of consciousness but also pharmacological perturbations of consciousness induced with the anaesthetic propofol and the psychoactive drugs ketamine and LSD. Decomposing this wide spectrum of states of consciousness in terms of “connectome harmonics” reveals a generalisable structure-function signature of loss of consciousness, whether due to anaesthesia or brain injury. A mirror-reverse of this harmonic signature characterises the altered state induced by LSD or ketamine, reflecting psychedelic-induced decoupling of brain function from structure. The topology and neuroanatomy of the human connectome are crucial for shaping the repertoire of connectome harmonics into a fine-tuned indicator of consciousness, correlating with physiological and subjective scores across datasets and capable of discriminating between behaviourally indistinguishable sub-categories of brain-injured patients, tracking the presence of covert consciousness. Overall, connectome harmonic decomposition identifies meaningful relationships between neurobiology, brain function, and conscious experience.

## Introduction

Understanding the neural underpinnings of human consciousness is a major challenge of contemporary neuroscience (Koch et al., 2016). Converging evidence suggests that consciousness is supported by a dynamic repertoire of brain activity (Barttfeld et al., 2015; Campbell et al., 2020; Demertzi et al., 2019; Golkowski et al., 2019; Gutierrez-Barragan et al., 2021; Huang et al., 2016, 2020; Luppi et al., 2019, 2021b, 2021a; Noirhomme et al., 2010; Tanabe et al., 2020). These discoveries raise the question of how the rich dynamics that support consciousness can arise from a fixed network of anatomical connections – the brain’s structural connectome (Abdelnour et al., 2014; Atasoy et al., 2018a; Cabral et al., 2017; Deco et al., 2011, 2013; Fukushima et al., 2018; Honey et al., 2007; Sporns et al., 2005; Xie et al., 2021). Initial progress on this question has provided fundamental insights into loss of consciousness and its signatures: when consciousness is lost, the pattern of co- fluctuations between regional BOLD time-series (“functional connectivity”) becomes more similar to the pattern of anatomical connections between regions (Barttfeld et al., 2015; Demertzi et al., 2019; Gutierrez-Barragan et al., 2021; Hahn et al., 2020; Lee et al., 2019; Tagliazucchi et al., 2016; Uhrig et al., 2018).

However, existing investigations of structure-function correspondence during altered states of consciousness have typically relied on correlation or distance metrics, which do not account for the inherently asymmetric relationship between brain structure and function: the organization of anatomical connections guides and constrains the propagation of functional signals, but not vice-versa^1^.

Additionally, existing approaches quantify structure-function correspondence at a single scale, even though brain structure and function both exhibit multi-scale, hierarchical network organization (Petersen and Sporns, 2015).

Here, we capitalize on the emerging mathematical framework of connectome harmonic decomposition (CHD) to overcome both of these limitations, by generalising the well-known Fourier transform to the network structure of the human brain. The traditional Fourier transform re-represents a signal from the time domain to the domain of temporal harmonic modes (Figure 1A). The signal is decomposed into a new set of basis functions: temporal harmonics (sinusoidal waves), each associated with a specific temporal frequency (Figure 1B). Generalising this mathematical principle, CHD uses the harmonic modes of the human structural connectome to perform an analogous change of basis functions (Figure 1C). Functional brain signals are re-represented from the spatial domain, to the domain of “connectome harmonics”: distributed patterns of activity, each associated with a specific spatial frequency, from coarse- to fine-grained (Atasoy et al., 2016, 2017, 2018a) (Figure 1D).

**Figure 1.**
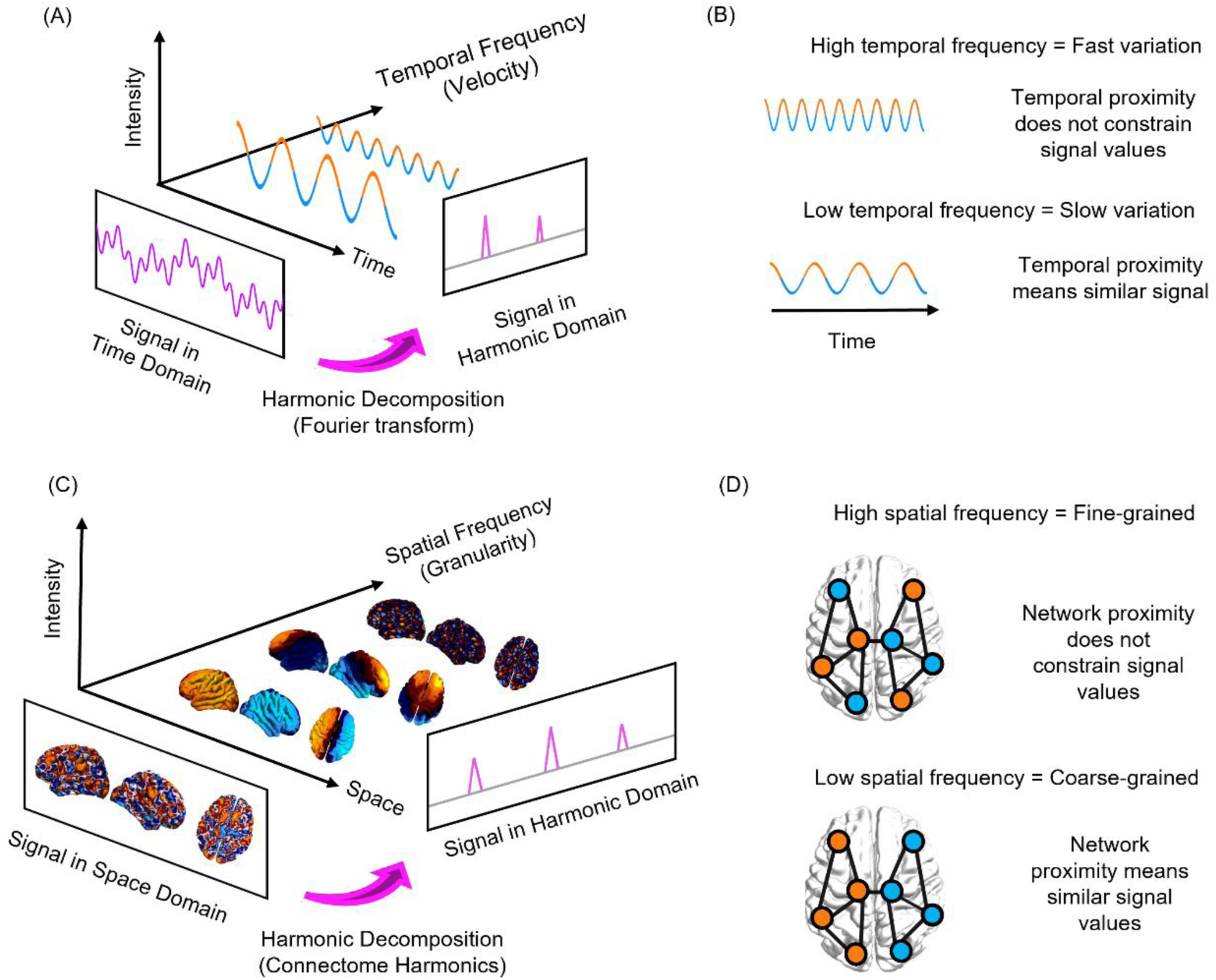
Connectome harmonic decomposition generalises the Fourier transform to the network structure of the human brain. (A) In traditional Fourier analysis, a signal in the time domain (represented in terms of sequential time-points) is decomposed into temporal harmonics of different frequency, thereby re-representing it in terms of a new set of basis functions. (B) High-frequency temporal harmonics correspond to fast-changing signals, such that data-points may have very different values even if they are close in time; in contrast, low-frequency temporal harmonics correspond to signals that vary slowly over time, such that temporally contiguous data- points have similar values, reflecting greater time-dependence of the signal. (C) In connectome harmonic decomposition, a signal in the space domain (represented in terms of BOLD activation at discrete spatial locations over the cortex) is decomposed into harmonic modes of the human structural connectome, providing a new set of basis functions in terms of whole-brain distributed patterns of activity propagation at different scales, from global patterns of smooth variation along geometrical axes (left–right and anterior–posterior being the most prominent) to increasingly complex and fine-grained patterns. Note that here, frequency is not about time, but about spatial scale (granularity). (D) Low-frequency (coarse-grained) connectome harmonics indicate that the spatial organisation of the functional signal is closely aligned with the underlying organisation of the structural connectome: nodes that are highly interconnected to one another exhibit similar functional signals to one another (indicated by colour). High-frequency (fine-grained) patterns indicate a divergence between the spatial organisation of the functional signal and the underlying network structure, whereby nodes may exhibit different functional signals even if they are closely connected in the structural network.

CHD is appealing for two reasons. Mathematically, it directly re-expresses functional signals in terms of their dependence on the underlying structural connectome. By direct analogy with the Fourier transform (Figure 1B), low- frequency (coarse-grained) connectome harmonics indicate that the functional signal is closely constrained by the underlying organisation of the structural connectome: nodes that are highly interconnected to one another exhibit similar functional signals to one another. In turn, high-frequency (fine-grained) connectome harmonics indicate a divergence between the spatial organisation of the functional signal and the underlying network structure: nodes may exhibit different functional signals even if they are closely connected in the structural network (Figure 1D). Therefore, just like temporal harmonics reflect time-dependence in the signal, so connectome harmonics quantify how brain activity is constrained by the underlying structural network on which it unfolds.

More broadly, CHD provides an alternative approach to conceptualize brain function in terms of distributed activity. The dominant perspective in neuroimaging views brain activity in terms of discrete, spatially localized signals. Operating within this spatially-localised framework, previous studies have sought to identify neural correlates of consciousness in terms of regional changes: whether pertaining to the intrinsic properties of a region, or its relationship with other regions (e.g., connectivity-based approaches). This endeavor has driven major progress in our understanding of consciousness and its neural bases (Adapa et al.; Bonhomme et al., 2016; Boveroux et al., 2010; Craig et al., 2020; Demertzi et al., 2015; Guldenmund et al., 2016; Hannawi et al., 2015; Huang et al., 2020; Kafashan et al., 2016; Luppi et al., 2019, 2021b; MacDonald et al., 2015; Palanca et al., 2015; di Perri et al., 2016; Di Perri et al., 2018; Ranft et al., 2016; Spindler et al., 2021; Stamatakis et al., 2010; Threlkeld et al., 2018; Vanhaudenhuyse et al., 2010; Vatansever et al., 2020). However, localized and distributed function co-exist in the brain, and its regions are intricately interconnected: local perturbations can have wide-ranging repercussions (“diaschisis”) (E Carrera, 2014), highlighting the limits of the location-centric view. Conceptually, the mathematical analogy between CHD and the Fourier transform highlights that viewing brain activity in terms of connectome harmonics (distributed patterns of different spatial scale) is just as legitimate as viewing it in terms of discrete spatial locations (Figure 1C): the two approaches provide perspectives that are neither redundant nor antithetical, but rather complementary.

This raises a pressing question: what insights are we missing out on as a field, by limiting ourselves to the spatially-localised view of brain function? The central hypothesis of this work is that the connectome harmonic view of brain activity will provide insights about consciousness that are complementary to the spatially-localised perspective, which has dominated neuroimaging research to date.

Crucially, loss of consciousness can occur through different mechanisms, ranging from transient pharmacological interventions to chronic neuroanatomical injuries. To make progress in our understanding of consciousness, it is imperative to identify signatures of consciousness *per se,* which generalise across different neurophysiological states. Therefore, here we leverage connectome harmonic decomposition of human functional MRI data to investigate the connectome harmonic signatures of loss of consciousness induced by different means: acutely, with the intravenous anaesthetic propofol (Stamatakis et al., 2010); and in brain-injured patients with chronic disorders of consciousness (Luppi et al., 2019). Additionally, a comprehensive characterisation of human consciousness should account for different kinds of perturbations, whereby consciousness is not lost but rather subjectively altered – such as the states induced by the classic psychedelic LSD and the “atypical psychedelic” ketamine. Since previous work has shown that CHD can identify a consistent neural signature across the serotonergic psychedelics LSD and psilocybin (Atasoy et al., 2017, 2018b, 2018a), extending this signature to ketamine (which is an N-methyl-D-aspartate receptor antagonist (Olney et al., 1999)) will provide critical insights into CHD’s ability to identify generalizable links between brain dynamics and alterations of consciousness.

Overall, here we aim to depart from the predominant location-centric view in neuroimaging and provide an alternative, mathematically principled perspective on the neural signatures of consciousness: one that is intrinsically centered on how the distributed network architecture of the human structural connectome shapes neural activation across scales.

## Materials and Methods

### Generalising the Fourier transform to the network structure of the human connectome: theoretical background

Connectome harmonic decomposition (CHD) generalises the mathematics of the Fourier transform to the network structure of the human brain. The traditional Fourier transform operates in the temporal domain (Figure 1A): decomposition into temporal harmonics quantifies to what extent the signal varies slowly (low-frequency temporal harmonics) or quickly (high-frequency temporal harmonics) over time (Figure 1B). Analogously, CHD re-represents a spatial signal in terms of harmonic modes of the human connectome, so that the spatial frequency (granularity) of each connectome harmonic quantifies to what extent the organization of functional brain signals deviates from the organization of the underlying structural network (Figure 1C,D). Therefore, CHD is fundamentally different from, and complementary to, traditional approaches to functional MRI data analysis. This is because CHD does not view functional brain activity as composed of signals from discrete spatial locations, but rather as composed of contributions from distinct spatial frequencies: each connectome harmonic is a whole-brain pattern with a characteristic spatial scale (granularity) – from an entire hemisphere to just a few millimetres.

On one hand, this means that CHD is unsuitable to address questions pertaining to spatial localisation and the involvement of specific neuroanatomical regions; such questions have been extensively investigated within the traditional framework of viewing brain activity in terms of spatially discrete regions, and several previous studies have implicated specific neuroanatomical regions in supporting consciousness (Boveroux et al., 2010; Hannawi et al., 2015; Huang et al., 2020; Luppi et al., 2019, 2021b; MacDonald et al., 2015; di Perri et al., 2018; Ranft et al., 2016; Vanhaudenhuyse et al., 2010). On the other hand, CHD enables us to consider how brain activity across states of consciousness is shaped by the brain’s distributed network of structural connections, reflecting the contribution of global patterns at different spatial scales - each arising from the network topology of the human connectome. We emphasise that neither approach is inherently superior, but rather they each provide a unique perspective on brain function: one localised, the other distributed. The traditional spatially-resolved approach and our frequency-resolved approach are two synergistic sides of the same coin.

Crucially, the use of the word “frequency” for both connectome harmonic decomposition and traditional Fourier analysis should not give the misleading impression that our analyses are redundant with previous literature based on traditional Fourier analysis in the domain of temporal frequencies (just like the “connectome harmonic energy” defined below is entirely different from metabolic energy). On the contrary, signal decomposition in terms of temporal frequencies (i.e., Fourier analysis) and connectome-based frequencies (i.e., CHD) operate in entirely separate domains and provide very different information about the signal (temporal dependence versus spatial dependence on the connectome network structure). Indeed, this means that even the most suitable neuroimaging modalities for each analysis are different: CHD relies on fMRI data with high spatial resolution, but which have a restricted content of temporal frequencies (the BOLD signals used here were all band-pass filtered in the low-frequency range as part of standard denoising procedures), whereas Fourier investigations of consciousness require high temporal resolution and are therefore typically performed on electro- or magneto-encephalography data. Please note that throughout this article, unless otherwise specified, our use of the word “frequency” refers to the frequency of connectome harmonics (spatial granularity, from fine-grained to coarse-grained).

### High-resolution structural connectome

Whereas alternative approaches to harmonic mode decomposition rely exclusively on white-matter connectivity between macroscopic brain regions defined by sub- dividing the brain into discrete parcels (Abdelnour et al., 2014; Glomb et al., 2020; Huang et al., 2018a; Kuceyeski et al., 2016; Preti and Van De Ville, 2019), CHD combines long-range white-matter connections with local connectivity within the grey matter on a continuous cortical surface, thereby achieving a representation of the human structural connectome at the highest resolution available for MR imaging (Atasoy et al., 2016; Naze et al., 2021).

For consistency with previous work employing connectome harmonic decomposition, for our main results we used connectome harmonics obtained from the same reconstruction of the human structural connectome used by Atasoy and colleagues (Atasoy et al., 2016, 2017). However, note that we also replicated our results using two alternative reconstructions of the human connectome at higher resolution (described below), including one obtained from aggregating 985 subjects from the Human Connectome Project (HCP) (van Essen et al., 2013): arguably one of the most representative reconstructions of the human structural connectome available to date, corresponding to a nearly 100-fold increase in sample size with respect to previous CHD studies (Atasoy et al., 2016, 2017).

### Connectome reconstruction

The workflow was the same as described in previous work by Atasoy and colleagues (Atasoy et al., 2016, 2017), who derived a high-resolution human structural connectome from derived from DTI and structural MRI data from an independent sample of 10 HCP subjects (six female, age 22–35), preprocessed according to minimal preprocessing guidelines of the HCP protocol. For each of these HCP subjects, Freesurfer (http://freesurfer.net) was used to reconstruct the cortical surfaces of each hemisphere at the interface of white and grey matter, based on the 0.7mm resolution data from T1-weighted MRI. This resulted in a representation of 18,715 cortical surface vertices for each subject. Subsequently, deterministic tractography was used to reconstruct long-range white matter fibres. After co-registering each subject’s diffusion imaging and cortical surface data, each of the 18,715 vertices of the reconstructed cortical surface was used as a centre to initialise eight seeds for deterministic tractography, implemented with the MrDiffusion tool (http://white.stanford.edu/newlm/index.php/MrDiffusion). Tracking was terminated when fractional anisotropy (FA) was below a threshold of 0.3, with 20mm minimum tract length, and setting 30 degrees as the maximum allowed angle between consecutive tracking steps (Atasoy et al., 2016, 2017).

The structural connectome of each subject was then represented as a binary adjacency matrix *A,* treating each cortical surface vertex as a node: for each pair *i* and *j* of the *n* = 18,715 cortical surface grey matter nodes, *A_ij_* was set to 1 if there was a white matter tract connecting them, as estimated from the deterministic tractography step described above (in order to account for long-range connections); or if they were adjacent in the gray matter cortical surface representation, thereby accounting for the presence of local (1-6mm) connections within the grey matter (the importance of accounting for short-range grey-matter connections in addition to long-range white matter tracts was demonstrated in a recent study (Naze et al., 2021)). If neither long-range nor short-range connections between *i* and *j* existed, *A_ij_* was set to 0. This procedure resulted in a symmetric (undirected) binary matrix (Atasoy et al., 2016).

The individual adjacency matrices were then averaged across the 10 HCP subjects to obtain a group-average matrix *A*, encoding a representative structural conenctome We then define the degree matrix D of the graph as:

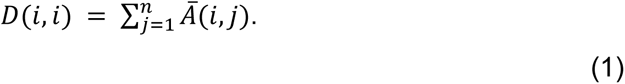

### Extraction of connectome harmonics

Following (Atasoy et al., 2016; Chung, 1997), we compute the symmetric graph Laplacian Δ_*G*_ on the group-average adjacency matrix *A* that represents the human connectome, in order to estimate the Laplacian (discrete counterpart of the Laplace operator Δ (Chung, 1997)) of the human structural connectome:

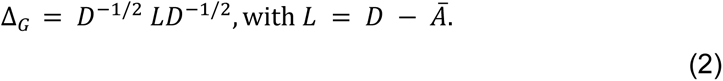

We then calculate the connectome harmonics *φ_k_*, k ∈ {1, …, 18,715} by solving the following eigenvalue problem:

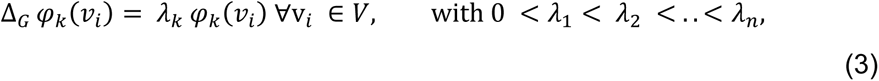

where *λ_k_*, *k* ∈ {1, …, n} is the corresponding eigenvalue of the eigenfunction *φ_k_*,

*V* is the set of cortical surface vertices and n represents the number of vertices. In other words, *λ_k_* and *φ_k_* are the eigenvalues and eigenvectors of the Laplacian of the human structural connectivity network, respectively. Therefore, if *φ_k_* is the connectome harmonic pattern of the *k*^th^ spatial frequency, then the corresponding eigenvalue *λ_k_* is a term relating to the intrinsic energy of the that particular harmonic mode. Crucially, we reiterate that the frequencies associated with each connectome harmonic are in the spatial rather than temporal domain, and should not be confused with the temporal frequencies identified by Fourier transform in the temporal domain (e.g. for denoising of timeseries).

### Connectome-harmonic decomposition of fMRI data

At each timepoint *t* ∈ {1,…,T} (corresponding to one TR), the preprocessed and denoised fMRI data (see below for details of these steps) were projected onto cortical surface coordinates by means of the Human Connectome Project Workbench *-volume-to-surface*-*mapping* tool. Then, the spatial pattern of cortical activity over vertices *v* at time *t,* denoted as *F_t_(v*), was decomposed as a linear combination of the set of connectome harmonics 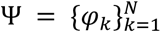:

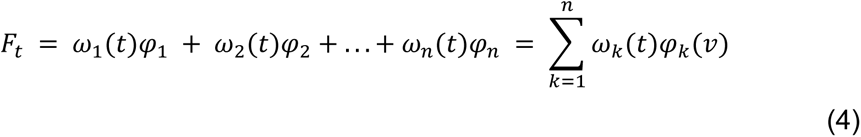

with the contribution *ω_k_*(*t*) of each connectome harmonic *φ_k_* at time *t* being estimated as the projection (dot product) of the fMRI data *F_t_(v*) onto *φ_k_*:

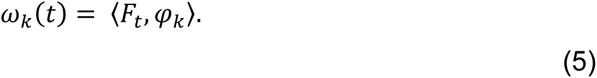

### Power and Energy of connectome harmonics

Once the fMRI cortical activation pattern at time *t* has been decomposed into a linear combination of connectome harmonics, the magnitude of contribution to cortical activity of each harmonic *φ_k_, k* ∈ {1, …, *n*} (regardless of sign) at any given timepoint *t* (*P*(*φ_k_*, *t*)), called its “power” for analogy with the Fourier transform, is computed as the amplitude of its contribution:

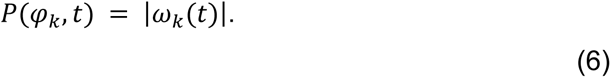

In turn, the normalized frequency-specific contribution of each harmonic *φ_k_*, k ∈ {1, …, *n*} at timepoint *t,* termed “energy”, is estimated by combining the strength of activation (power) of a particular connectome harmonic with its own intrinsic energy given by *λ_k_*^2^:

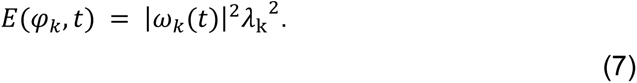

Consequently, total brain energy at time *t* is given by

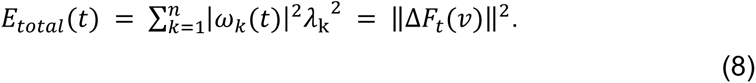

Since the Laplace operator Δ represents the amount of activity flow, the latter part of Equation 8 indicates that the total brain energy at a given point in time can be interpreted as the total cortical flow of neural activity at that time (Atasoy et al., 2017). A binned energy spectrum across subjects and time points is constructed by discretising the energy of connectome harmonics into 15 logarithmically-spaced frequency-specific bins, following previous work showing that this procedure can successfully highlight the connectome harmonic signatures of altered states of consciousness (Atasoy et al., 2017, 2018b) (Figure S1).

### Data-driven extraction of multivariate connectome harmonic signatures

Partial least squares (PLS, also known as Projection on Latent Spaces) is a multivariate statistical analysis used to identify relationships between one or more targets (Y) and a set of predictor variables X. This method extracts principal components as linear combinations of variables in each set that maximally covary with each other. In the present case, for each pair of states of consciousness under comparison, X was the matrix of 15 binned energy values (see above) for each subject (averaged over timepoints), and Y was the vector of binary classification between the two states (here, target vs baseline state of consciousness, e.g. anaesthetised vs awake, ketamine vs placebo, fMRI- vs fMRI+ DOC, etc) – making this an application of Partial Least Squares Discriminant Analysis (PLS-DA), owing to the binary nature of Y (Chuen Lee et al., 2018). The first principal component extracted by PLS-DA represents the single most discriminative pattern present in the data, in terms of distinguishing observations (subjects) belonging to the two different classes (states of consciousness).

### Diversity of connectome harmonic repertoire

To quantify the diversity of the repertoire of connectome harmonics recruited at each point in time, we start by observing that a diverse repertoire is one in which different harmonic modes contribute in different degrees to brain activity – neither one single mode dominating (which would correspond to a periodic oscillation, in analogy with the traditional Fourier transform) nor every mode contributing the same as every other mode (which would correspond to white noise). To capture this intuition, we quantify repertoire diversity in terms of the entropy of the distribution of connectome harmonic power (absolute strength of contribution to the cortical activation pattern) across all 18,715 connectome harmonics (i.e., binning was not used for this analysis). Specifically, to deal with continuous data (as in the present case) we rely on the Kozachenko approximation, as implemented in the Java Information Dynamics Toolbox (JIDT; http://jlizier.github.io/jidt/) (Lizier, 2014). We note that when dealing with continuous variables, entropy can have negative values (Cover and Thomas, 2005), but its interpretation remains the same: a more entropic distribution (i.e. having a value of entropy closer to positive infinity) will correspond to a more diverse repertoire. We calculate this entropy for each timepoint of each subject.

### High-resolution alternative reconstructions of the human connectome

To (Atasoy et al., 2016, 2017)demonstrate that our results are not fundamentally dependent on this specific operationalisation of the human connectome, we also used two alternative representative human connectomes. The first alternative connectome was constructed from multi-shell diffusion-weighted imaging data from 985 subjects of the HCP 1200 data release (http://www.humanconnectome.org/), each scanned for approximately 59 minutes. This represents a nearly 100-fold increase in sample size compared with the original connectome used for connectome harmonic decomposition (Atasoy et al., 2016, 2017). We refer to the human connectome constructed from these data as the HCP-985 connectome. The second alternative connectome was constructed from 32 healthy volunteers from the HCP database who were scanned for a full 89 minutes at Massachusetts General Hospital with high-resolution diffusion spectrum imaging, which can better resolve crossing fibers. We refer to this connectome as the MGH-32 connectome. Acquisition parameters for both groups are described in detail in the relative documentation (http://www.humanconnectome.org/) (Setsompop et al., 2013), and both dMRI datasets were preprocessed and made available as part of the freely available Lead-DBS software package (http://www.lead-dbs.org/).

For the reconstruction of long-range white matter tracts of each individual, we followed the procedures previously used on these data by (Deco et al., 2021): the diffusion data were processed using a generalized *q*-sampling imaging algorithm implemented in DSI Studio (http://dsi-studio.labsolver.org). A white-matter mask was obtained from segmentation of the T2-weighted anatomical images, which were co-registered to the b0 image of the diffusion data using SPM12. In each HCP participant, 200,000 fibres were sampled within the white-matter mask, using a tracking method that previously achieved the highest (92%) valid connection score among 96 methods submitted from 20 different research groups in a recent open competition (Maier-Hein et al., 2017). Finally, the fibres were transformed into standard Montreal Neurological Institute (MNI-152I) space using Lead-DBS (Horn and Blankenburg, 2016). The remaining procedures for obtaining individual connectomes and aggregating them into a group-average representative connectome, and subsequent connectome harmonic decomposition, were the same as described above. To ensure that our results were not unduly influenced by potential aliasing effects introduced by the use of high-resolution diffusion data, for both the HCP-985 and MGH-32 analyses we only used the first 14 logarithmically spaced bins (instead of 15 as for the previous analyses), showing that our results are not critically dependent on the precise number of bins.

### Rotated harmonics

To demonstrate the importance of the neuroanatomical distribution of connectome harmonics, we assessed whether our results would replicate when using spherically rotated connectome harmonics, following a recently described approach (Glomb et al., 2021), based on freely available code (github.com/spin-test/spin-test) (Alexander-Bloch et al., 2018). After obtaining the connectome harmonics following the procedure described above, the corresponding surface maps were projected onto a spherical surface, and subsequently rotated by a random angle, before mapping back the rotated values onto the nearest vertex (ignoring parts of the corpus callosum that are mapped onto the cortical surface). Since we used multi-dimensional basis functions, we rotated the surface maps corresponding to each dimension by the same angle. Note that the resulting rotated maps are not orthonormal anymore, because each rotated map is symmetrised to preserve this important property. We then proceeded with the normal CHD analysis workflow as described above.

### Randomised connectome

To demonstrate the importance of the specific topology of the human connectome, obtained by combining local grey matter connectivity and long-range white matter fibres, we also tested whether our results would replicate when using harmonics obtained from a randomised connectome (Naze et al., 2021). Before performing Laplacian decomposition, the original connectome was therefore turned into a random network using the degree-preserving procedure implemented in the Brain Connectivity Toolbox (Rubinov and Sporns, 2010). Harmonics were then extracted from Laplacian eigendecomposition, and the full connectome harmonic decomposition pipeline was followed. As for the validation analyses using alternative operationalisations of the human connectome, for this analysis we also only used the first 14 logarithmically spaced bins.

### Propofol Dataset

Sixteen healthy volunteer subjects were recruited for scanning. The acquisition procedures are described in detail in a previous study (Stamatakis et al., 2010). In addition to the original 16 volunteers, data were acquired for nine additional participants using the same procedures, bringing the total number of participants in this dataset to 25 (11 males, 14 females; mean age 34.7 years, SD = 9.0 years). Ethical approval for these studies was obtained from the Cambridgeshire 2 Regional Ethics Committee, and all subjects gave informed consent to participate in the study. Volunteers were informed of the risks of propofol administration, such as loss of consciousness, respiratory and cardiovascular depression. They were also informed about more minor effects of propofol such as pain on injection, sedation and amnesia. In addition, standard information about intravenous cannulation, blood sampling and MRI scanning was provided. The GABA-ergic intravenous agent propofol is one of the most commonly used anaesthetic drugs, owing to the stability and predictability of its effects. Three target plasma levels of propofol were used - no drug (Awake), 0.6 mg/ml (Mild sedation) and 1.2 mg/ml (Moderate sedation). Scanning (rs-fMRI) was acquired at each stage, and also at Recovery; anatomical images were also acquired. The level of sedation was assessed verbally immediately before and after each of the scanning runs.

### Propofol study: infusion protocol

Propofol was administered intravenously as a “target controlled infusion” (plasma concentration mode), using an Alaris PK infusion pump (Carefusion, Basingstoke, UK). A period of 10 min was allowed for equilibration of plasma and effect-site propofol concentrations. Blood samples were drawn towards the end of each titration period and before the plasma target was altered, to assess plasma propofol levels. In total, 6 blood samples were drawn during the study. The mean (SD) measured plasma propofol concentration was 304.8 (141.1) ng/ml during mild sedation, 723.3 (320.5) ng/ml during moderate sedation and 275.8 (75.42) ng/ml during recovery. Mean (SD) total mass of propofol administered was 210.15 (33.17) mg, equivalent to 3.0 (0.47) mg/kg. Two senior anaesthetists were present during scanning sessions and observed the subjects throughout the study from the MRI control room and on a video link that showed the subject in the scanner. Electrocardiography and pulse oximetry were performed continuously, and measurements of heart rate, non-invasive blood pressure, and oxygen saturation were recorded at regular intervals.

### Propofol study: MRI Data Acquisition

The acquisition procedures are described in detail in the original study (Stamatakis et al., 2010). Briefly, MRI data were acquired on a Siemens Trio 3T scanner (WBIC, Cambridge). For each level of sedation, 150 rs-fMRI volumes (5 min scanning) were acquired. Each functional BOLD volume consisted of 32 interleaved, descending, oblique axial slices, 3 mm thick with interslice gap of 0.75 mm and in-plane resolution of 3 mm, field of view = 192x192 mm, repetition time = 2000 ms, acquisition time = 2 s, time echo = 30 ms, and flip angle 78. We also acquired T1- weighted structural images at 1 mm isotropic resolution in the sagittal plane, using an MPRAGE sequence with TR = 2250 ms, TI = 900 ms, TE = 2.99 ms and flip angle = 9 degrees, for localization purposes. During scanning we instructed volunteers to close their eyes and think about nothing in particular throughout the acquisition of the resting state BOLD data. Of the 25 healthy subjects, 15 were ultimately retained (7 males, 8 females): 10 were excluded, either because of missing scans (n=2), or due of excessive motion in the scanner (n=8, 5mm maximum motion threshold).

### Disorders of Consciousness Patient Dataset

As previously reported (Luppi et al., 2019, 2021c), 71 DOC patients were recruited from specialised long-term care centres from January 2010 to December 2015. Ethical approval for this study was provided by the National Research Ethics Service (National Health Service, UK; LREC reference 99/391). Patients were eligible to be recruited in the study if they had a diagnosis of chronic disorder of consciousness, provided that written informed consent to participation was provided by their legal representative, and provided that the patients could be transported to Addenbrooke’s Hospital (Cambridge, UK). The exclusion criteria included any medical condition that made it unsafe for the patient to participate, according to clinical personnel blinded to the specific aims of the study; or any reason that made a patient unsuitable to enter the MRI scanner environment (e.g. non-MRI-safe implants). Patients were also excluded based on significant pre- existing mental health problems, or insufficient fluency in the English language prior to their injury. After admission to Addenbrooke’s Hospital, each patient underwent clinical and neuroimaging testing (including task-based, resting-state, and anatomical scans), spending a total of five days in the hospital (including arrival and departure days). Neuroimaging scanning took place at the Wolfson Brain Imaging Centre (Addenbrooke’s Hospital, Cambridge, UK), and medication prescribed to each patient was maintained during scanning.

For each day of admission, Coma Recovery Scale-Revised (CRS-R) assessments were recorded at least daily. Patients whose behavioural responses were not indicative of awareness at any time, were classified as UWS. In contrast, patients were classified as being in a minimally conscious state (MCS) if they provided behavioural evidence of simple automatic motor reactions (e.g., scratching, pulling the bed sheet), visual fixation and pursuit, or localisation to noxious stimulation) (note that due to the limited size of our sample of MCS patients, we do not sub- divide these patients into MCS- and MCS+) (Bruno et al., 2011; Wannez et al., 2018). Since this study focused on whole-brain properties, coverage of most of the brain was required, and we followed the same criteria as in our previous studies (Luppi et al., 2019): before analysis took place, patients were systematically excluded if an expert neuroanatomist blinded to diagnosis judged that they displayed excessive focal brain damage (over one third of one hemisphere), or if brain damage led to suboptimal segmentation and normalisation, or due to excessive head motion in the MRI scanner (exceeding 3mm translation or 3 degrees rotation). Forty-one patients were excluded due to excessive brain damage and distortion preventing satisfactory segmentation and normalisation; 8 further patients due to excessive motion. A total of 22 adults (14 males, 8 females; age range 17-70 years; mean time post injury: 13 months) meeting diagnostic criteria for Unresponsive Wakefulness Syndrome/Vegetative State (N = 10) or Minimally Conscious State (N = 12) due to brain injury were included in this study (Table 1).

**Table 1:** Demographic information for patients with Disorders of Consciousness. CRS-R, Coma Recovery Scale-Revised; UWS, Unresponsive Wakefulness Syndrome; MCS, Minimally Conscious State; TBI, Traumatic Brain Injury; fMRI-, negative responders to mental imagery task; fMRI+, positive responders to mental imagery task; SMA, supplementary motor area; PPA, parahippocampal place area; PMC, pre-motor cortex.

| Sex | Age | Aetiology | Diagnoses | CRS-R Score | Tennis | Spat Nav | Classification |
| --- | --- | --- | --- | --- | --- | --- | --- |
| M | 46 | TBI | UWS | 6 | no evidence | no evidence | fMRI- |
| M | 57 | TBI | MCS | 12 | no evidence | no evidence | fMRI- |
| M | 46 | TBI | MCS | 10 | no evidence | no evidence | fMRI- |
| M | 35 | Anoxic | UWS | 8 | no evidence | no evidence | fMRI- |
| M | 17 | Anoxic | UWS | 8 | no evidence | positive | fMRI+ |
| F | 31 | Anoxic | MCS | 10 | no evidence | no evidence | fMRI- |
| F | 38 | TBI | MCS | 11 | positive | no evidence | fMRI+ |
| M | 29 | TBI | MCS | 10 | SMA +ve | PPA +ve | fMRI+ |
| M | 23 | TBI | MCS | 7 | SMA +ve | no evidence | fMRI+ |
| F | 70 | Cerebral bleed | MCS | 9 | no evidence | no evidence | fMRI- |
| F | 30 | Anoxic | MCS | 9 | PMC +ve | no evidence | fMRI+ |
| F | 36 | Anoxic | UWS | 8 | no evidence | PPA +ve | fMRI+ |
| M | 22 | Anoxic | UWS | 7 | no evidence | no evidence | fMRI- |
| M | 40 | Anoxic | UWS | 7 | no evidence | no evidence | fMRI- |
| F | 62 | Anoxic | UWS | 7 | no evidence | no evidence | fMRI- |
| M | 46 | Anoxic | UWS | 5 | no evidence | no evidence | fMRI- |
| M | 21 | TBI | MCS | 11 | no evidence | no evidence | fMRI- |
| M | 67 | TBI | MCS | 11 | SMA +ve | PPA +ve | fMRI+ |
| F | 55 | Hypoxia | UWS | 7 | no evidence | negative | fMRI- |
| M | 28 | TBI | MCS | 8 | positive | positive | fMRI+ |
| M | 22 | TBI | MCS | 10 | no evidence | negative | fMRI- |
| F | 28 | ADEM | UWS | 6 | no evidence | negative | fMRI- |

### Stratification of DOC Patients into fMRI+ and fMRI-

Patients were stratified into two groups based on their ability to perform volitional tasks (mental imagery) in the scanner (Craig et al., 2021; Luppi et al., 2021c). The mental imagery tasks used here have been previously used to assess the presence of covert consciousness in DOC patients (Monti et al., 2010; Owen et al., 2006) and their validity has been confirmed in healthy individuals (Fernández-Espejo et al., 2014). Patients were instructed to perform two mental imagery tasks. The first task involved motor imagery (“tennis task”): each patient was asked to imagine being on a tennis court swinging their arm to hit the ball back and forth with an imagined opponent. The second was a task of spatial imagery (“navigation task”): the patient was required to imagine walking around the rooms of their house, or the streets of a familiar city, and to visualise what they would see if they were there. Each task comprised five cycles of alternating imagery and rest blocks, each lasting 30 seconds. The two kinds of mental imagery blocks were cued with the spoken word “tennis” or “navigation”, respectively, whereas the resting blocks were cued with the word “relax”, corresponding to instructions for the patient to just stay still and keep their eyes closed. Univariate fMRI analysis was conducted on all 22 patients for both the motor and spatial mental imagery tasks (Craig et al., 2021). The analyses were performed using FSL version 5.0.9 (https://fsl.fmrib.ox.ac.uk/fsl/fslwiki/). The results of these analyses determined which patients would be placed in each classification condition. For each functional scan, a general linear model consisting of contrasting periods of rest and active imagery was computed. Results were considered significant at a cluster level of z > 2.3 (corrected p < 0.05 at the cluster level) (Craig et al., 2021; Monti et al., 2010). Patients who exhibited significantly greater brain activation in the appropriate regions (supplementary motor area (SMA) for the tennis task, and parahippocampal place area (PPA) for the navigation task, respectively) during either of the volitional mental imagery tasks than rest (i.e., those who exhibited evidence of being able to respond to the task) were deemed to be covertly conscious (N = 8); for brevity, we refer to these positive responders as “fMRI+”. Conversely, we refer to patients who did not respond to either task (negative responders), and who therefore did not exhibit detectable evidence of covert consciousness (N = 14), as “fMRI-”. (Table 1).

### DOC patients: MRI Data Acquisition

Resting-state fMRI was acquired for 10 minutes (300 volumes, TR=2000ms) using a Siemens Trio 3T scanner (Erlangen, Germany). Functional images (32 slices) were acquired using an echo planar sequence, with the following parameters: 3 x 3 x 3.75mm resolution, TR = 2000ms, TE = 30ms, 78 degrees FA. Anatomical scanning was also performed, acquiring high-resolution T1-weighted images with an MPRAGE sequence, using the following parameters: TR = 2300ms, TE = 2.47ms, 150 slices, resolution 1 x 1 x 1mm.

### Ketamine dataset

A total of 21 participants (10 males; mean age 28.7 years, SD = 3.2 years) were recruited via advertisements placed throughout central Cambridge, UK (Dandash et al., 2015). All participants underwent a screening interview in which they were asked whether they had previously been diagnosed or treated for any mental health problems and whether they had ever taken any psychotropic medications. Participants reporting a personal history of any mental health problems or a history of any treatment were excluded from the study. All participants were right-handed, were free of current of previous psychiatric or neurological disorder or substance abuse problems, and had no history of cardiovascular illness or family history of psychiatric disorder/substance abuse. The study was approved by the Cambridge Local Research and Ethics Committee, and all participants provided written informed consent in accordance with ethics committee guidelines. Participants were scanned (resting-state functional MRI and anatomical T1) on two occasions, separated by at least 1 week. On one occasion, they received a continuous computer-controlled intravenous infusion of a racemic ketamine solution (2 mg/ml) until a targeted plasma concentration of 100 ng/ml was reached. This concentration was sustained throughout the protocol. A saline infusion was administered on the other occasion. Infusion order was randomly counterbalanced across participants.

### Ketamine study: infusion protocol

The infusion was performed and monitored by a trained anesthetist (RA) who was unblinded for safety reasons, but who otherwise had minimal contact with participants. At all other times, participants were supervised by investigators blinded to the infusion protocol. The participants remained blinded until both assessments were completed. Bilateral intravenous catheters were inserted into volunteers’ forearms, one for infusion, and the other for serial blood sampling. We used a validated and previously implemented (Corlett et al., 2006) three-compartment pharmacokinetic model to achieve a constant plasma concentration of 100 ng/ml using a computerized pump (Graseby 3500, Graseby Medical, UK).

The infusion continued for 15 min to allow stabilization of plasma levels. Blood samples were drawn before and after the resting fMRI scan and then placed on ice. Plasma was obtained by centrifugation and stored at −70 °C. Plasma ketamine concentrations were measured by gas chromatography–mass spectrometry.

### Ketamine study: MRI Data Acquisition

All MRI and assessment procedures were identical across assessment occasions. Scanning was performed using a 3.0 T MRI scanner (Siemens Magnetom, Trio Tim, Erlangen, Germany) equipped with a 12-channel array coil located at the Wolfson Brain Imaging Centre, Addenbrooke’s Hospital, Cambridge, UK. T2*- weighted echo-planar images were acquired under eyes-closed resting-state conditions. Participants were instructed to close their eyes and let the minds wander without going to sleep. Subsequent participant debriefing ensured that no participants fell asleep during the scan. Imaging parameters were: 3x3x3.75mm voxel size, with a time-to-repetition (TR) of 2000 ms, time-to-echo (TE) of 30 ms, flip angle of 781 in 64x64 matrix size, and 240mm field of view (FOV). A total of 300 volumes comprising 32 slices each were obtained. In addition, high- resolution anatomical T1 images were acquired using a three-dimensional magnetic- prepared rapid gradient echo (MPPRAGE) sequence. In all, 176 contiguous sagittal slices of 1.0mm thickness using a TR of 2300 ms, TE of 2.98 ms, flip angle of 91, and a FOV of 256mm in 240x256 matrix were acquired with a voxel size of 1.0mm^3^. One participant was excluded due to excessive movement, resulting in a final sample of N=20 subjects.

### LSD Dataset

The original study (Carhart-Harris et al., 2016) was approved by the National Research Ethics Service Committee London–West London and was conducted in accordance with the revised declaration of Helsinki (2000), the International Committee on Harmonization Good Clinical Practice guidelines and National Health Service Research Governance Framework. Imperial College London sponsored the research, which was conducted under a Home Office license for research with schedule 1 drugs. All participants were recruited via word of mouth and provided written informed consent to participate after study briefing and screening for physical and mental health. The screening for physical health included electrocardiogram (ECG), routine blood tests, and urine test for recent drug use and pregnancy. A psychiatric interview was conducted and participants provided full disclosure of their drug use history. Key exclusion criteria included: < 21 years of age, personal history of diagnosed psychiatric illness, immediate family history of a psychotic disorder, an absence of previous experience with a classic psychedelic drug (e.g. LSD, mescaline, psilocybin/magic mushrooms or DMT/ayahuasca), any psychedelic drug use within 6 weeks of the first scanning day, pregnancy, problematic alcohol use (i.e. > 40 units consumed per week), or a medically significant condition rendering the volunteer unsuitable for the study. Twenty healthy volunteers with a previous experience using psychedelic drugs were scanned. Volunteers underwent two scans, 14 days apart. On one day they were given a placebo (10-mL saline) and the other they were given an active dose of LSD (75 μg of LSD in 10-mL saline). The order of the conditions was balanced across participants, and participants were blind to this order but the researchers were not. Participants carried out VAS-style ratings via button-press and a digital display screen presented after each scan, and the 11-factor altered states of conscious- ness (ASC) questionnaire was completed at the end of each dosing day (Carhart-Harris et al., 2016). All participants reported marked alterations of consciousness under LSD.

### LSD study: Infusion Protocol

The data acquisition protocols were described in detail in a previous paper (Carhart-Harris et al., 2016), so we will describe them in brief here. The infusion (drug/placebo) was administered over 2 min and occurred 115min before the resting-state scans were initiated. After infusion, subjects had a brief acclimation period in a mock MRI scanner to prepare them for the experience of being in the real machine. ASL and BOLD scanning consisted of three seven-minute eyes closed resting state scans. The ASL data were not analysed for this study, and will not be discussed further.

### LSD study: MRI Data Acquisition

The first and third scans were eyes-closed, resting state without stimulation, while the second scan involved listening to music; however, this scan was not used in this analysis. The precise length of each of the two BOLD scans included here was 7:20 minutes. For the present analysis, these two scans were concatenated together in time. Imaging was performed on a 3T GE HDx system. High-resolution anatomical images were acquired with 3D fast spoiled gradient echo scans in an axial orientation, with field of view = 256x256x192 and matrix = 256x256x129 to yield 1mm isotropic voxel resolution. TR/TE = 7.9/3.0ms; inversion time = 450ms; flip angle = 20. BOLD-weighted fMRI data were acquired using a gradient echo planer imaging sequence, TR/TE = 2000/35ms, FoV = 220mm, 64x64 acquisition matrix, parallel acceleration factor = 2, 90 flip angle. Thirty five oblique axial slices were acquired in an interleaved fashion, each 3.4mm thick with zero slice gap (3.4mm isotropic voxels). One subject aborted the experiment due to anxiety and four others were excluded for excessive motion (measured in terms of frame-wise displacement), leaving 15 subjects for analysis (11 males, 4 females; mean age 30.5 years, SD = 8.0 years) (Carhart-Harris et al., 2016).

### Test-Retest Dataset

Right-handed healthy participants (N=22, age range, 19–57 years; mean age, 35.0 years; SD 11.2; female-to-male ratio, 9/13) were recruited via advertisements in the Cambridge area and were paid for their participation. Cambridgeshire 2 Research Ethics Committee approved the study (LREC 08/H0308/246) and all volunteers gave written informed consent before participating. Exclusion criteria included National Adult Reading Test (NART) <70, Mini Mental State Examination (MMSE) <23, left- handedness, history of drug/alcohol abuse, history of psychiatric or neurological disorders, contraindications for MRI scanning, medication that may affect cognitive performance or prescribed for depression, and any physical handicap that could prevent the completion of testing. The study consisted of two scanning visits (separated by 2–4 weeks).

### Test-retest dataset: MRI Data Acquisition

For each visit, resting-state fMRI was acquired for 5:20 minutes using a Siemens Trio 3T scanner (Erlangen, Germany). Functional imaging data were acquired using an echoplanar imaging (EPI) sequence with parameters TR 2,000 ms, TE 30 ms, Flip Angle 78◦, FOV 192 × 192mm2, in-plane resolution 3.0 × 3.0mm, 32 slices 3.0mm thick with a gap of 0.75mm between slices. A 3D high resolution MPRAGE structural image was also acquired, with the following parameters: TR 2,300 ms, TE 2.98 ms, Flip Angle 9◦, FOV 256 × 256 mm^2^. Task-based data were also collected, and have been analysed before to investigate separate experimental questions (Manktelow et al., 2017; Moreno-López et al., 2017; Vatansever et al., 2015). A final set of 18 participants had usable data for both resting-state fMRI scans and were included in the present analysis.

### FMRI Preprocessing and denoising

We preprocessed the functional imaging data using a standard pipeline, implemented within the SPM12-based (http://www.fil.ion.ucl.ac.uk/spm) toolbox CONN (http://www.nitrc.org/projects/conn), version 17f (Whitfield-Gabrieli and Nieto-Castanon, 2012). The pipeline comprised the following steps: removal of the first five scans, to allow magnetisation to reach steady state; functional realignment and motion correction; slice-timing correction to account for differences in time of acquisition between slices; identification of outlier scans for subsequent regression by means of the quality assurance/artifact rejection software *art* (http://www.nitrc.org/projects/artifact_detect); structure-function coregistration using each volunteer’s high-resolution T1-weighted image; spatial normalisation to Montreal Neurological Institute (MNI-152) standard space with 2mm isotropic resampling resolution, using the segmented grey matter image, together with an *a priori* grey matter template.

To reduce noise due to cardiac and motion artifacts, we applied the anatomical CompCor method of denoising the functional data (Behzadi Y et al., 2007), also implemented within the CONN toolbox. The anatomical CompCor method involves regressing out of the functional data the following confounding effects: the first five principal components attributable to each individual’s white matter signal, and the first five components attributable to individual cerebrospinal fluid (CSF) signal; six subject-specific realignment parameters (three translations and three rotations) as well as their first- order temporal derivatives; the artifacts identified by *art;* and main effect of scanning condition (Behzadi Y et al., 2007). Linear detrending was also applied, and the subject-specific denoised BOLD signal timeseries were band- pass filtered to eliminate both low-frequency drift effects and high-frequency noise, thus retaining temporal frequencies between 0.008 and 0.09 Hz. Importantly, note that this bandpass filtering pertains to *temporal* frequencies, which are distinct from the *spatial* frequencies obtained from connectome harmonic decomposition (as described below).

Due to the presence of deformations caused by brain injury, rather than relying on automated pipelines, DOC patients’ brains were individually preprocessed using SPM12, with visual inspections after each step. Additionally, to further reduce potential movement artifacts, data underwent despiking with a hyperbolic tangent squashing function. The remaining preprocessing and denoising steps were the same as described above for the ketamine and propofol data.

Finally, the preprocessed and denoised functional MRI data were then projected onto the cortical surface and decomposed in terms of connectome harmonics, as described above.

### Statistical Analysis

Linear Mixed Effects models (implemented as the MATLAB function *fitlme*) were used to assess the statistical significance of the differences between conditions (states of consciousness), treating condition as a fixed effect, and subjects as random effects. When one measurement was obtained for each timepoint, timepoints were also included as random effects, nested within subjects. Results are reported in terms of the fixed effect of condition, and the upper and lower bounds of its 95% confidence interval, with associated p-value. For comparison of frequency-specific harmonic energy, the False Discovery Rate for multiple comparisons across 15 frequency bins was controlled by means of the Benjamini- Hochberg procedure (Benjamini and Hochberg, 1995). Correlations were assessed using Spearman’s non-parametric rank-based ρ. All analyses were two- sided, and statistical significance was assessed at the standard alpha threshold of 0.05.

## Results

Here, we adopt the mathematical framework of connectome harmonic decomposition to undertake an empirical investigation of the similarities and differences between perturbations of consciousness induced by the anaesthetic propofol, severe brain injury, sub-anaesthetic ketamine and LSD.

### Connectome harmonic decomposition: relating brain structure and function to characterise states of consciousness

To map the landscape of consciousness, we decompose brain activity (BOLD signals from functional MRI) during each state of consciousness in terms of multi- scale contributions from the harmonic modes of a representative human structural connectome. These harmonic modes are obtained from eigen-decomposition of the graph Laplacian applied to a high-resolution reconstruction of a representative human connectome, and used as a new set of basis functions to re-represent functional brain signals into whole-brain patterns of different spatial scale (frequency): from an entire hemisphere to just a few millimetres (Atasoy et al., 2018a).

In traditional Fourier analysis, the temporal frequency of each temporal harmonic (sinusoid) reflects how much the signal varies over time (Figure 1A,B); likewise, CHD quantifies to what extent the BOLD signal is constrained by the global network organisation of the connectome, or deviates from it. Low-frequency connectome harmonics correspond to coarse-grained patterns of spatial variation, whereby structurally connected nodes have similar values of the functional signal; in contrast, high-frequency connectome harmonics denote fine-grained patterns of spatial variation, such that nodes can have different values of the functional signal irrespective of whether they are structurally connected (Figure 1C,D). An overview of our analytic workflow is provided in Figure 2.

**Figure 2.**
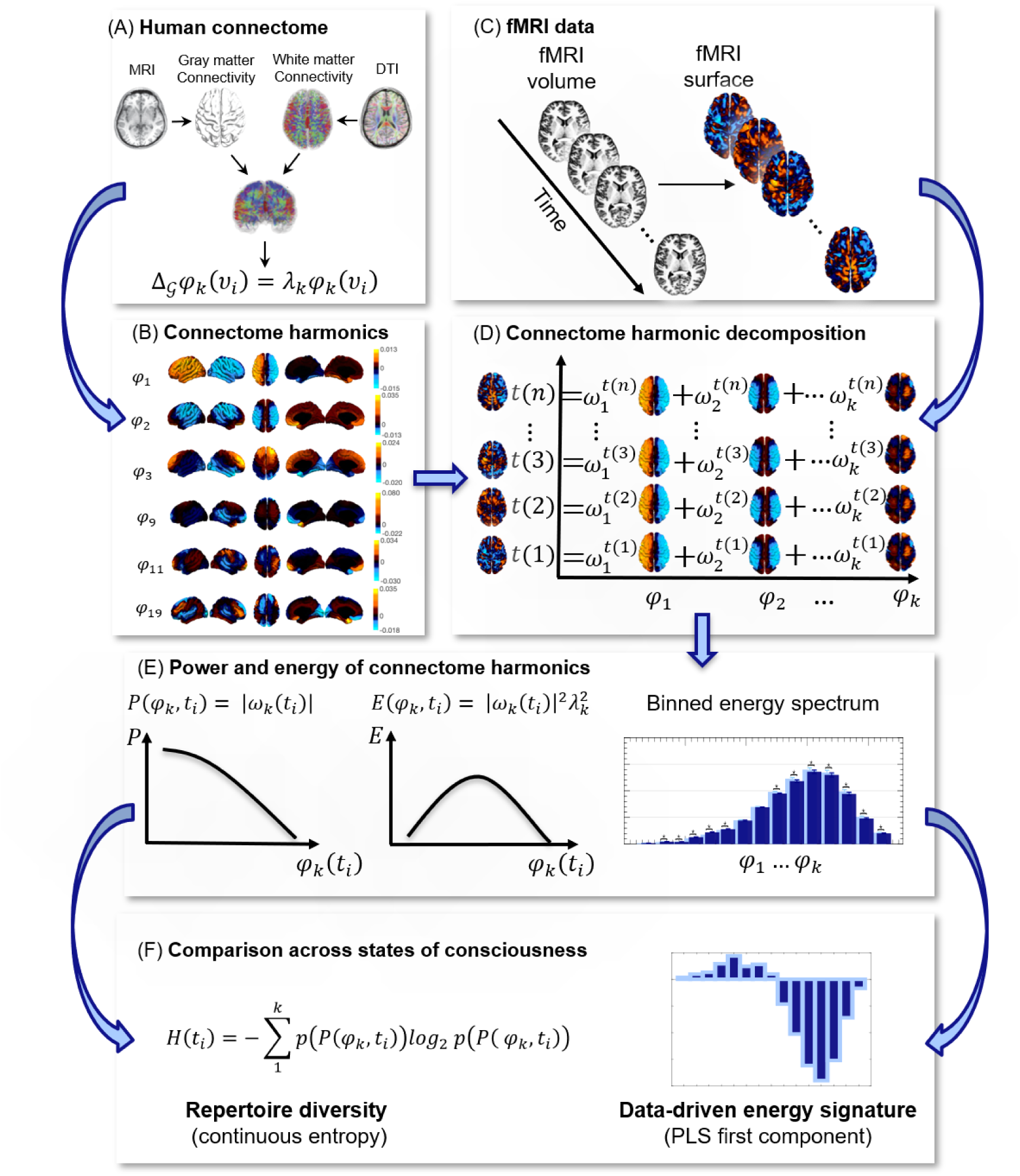
Connectome harmonic decomposition: relating brain structure and function to characterise states of consciousness. (A) High-resolution rendering of the representative human connectome is obtained from HCP subjects by combining the surface-based local connections within the grey matter, reconstructed from structural magnetic resonance imaging (sMRI); and the long-range white-matter axonal tracts calculated with diffusion tensor imaging (DTI), thereby taking into account both local and long-range connectivity. (B) The graph Laplacian of this high-resolution connectome is then decomposed into its eigenvectors *φ*_1…*n*_ (harmonic modes) and their associated eigenvalues *λ*_1…*n*_ (spatial frequencies with increasing granularity). With an increasing connectome harmonic number k, we obtain more complex and fine-grained spatial patterns. (C) For every timepoint *ti*, functional magnetic resonance imaging (fMRI) data are projected from volumetric space onto the cortical surface. (D) Connectome harmonic decomposition (CHD) of the fMRI data estimates the contribution ω_k_(t_i_) of each harmonic mode φ_k_ to the cortical activity at every timepoint *ti*. (E) The connectome harmonic power spectrum is estimated as the absolute magnitude of contribution of each individual harmonic φ_k_ to the fMRI data at every time point *ti*: *P*(*φ*_*k*_, *t_i_*) = |*ω*_*k*_(*t_i_*)|. Similarly, the connectome harmonic energy spectrum is estimated as the square of the absolute contribution |ω_k_(t_i_)| of individual harmonics φ_k_ to the fMRI data, weighted by the square of the harmonics’ corresponding eigenvalue λ_k_ (intrinsic energy) at every time point *t_i_*: *E*(*φ*_*k*_, *t_i_*) = |*ω*_*k*_(*t_i_*)|^2^*λ*_k_^2^). The overall binned energy spectrum across subjects and time points is constructed by discretising the energy of connectome harmonics in 15 logarithmically-spaced frequency-specific bins, here shown for a target state (dark blue) and a reference state (light blue), following previous work showing that this procedure can successfully highlight the connectome harmonic signatures of altered states of consciousness (Atasoy et al., 2017, 2018b) (Figure S1). (F) Repertoire entropy is defined as the entropy of the power spectrum across all 18,715 harmonics, for every timepoint *ti*, computed with the continuous Kozachenko approximation; the data-driven energy signature of a target state of consciousness is obtained from the first principal component of Partial Least Squares-Discriminant Analysis (PLS-DA), which maximally discriminates the target state (dark blue) from the reference state (light blue), based on their respective binned connectome harmonic energy spectra.

We begin by demonstrating the test-retest reliability of CHD. In a test-retest fMRI dataset of 18 individuals, each scanned twice during resting wakefulness within a timespan of 2-4 weeks, we show that no discernible pattern of differences can be identified when comparing the energy spectra of the first and second scans (Figure S2). This evidence indicates that connectome harmonic patterns remain stable across scans of the same individuals, when they are in the same state of consciousness (resting wakefulness) – providing an important “negative control” for our subsequent results.

### Loss of consciousness and the psychedelic state are characterized by specific and opposite connectome harmonic signatures

Based on computational modelling, we had previously predicted that increased global inhibition should lead to a shift in the frequency-specific contribution (energy) of connectome harmonics: from high-frequency (fine-grained), structurally decoupled harmonic patterns to low-frequency (coarse-grained), structurally coupled ones (Atasoy et al., 2016, 2018a). Here, we began by testing this prediction: as an agonist of the chief inhibitory neurotransmitter GABA, propofol induces globally increased neuronal inhibition (Brown et al., 2010). In accordance with our hypothesis, across N=15 volunteers undergoing increasing levels of propofol sedation, we observed significantly increased energy of low-frequency harmonics and significantly decreased energy of high-frequency harmonics (Figure 3A-E, Figure S3 and Table S1), reflecting greater dependence of the signal on the network structure of the human connectome.

**Figure 3.**
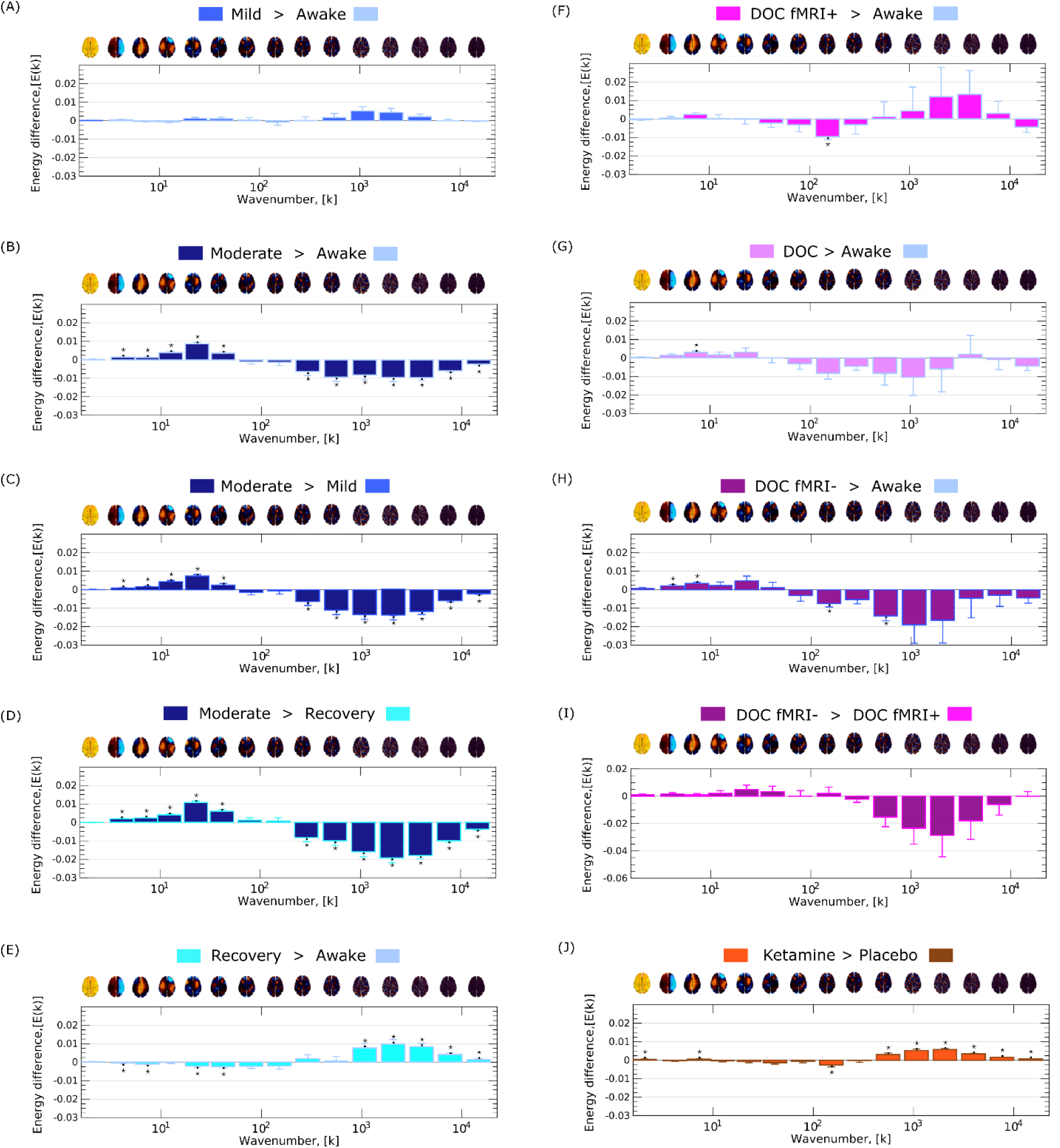
Loss of consciousness and the psychedelic state are characterized by specific and opposite signatures of connectome harmonic energy. (A) Mild propofol sedation > wakefulness. (B) Moderate anaesthesia > wakefulness. (C) Moderate anaesthesia > mild sedation. (D) Moderate anaesthesia > post-anaesthetic recovery. (E) Recovery > wakefulness. (F) DOC patients > awake healthy controls. (G) DOC fMRI+ patients > awake healthy controls. (H) DOC fMRI- patients > awake healthy controls. (I) fMRI- > fMRI+ DOC patients. (J) Ketamine > placebo. * *p* < 0.05, FDR-corrected across 15 frequency bins. A brain surface projection of the connectome harmonic pattern corresponding to each frequency bin, averaged over the constituent spatial frequencies, is shown above each bin. See Figure S4 for our re-derivation of the LSD signature using the same preprocessing and denoising procedures as for our other data to ensure consistency.

Note that this characteristic connectome harmonic signature is not found at just any level of propofol: it is not present for mild sedation, when participants are drowsy but still conscious (Figure 3A). Instead, it only arises for a dose of propofol sufficient to induce loss of responsiveness (Figure 3B,C), and it is then reversed upon post-anaesthetic recovery of responsiveness (Figure 3D) (note that here we follow the relevant literature in considering individuals to be unconscious when they fail to provide evidence of consciousness through either overt or covert responsiveness; we return to this issue in the Discussion). Therefore, the appearance of this connectome harmonic signature is tied to propofol-induced loss of responsiveness.

Crucially, an analogous connectome harmonic signature is also observed for chronic loss of responsiveness induced by severe brain injury – but only when restricting the comparison to patients who provided no evidence of being conscious (Figure 3F-I). Specifically, we studied a cohort of N=22 patients: although all met diagnostic criteria for disorders of consciousness based on overt behaviour, eight patients nevertheless provided evidence of covert consciousness by successfully performing mental imagery tasks in the fMRI scanner (labelled fMRI+), whereas the remaining 14 did not (labelled fMRI-) (Craig et al., 2021; Luppi et al., 2021c).

When comparing the entire cohort of DOC patients against awake healthy controls, connectome harmonic decomposition revealed an energy signature with significant similarity to the signature of moderate propofol anaesthesia (Spearman’s *ρ* = 0.60, CI_95%_ [0.13, 0.85], *p* = 0.020), although most frequency-specific differences did not reach statistical significance after correction for multiple comparisons (Figure 3G). Crucially, when the comparison was restricted to fMRI- patients versus controls, the overall signature increased its similarity the connectome harmonic signature of moderate propofol anaesthesia, both visually and numerically (Spearman’s *ρ* = 0.87, CI_95%_ [0.64, 0.96], *p* < 0.001; Figure 3H). Indeed, in this case we observed both significant increases in the energy of some low-frequency connectome harmonics, and also significant decreases in the energy of some high-frequency harmonics - despite using a smaller sample of patients (Figure 3H).

Remarkably, this overall harmonic signature also persisted when comparing fMRI- patients against fMRI+ patients, despite the fact that both groups of patients are diagnosed as suffering from disorders of consciousness based on their overt behaviour (Figure 3I). Although individual frequency-specific differences did not reach statistical significance after correction for multiple comparisons, the overall signature remained strongly and significantly correlated with the signature of moderate propofol anaesthesia (Spearman’s *ρ* = 0.94, CI_95%_ [0.81, 0.98], *p* < 0.001). Crucially, however, this putative unconsciousness-specific pattern of connectome harmonic energy was *not* observed when comparing the subgroup of fMRI+ DOC patients with awake volunteers (Figure 3F). This is reassuring, being what we should expect from a specific marker of unconsciousness, given that each fMRI+ patient had previously exhibited evidence of being covertly conscious.

Having investigated connectome harmonic signatures across different ways of losing consciousness, we next sought to further expand our investigation of human consciousness by considering the altered state induced by a sub-anaesthetic dose of the NMDA receptor antagonist, ketamine, and comparing it with the previously published connectome harmonic signatures of classic serotonergic psychedelics (Atasoy et al., 2017, 2018b). At sub-anaesthetic doses, ketamine induces an altered state of consciousness with psychedelic-like subjective experiences, including perceptual distortions, vivid imagery and hallucinations, and dissociative symptoms (Corlett et al., 2016; Dandash et al., 2015; Moore et al., 2011).

Connectome harmonic decomposition of fMRI data from N=20 volunteers revealed a significant increase in total brain energy during infusion with a sub-anaesthetic dose of ketamine, compared with placebo (Table S1 and Figure S3J). Sub-anaesthetic ketamine increased the energy of high-frequency harmonics just like LSD and psilocybin (Atasoy et al., 2017, 2018b) (Figure 3J); despite lacking LSD’s pronounced suppression of the low-frequency harmonics, the energy signature of ketamine was strongly correlated with the signature of LSD (Spearman’s *ρ* = 0.98, CI_95%_ [0.95, 0.99], *p* < 0.001). Validating theoretical predictions (Atasoy et al., 2018a), our findings demonstrate that the common psychoactive effects of ketamine and classic serotonergic psychedelics are reflected in their common increases of high-frequency connectome harmonics, despite occurring through different molecular mechanisms: increased global excitation arising from NMDA receptor antagonism and 5HT_2A_ receptor agonism, respectively. Thus, CHD can also identify similar alterations in consciousness induced by different pharmacological interventions.

Intriguingly, elevated energy in high-frequency connectome harmonics, and reduced energy in low-frequency ones, were also observed when comparing post-anaesthetic recovery and pre-anaesthetic wakefulness – resembling the pattern previously observed with classic psychedelics (Atasoy et al., 2017, 2018b). We elaborate on possible interpretations of this observation in the Discussion. We also show (Figure S5) that the connectome harmonic energy signatures are preserved if a different number of bins (25 instead of 15) is used (note that binning is logarithmic). Additionally, no significant correlations are found between the connectome harmonic signatures of different states of consciousness (Figure 3) and the signature obtained from test-retest scans of the same awake individuals (Figure S2) (Table S2).

### Connectome harmonic signatures generalise across states of consciousness

It is readily apparent from Figure 3 that anaesthesia and disorders of consciousness are characterised by similar patterns of change across the spectrum of connectome harmonic energy; in addition, this signature looks like a mirror-reversed version of the change in energy corresponding to the psychedelic state (whether induced by sub-anaesthetic ketamine or serotonergic psychedelics (Atasoy et al., 2017, 2018b) (Figures 3J, S4). These observations suggest that it should be possible to generalise connectome harmonic patterns across datasets, to establish the harmonic signature of a) unconsciousness and b) the psychedelic state.

To take into account the full spectrum of connectome harmonic changes at the same time, we turned to Partial Least Squares Discriminant Analysis (PLS-DA) (Krishnan et al., 2011): this data-driven technique allowed us to extract the multivariate patterns of connectome harmonic energy that maximally distinguish between each pair of conditions (termed “multivariate signatures”, MVS). This approach clearly revealed the existence of two mirror-reversed multivariate patterns characterising loss of consciousness and the psychedelic state (Figure 4). We focused on four key multivariate signatures: two intended to represent loss of consciousness (awake vs moderate propofol, Figure 4A; and DOC fMRI+ vs fMRI-, Figure 4I); and two intended to reflect the psychedelic state (sub-anaesthetic ketamine vs placebo, Figure 4J; and LSD vs placebo, using the LSD data previously analysed by Atasoy and colleagues (Atasoy et al., 2017, 2018b); Figure 4K).

**Figure 4.**
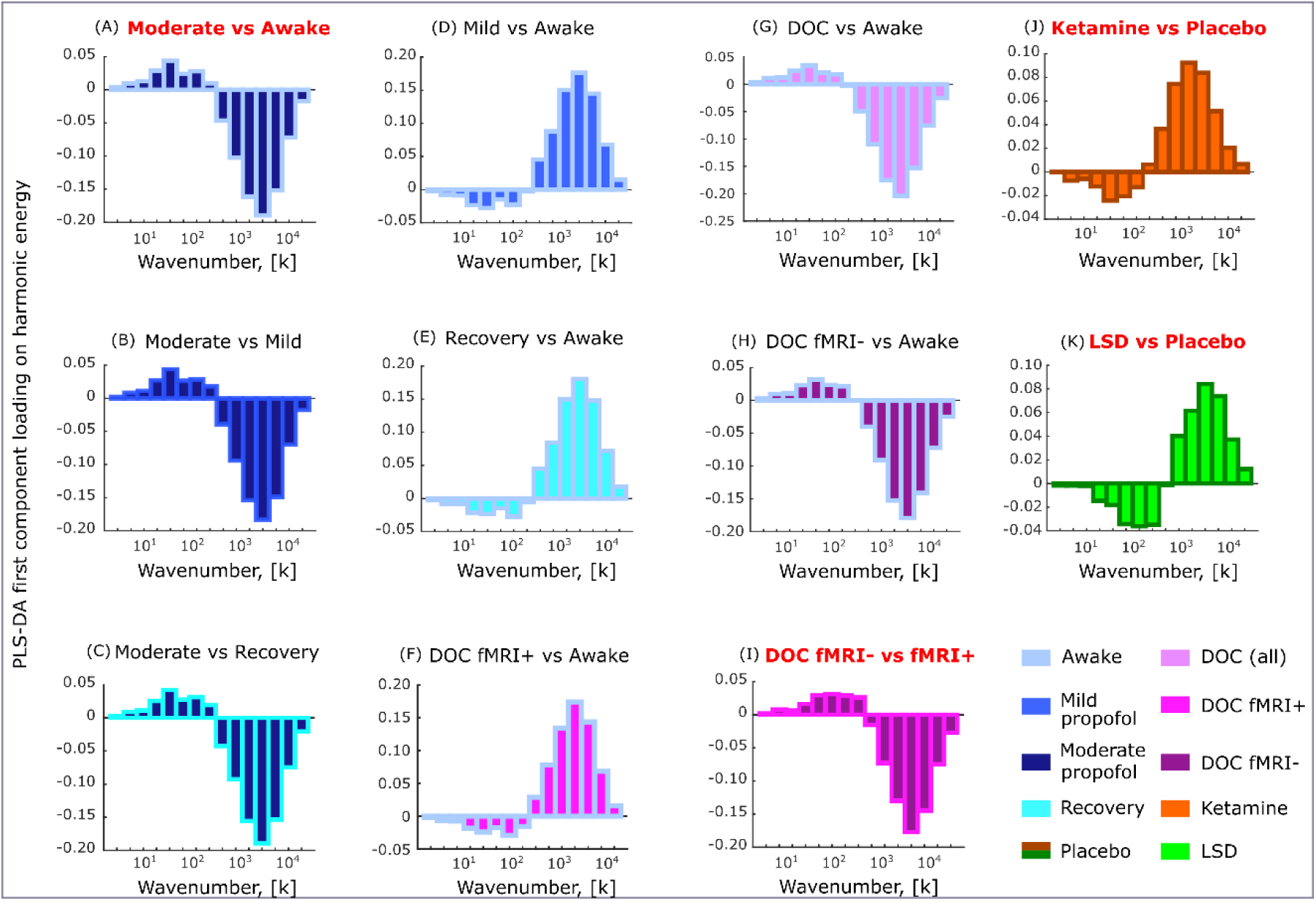
Similar and opposite multivariate signatures of connectome harmonic energy for different states of consciousness. (A) Moderate anaesthesia > wakefulness. (B) Moderate anaesthesia > mild sedation. (C) Moderate anaesthesia > post-anaesthetic recovery. (D) Mild sedation > wakefulness. (E) Post-anaesthetic recovery > wakefulness. (F) DOC fMRI+ patients > awake healthy controls. (G) DOC patients > awake healthy controls. (H) DOC fMRI- patients > awake healthy controls. (I) fMRI- > fMRI+ DOC patients. (J) Ketamine > placebo. (K) LSD > placebo. Bar colour indicates the target state; contours indicate the reference state. Titles in bold red font indicate the four contrasts used for the analysis described in Figure 5. Virtually identical patterns are obtained if using 25 logarithmically spaced bins instead of 15 (Figure S6).

For each of these four multivariate signatures, we asked whether it could be generalised to other states of consciousness, beyond the one it was derived from. We pursued this hypothesis by projecting each subject’s connectome harmonic energy spectrum onto a given MVS (thereby measuring the correspondence between them), and then comparing the value of this projection across groups, to test whether it could be used to identify a statistically significant difference between them.

This analysis indicated that datasets with a positive alignment with the propofol MVS also aligned positively with the DOC MVS, and aligned negatively with both the ketamine and LSD MVS – and vice-versa (Figure 5 and Table S3). In particular, whenever a statistically significant difference was observed in terms of the propofol projection, a significant difference in the same direction was also detected in terms of the DOC projection, and corresponding differences in the opposite direction were observed for the ketamine and LSD projections (Figure 5). Therefore, these results support the notion that two opposite energy patterns across the connectome harmonic spectrum characterise loss of consciousness and the psychedelic state, respectively. Note that nothing mandates that these should be the only two patterns observed; indeed, when considering our test-retest data, the MVS does not resemble either of the two main patterns that we observed for alterations of consciousness (Figure S2).

**Figure 5.**
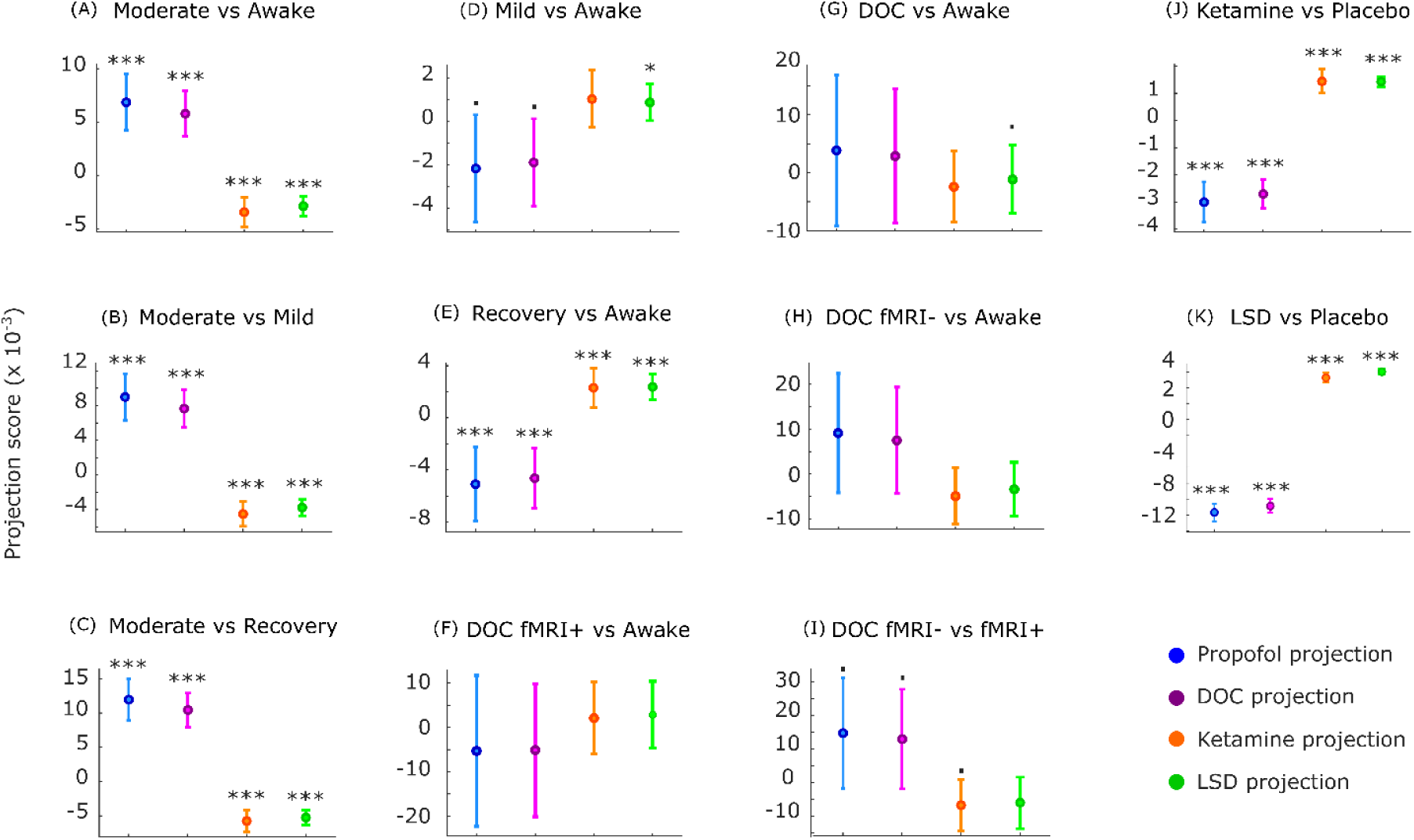
Multivariate signatures of connectome harmonic energy are generalisable across states of consciousness. Each panel shows the fixed effects (and 95% CI) of contrasting the projections (dot product) of MVS patterns from four source states (moderate propofol anaesthesia, DOC patients, ketamine and LSD; Figure 3) onto each state of consciousness under comparison. (A) Moderate anaesthesia > wakefulness. (B) Moderate anaesthesia > mild sedation. (C) Moderate anaesthesia > post-anaesthetic recovery. (D) Mild sedation > wakefulness. (E) Post-anaesthetic recovery > wakefulness. (F) DOC fMRI+ patients > awake healthy controls. (G) DOC patients > awake healthy controls. (H) DOC fMRI- patients > awake healthy controls. (I) fMRI- > fMRI+ DOC patients. (J) Ketamine > placebo. (K) LSD > placebo. * *p* < 0.05; ** *p* < 0.01; *** *p* < 0.001;. *p* < 010.

### Connectome harmonic signatures are related to pharmacological and subjective scores

Next, we aimed to provide an even more compelling demonstration that the signatures of unconsciousness extracted from CHD are generalisable across ways of losing consciousness, by relating them to the underlying neurobiology across subjects. For each stage of sedation in the propofol dataset (mild, moderate, and recovery) we projected the connectome harmonic energy of each subject onto the connectome harmonic signature of unconsciousness (designated as the MVS that best discriminates between fMRI+ and fMRI- DOC patients; Figure 4I), to quantify their alignment.

Extending the results of the previous analysis, we show that changes in this alignment across progressive stages of sedation correlate with changes in propofol concentration in the blood serum. We observed this both for the transition from consciousness to unconsciousness, i.e. from mild to moderate propofol anaesthesia (Spearman’s *ρ* = 0.57, CI_95%_ [0.08, 0.84], *p* = 0.026; Figure 6A), and also when transitioning back from moderate anaesthesia to recovery (Spearman’s *ρ* = 0.56, CI_95%_ [0.07, 0.83], *p* = 0.030; Figure 6B). In other words, a greater increase in propofol concentration when transitioning from consciousness to unconsciousness, corresponds to a greater neural alignment with the connectome harmonic signature of unconsciousness (extracted from DOC patients) – and vice-versa when awakening from anaesthesia. These results establish both the generalisability of this connectome harmonic signature of unconsciousness, and also its biological relevance, by correlating with a key pharmacological measure that we know to be causally related to the induction of unconsciousness (propofol concentration).

**Figure 6.**
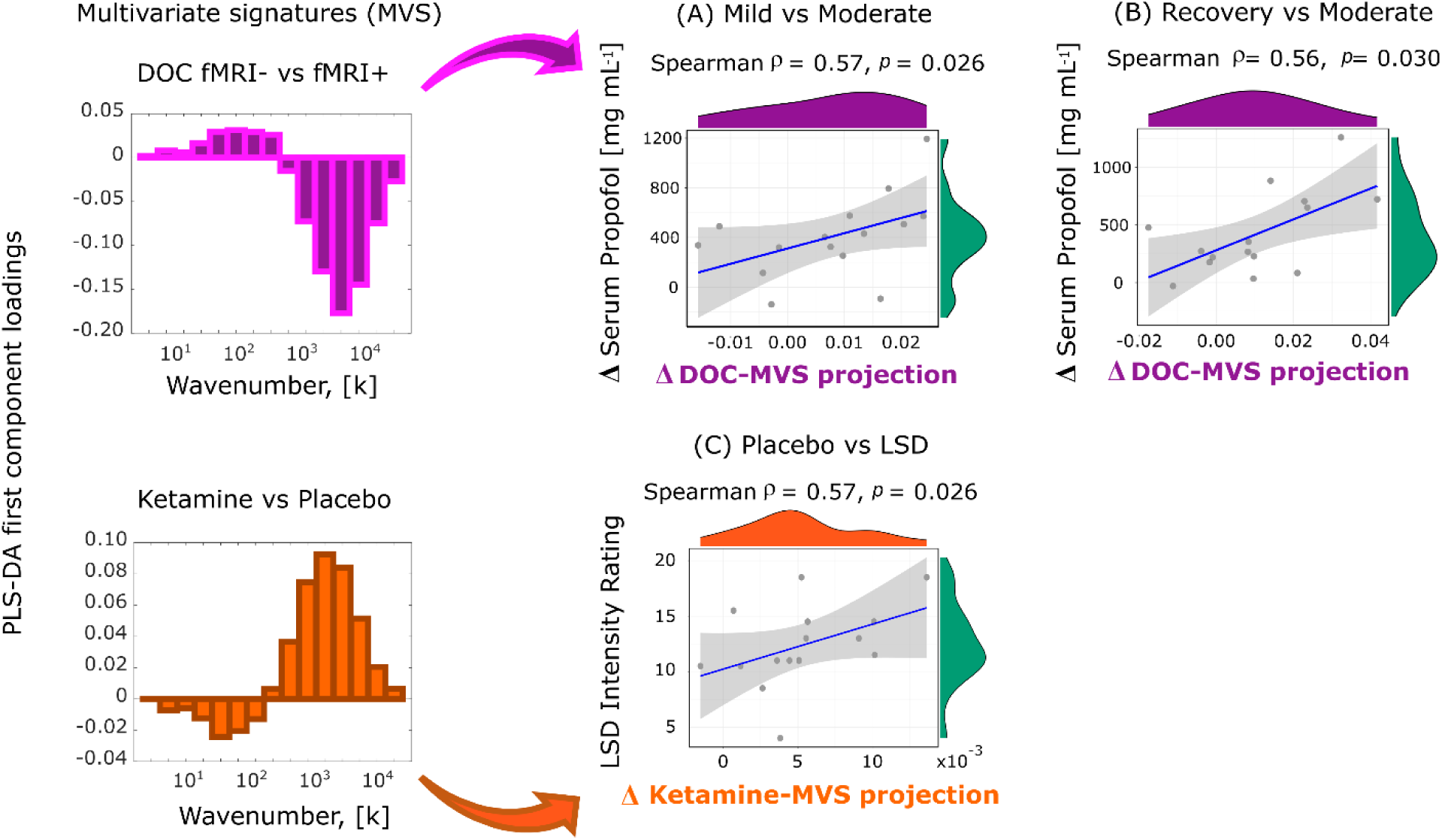
Neurobiological and subjective relevance of connectome harmonic signatures across states of consciousness. Each scatterplot displays individual scores (serum propofol change or subjective intensity) versus the drug-induced change in projection (alignment) of that subject’s spectrum of connectome harmonic energy, onto the multivariate energy signature (MVS) displayed on the left. (A) Change in energy projection onto the DOC energy signature between mild and moderate anaesthesia (moderate minus mild), versus the change of propofol levels in volunteers’ blood serum. (B) Change in energy projection onto the DOC energy signature between moderate anaesthesia and recovery (moderate minus recovery), versus the change of propofol levels in volunteers’ blood serum. (C) Change in energy projection onto the ketamine energy signature between placebo and LSD (LSD minus placebo), versus the subjective intensity of the psychedelic experience induced by LSD. Note that this similarity between similar subjective states of consciousness induced by different means is *not* brought about by generic confounding effects across the datasets, such as head motion: for both the propofol and LSD datasets, the same MVS projections were not significantly correlated with differences in head motion, thereby excluding this potential confound (Figure S7).

Furthermore, we demonstrate that the generalisability of connectome harmonic signatures also extends to the psychedelic state (note that from here on we use the term “psychedelic” to refer to the phenomenology that is shared by classic serotonergic psychedelics and sub-anaesthetic ketamine, in line with previous work (Li and Mashour, 2019; Schartner et al., 2017a)). Using the LSD data previously used by Atasoy and colleagues (Atasoy et al., 2017, 2018b), we show that for each individual, the subjective intensity of the psychedelic experience induced by LSD can be predicted by the change in alignment between the subjects’ energy spectrum and the connectome harmonic signature derived from ketamine (Spearman’s *ρ* = 0.57, CI_95%_ [0.08, 0.84], *p* = 0.026; Figure 6C). In other words, the more a subject’s energy pattern becomes similar to the connectome harmonic signature of the psychedelic state (as extracted from the ketamine dataset), the more intense that subject will rate the subjective experience induced by LSD. These results suggest a profound connection between the neural and phenomenological aspects of the psychedelic state, regardless of how induced.

Overall, these findings identify connectome harmonic signatures of conscious-to- unconscious transitions (and vice-versa) induced by both propofol anaesthesia and DOC, as well as signatures of the psychedelic experience induced by the serotonergic psychedelic LSD and the “atypical” psychedelic ketamine. In addition, we have shown that connectome harmonic signatures can relate brain activity to both pharmacology (correlating with the change in propofol in the bloodstream) and subjective phenomenology (correlating with the intensity of the psychedelic experience induced by LSD).

### Diversity of connectome harmonic repertoire tracks level of consciousness from loss of responsiveness to psychedelics

Having demonstrated that connectome harmonic signatures can generalise between specific states of consciousness (different ways of losing consciousness, or different psychedelic drugs), we sought to provide one further level of generalisation, by explicitly bringing all the states of consciousness considered here into the same continuum.

Specifically, recent theoretical efforts seeking to establish a correspondence between the dynamics of mind and brain (Carhart-Harris et al., 2014; Northoff et al., 2020) posit that states of diminished consciousness should be characterized by a more restricted repertoire of brain patterns – whereas the rich mental content and diversity of experiences that characterize the psychedelic state (Carhart- Harris and Friston, 2019; Carhart-Harris et al., 2014) should correspond to an expanded repertoire of brain patterns. Here, we pursued this hypothesis by quantifying the diversity of the connectome harmonics that are recruited to compose brain activity, across different states of consciousness. Specifically, we expected that anaesthesia and disorders of consciousness should exhibit reduced diversity (quantified in terms of entropy) of connectome harmonics, whereas increased entropy should be observed for ketamine and LSD.

Our results support each of these predictions (Figure 7 and Table S4). Ketamine and LSD exhibited significantly higher diversity of the repertoire of connectome harmonics than placebo, whereas moderate anaesthesia with propofol induced a collapse in the repertoire when compared with wakefulness, recovery, and even mild sedation. Remarkably, our analysis revealed that DOC patients who had previously exhibited evidence of covert consciousness (fMRI+), also exhibited entropy levels comparable to those of awake volunteers – in sharp contrast with fMRI- patients, who had provided no evidence of being conscious, and for whom the repertoire entropy of connectome harmonic signatures was significantly compromised (Figure 7 and Table S4). We also repeated this analysis with a different stratification of DOC patients, combining clinical diagnosis and fMRI- based assessment: on one hand, we grouped together covertly conscious patients (fMRI+) and patients diagnosed with a less severe disorder of consciousness based on their overt behaviour (minimally conscious state) (N=14); and on the other hand were patients (N=8) who were both classified as fMRI- based on lack of in-scanner brain responses, and diagnosed with unresponsive wakefulness syndrome based on lack of overt behaviour (Table 1). The binned harmonic energy signature and PLS-derived multivariate signature for this contrast (Any response vs No response) both showed the expected pattern. Crucially, the two groups also differed significantly in terms of the diversity (entropy) of the full connectome harmonic repertoire, which was significantly diminished for No-response (i.e., fMRI- UWS) patients (Figure S8 and Table S4).

**Figure 7.**
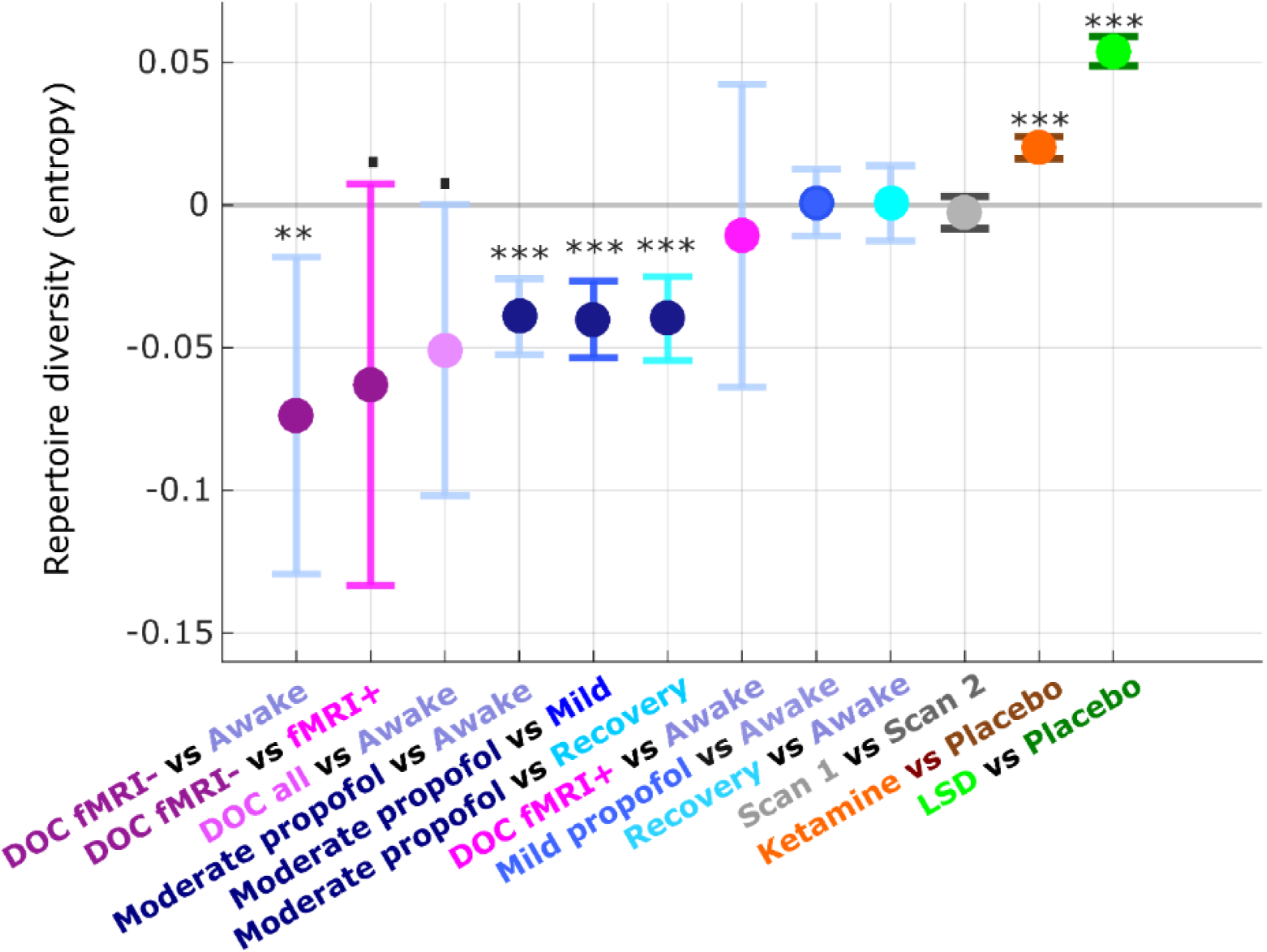
Diversity of connectome harmonic repertoire tracks level of consciousness from loss of responsiveness to psychedelics. At every timepoint *ti*, the contribution of each connectome harmonic to the overall pattern of brain activity is quantified by the harmonic power: *P*(*φ*_*k*_, *t_i_*) = |*ω*_*k*_(*t_i_*)|). Note that no binning was used for this analysis. The diversity of the repertoire of harmonic power can then be quantified in terms of the entropy of the power distribution: the higher the “harmonic repertoire diversity”, the wider the range of connectome harmonics that are recruited to compose cortical activity. Pairs of conditions (states of consciousness) were compared with linear mixed effects modelling, by treating condition as a fixed effect (indicated on the Y axis along with 95% CI) and subjects as random effects. Timepoints were also included as random effects, nested within subjects. *** *p* < 0.001; ** *p <* 0.01; . *p* < 0.10.

Conversely, neither mild sedation nor recovery (during which volunteers were conscious) was significantly different from normal wakefulness in terms of their diversity of harmonic repertoire, despite the presence of propofol in the blood in both cases. This confirms that diversity of the connectome harmonic repertoire is closely associated with the presence or absence of consciousness. As an important validation, no differences in repertoire diversity were observed in our test-retest dataset, when comparing two scans of the same healthy volunteers during normal wakefulness. Taken together, our results demonstrate that the diversity (entropy) of connectome harmonic repertoire can track variations in conscious state on a one-dimensional continuum.

### Role of the human connectome for mapping states of consciousness

Finally, we demonstrate that using the human structural connectome for harmonic mode decomposition of fMRI signals is crucial to obtaining a coherent mapping of states of consciousness. The ability of CHD to recover meaningful patterns among the various states of consciousness considered here is critically compromised, if the neuroanatomical distribution of the connectome harmonics is disrupted through spatial rotation (Alexander-Bloch et al., 2018; Glomb et al., 2021), or if the structural connectome is subjected to degree-preserving randomisation, thereby perturbing its topology (Figures S9-14 and Tables S5-6)).

In contrast, our discoveries are successfully replicated when employing alternative methods of reconstructing a high-resolution representative human connectome – whether from combining a much larger sample of 985 HCP subjects, corresponding to a 100-fold increase in sample size (Figures S15-18, Table S7), or by using high-quality DSI data (Figures S19-22, Table S8), thereby demonstrating the robustness of our approach. In fact, using these state-of-the-art connectome reconstructions highlights additional significant frequency-specific differences between DOC patients and healthy controls (Figures S16H,I S20H,I), as well as a significant difference in repertoire diversity between fMRI+ and fMRI- DOC patients (Figures S19, S22). Taken together, these results indicate that the harmonic modes of the representative human connectome may represent an especially suitable frame of reference for mapping the landscape of consciousness across individuals and datasets.

## Discussion

Here, we set out to address a key challenge of contemporary neuroscience: relating different states of consciousness to the underlying brain states. Leveraging the recent framework of connectome harmonic decomposition (CHD) of functional MRI data (Atasoy et al., 2016, 2018a), we investigated a wide range of perturbations of human consciousness: propofol anaesthesia, disorders of consciousness, and the altered states induced by psychoactive (sub-anaesthetic) doses of ketamine and by the serotonergic psychedelic LSD.

To understand how brain structure supports human consciousness and its alterations, we sought a mathematically principled interpretation of altered states of consciousness in terms of structure-function relationships across scales. Generalizing the Fourier transform to the network structure of the human brain (Atasoy et al., 2016), CHD explicitly re-expresses brain activity in terms of multi- scale contributions from the underlying structural network: each connectome harmonic is a distributed activation pattern characterized by a specific spatial scale (frequency). Just like temporal harmonics from traditional Fourier analysis quantify time-dependence in the signal, so the harmonic modes of the human connectome quantify connectome-dependence in brain activity across scales (Figure 1). Therefore, CHD provides a principled alternative to go beyond the dominant view of brain activity as consisting of discrete spatial locations, offering complementary insights that are not available from the location-centric perspective.

### Connectome harmonics reinterpret human consciousness in terms of structure-function relationships

Complementing extensive previous studies that have sought to implicate specific neuroanatomical regions in supporting consciousness (Boveroux et al., 2010; Hannawi et al., 2015; Huang et al., 2020; Luppi et al., 2019, 2021b; MacDonald et al., 2015; di Perri et al., 2018; Ranft et al., 2016; Spindler et al., 2021; Vanhaudenhuyse et al., 2010; Warnaby et al., 2016), our results reveal that the range of states of consciousness considered here can be characterised by their specific patterns of connectome harmonics, regardless of how they are induced. Despite their different molecular mechanisms of action, ketamine (at psychedelic-like dosage) and the serotonergic psychedelic LSD increase the contribution of high-frequency (fine-grained) connectome harmonics, whereas propofol- and brain injury-induced unconsciousness lead to reduced high-frequency and increased low-frequency connectome harmonics - demonstrating the value of CHD as a neural marker of conscious state across datasets.

Recall that formally, the progression from low- to high-frequency connectome harmonics reflects increasing decoupling of functional brain activity from the underlying structural connectivity (Medaglia et al., 2018; Preti and Van De Ville, 2019) (Figure 1). Therefore, our results can be interpreted as showing that unconsciousness and the psychedelic state stand in opposite relationships with respect to human structural connectivity (as encoded in the representative high-resolution connectome obtained from HCP data). In unconsciousness, our analysis reveals that functional brain activity becomes more constrained by the structure of the human connectome, as indicated by an increased contribution of low-frequency, large-scale harmonic modes and a more restricted repertoire. In contrast, the psychedelic-induced energy shift to high-frequency harmonics indicates a departure from standard activity patterns encoded in the structural connectome, in favour of increasingly diverse and variable ones (Naze et al., 2021) - a plausible neural correlate for the phenomenologically rich state of mind induced by psychedelics (Atasoy et al., 2017, 2018b, 2018a), according to leading theoretical accounts (Carhart-Harris, 2018; Carhart-Harris and Friston, 2019; Carhart-Harris et al., 2014; Sarasso et al., 2021).

CHD overcomes the main limitations of current structure-function investigations of consciousness, which typically ignore the multi-scale network organization of the connectome, as well as the asymmetric relation between structure and function in the brain. However, it is reassuring that the present results about structure-function relationships of human brain *activity* are complementary with previous evidence pertaining to functional *connectivity,* whereby both anaesthesia and disorders of consciousness increase the correlation between the patterns of structural and functional connections (Barttfeld et al., 2015; Demertzi et al., 2019; Tagliazucchi et al., 2016), whereas LSD reduces structure-function correlation (Luppi et al., 2021a). Thus, converging evidence implicates a key role of structure-function relationships to understand human consciousness – supporting the explicitly connectome-oriented view that is offered by the connectome harmonics.

### Connectome harmonics and the temporal dimension

By comparing different states of consciousness in terms of the same spectrum of connectome harmonics (i.e., harmonic modes obtained from the same high-resolution representative human connectome), we have shown that diversity of the connectome harmonic repertoire provides a powerful one-dimensional indicator of level of consciousness, sensitive to differences in anaesthetic dose (mild sedation vs moderate anaesthesia vs recovery) as well as behaviourally indistinguishable sub-categories of patients with disorders of consciousness.

It is important to emphasise that this result is fundamentally distinct from previous evidence of diminished entropy of temporal signals during loss of consciousness (Burioka et al., 2005; Liu et al., 2019; Luppi et al., 2019; Olofsen et al., 2008; Schartner et al., 2015, 2017b; Varley et al., 2020) (but see (Pal et al., 2019)) and increased temporal entropy in the psychedelic state (Carhart-Harris and Friston, 2019; Lebedev et al., 2016; Li and Mashour, 2019; Schartner et al., 2017a; Tagliazucchi et al., 2014) (see (Sarasso et al., 2021) for a recent review). Those previous studies quantified the diversity of brain signals by focusing on the temporal dimension (“Does the brain visit few or many states in a given period of time?”). In contrast, here we quantified diversity in terms of the repertoire of connectome harmonic frequencies that contribute to brain activity (“Do we need a wide or restricted repertoire of connectome harmonics to build the brain activity we observed?”). Therefore, our harmonic-based measure of diversity and the entropy of temporal signals provide complementary rather than redundant perspectives.

Likewise, it is intriguing that loss of consciousness also tends to increase the prevalence of low-frequency (i.e., slow) temporal oscillations, and vice-versa for psychedelics (Bastos et al., 2021; Ching et al., 2010; Colombo et al., 2019; Donoghue et al., 2019; Ní Mhuircheartaigh et al., 2013; Pal et al., 2019; Stephen et al., 2020; Warnaby et al., 2017). However, temporal frequencies and connectome harmonic frequencies are distinct concepts, each providing a unique perspective, and should not be confused or conflated: recent work has begun to investigate the relationship between connectome eigenmodes and M/EEG temporal frequencies (Glomb et al., 2020; Raj et al., 2020; Verma et al., 2022), opening the door for future multi-modal studies combining fMRI and EEG to elucidate the complex inter-relationships between connectome harmonics, temporal frequencies, and consciousness in the human brain.

### Connectome harmonics relate brain structure and function with neurophysiology and phenomenology

Computational modelling work has shown that the relative prevalence of high- vs low-frequency connectome harmonics in brain activity is governed by the global balance between excitation and inhibition (Atasoy et al., 2016, 2018a). Therefore, changes in the connectome harmonic repertoire reflect the influence of both neuroanatomy (as the source of harmonics) and neurophysiology (governing their relative prevalence) on brain function. This suggests that propofol-induced global inhibition may provide an explanation for the increased structure-function coupling observed during anaesthesia, by restricting the repertoire of connectome harmonic “building blocks” that are available to contribute to brain activity. Indeed, our results demonstrate that change in propofol concentration correlates with the alignment between an individual’s harmonic spectrum and the connectome harmonic signature of unconsciousness – even when extracted from a different dataset (DOC patients; Figure 6).

More broadly, the entire spectrum of connectome harmonics may be used to characterise the quality of various states of consciousness, in terms of being “unconscious-like” or “psychedelic-like” (or neither, as in the case of our test-retest data). Indeed, our analyses uncovered that the connectome harmonic signature of post-anaesthetic recovery (and, to a lesser extent, mild sedation), resembled the signature of the psychedelic state – even though diversity of the repertoire was near baseline levels. This intriguing observation may reflect the phenomenon whereby individuals emerging from anaesthesia can exhibit symptoms of delirium, cognitive alterations, and even hallucinations (Radtke et al., 2010; Xará et al., 2013). Thus, the similarity that we observed between anaesthetic emergence and psychedelics in terms of connectome harmonics may provide a link between the shared aspects of their phenomenology, demonstrating the usefulness of CHD for generating empirically testable predictions.

Crucially, these results demonstrate that connectome harmonics offer not only an effective one-dimensional indicator, complementing existing complexity measures (Schartner et al., 2015, 2017b, 2017a), but also a richer multi-dimensional characterisation of conscious states. Although a deeper understanding of the underlying neurophysiology will be required, it is noteworthy that individual alignment with our connectome harmonic signature of the psychedelic state correlates with the subjective intensity of the psychedelic experience induced by LSD (Figure 6), suggesting that connectome harmonics may provide a bridge between brain structure, function, and phenomenology (subjective experience).

### Role of the high-resolution representative human connectome for decomposing consciousness

It is essential to realize that the general principle of harmonic mode decomposition does *not* require the harmonic modes to be derived from the same subject who is providing the functional data. In fact, the harmonic modes do not even need to have a biological origin at all. At one extreme, researchers have successfully employed harmonics derived from a sphere to investigate how brain activity depends on the most general geometric properties of the brain and skull (Gabay and Robinson, 2017; Mukta et al., 2017). At the other extreme, investigators whose focus is subject-specific insight, rather than generalization across datasets, could perform CHD using each individual’s own connectome.

Our choice of using the harmonic modes of a high-resolution representative human connectome (replicated using data from 985 HCP subjects) enabled us to strike a balance between these two extremes, combining neurobiological insight with generalizability. On one hand, given our goal of obtaining connectome harmonic signatures of each state of consciousness that can be meaningfully compared across subjects and across datasets, it was imperative for us to use the same set of basis functions (i.e., harmonic modes of the same representative connectome) to decompose different datasets. In this respect, CHD based on a representative connectome is not conceptually different from the traditional spatially-resolved view of brain activity, which to be able to refer to the same localized region across individuals, requires spatial normalization to a standard template (e.g., MNI-152), and use of a standard parcellation, both obtained from aggregating neuroimaging data across healthy individuals (Eickhoff et al., 2018).

On the other hand, in addition to the conceptual advantages of taking into account known physical and anatomical properties of the human brain (e.g. cortical folding, local grey-matter connectivity and long-range white matter projections; see (Naze et al., 2021) for a detailed discussion), our results provided empirical demonstration that both the specific anatomical distribution of connectome harmonics, and the specific topological organization of the high-resolution human structural connectome, play a crucial role in identifying consistent patterns across different states of consciousness: perturbing either of these two aspects (via spatial rotation or randomization, respectively) obliterated the ability of harmonic decomposition to recover common patterns across the various states of consciousness considered here.

Importantly, our results revealed a prominent role of the fine-grained, high-frequency harmonics to distinguish between states of consciousness. This insight was only possible thanks to the availability of high-quality, high-resolution HCP diffusion data, which we could aggregate across subjects to obtain high-fidelity reconstructions even at the finest scale (up to three orders of magnitude more fine-grained than other approaches to harmonic mode decomposition based on the connectome, which have relied on parcellated data (Abdelnour et al., 2014; Glomb et al., 2020; Medaglia et al., 2018; Preti and Van De Ville, 2019; Wang et al., 2017; Xie et al., 2021). In fact, we successfully replicated our results using a high-resolution connectome obtained by combining 985 HCP subjects: arguably the most representative operationalization of the structural wiring of the human brain that is available to date. The need for high-quality connectome reconstruction may pose a challenge when seeking to perform CHD based on individual subjects’ connectomes, whereupon diffusion data of sufficiently high quality is not always available, and this issue could not be mitigated via aggregation across subjects. Even so, diffusion imaging and tractography are not without limitations: chief among them, their inability to infer fibre directionality, an important feature of brain wiring that will need to be accounted for through both technological and conceptual advances. Likewise, future work could extend our results by taking into account transmission delays based on tract length (Xie et al., 2021).

### Converging on consciousness

It is noteworthy that although we stratified our DOC patients based on their performance on mental imagery tasks in the scanner, our connectome harmonic analysis was entirely based on resting-state (i.e., task-free) fMRI data, which imposes no cognitive demands on patients, unlike task-based paradigms (Naci et al., 2017). Connectome harmonics analysis of rs-fMRI may represent a useful screening tool in the clinic to identify patients for more in-depth assessment – contributing to alleviate the high rate of misdiagnoses for DOC patients when relying solely on behavioural criteria (Naci et al., 2017).

However, even failure to respond to the fMRI mental imagery tasks cannot conclusively rule out residual consciousness in DOC patients, and loss of behavioural responsiveness during anaesthesia may not always coincide with loss of brain responsiveness and subjective experience (Huang et al., 2018b; Leslie et al., 2009; Ní Mhuircheartaigh et al., 2013). More broadly, to further establish the potential clinical value of CHD as a general neural marker of consciousness, it will be essential to obtain a convergence of multi-modal markers of consciousness that bypass overt behaviour, across different neuroimaging modalities (Casali et al., 2013; Naci et al., 2017; Ní Mhuircheartaigh et al., 2013). For instance, given evidence that EEG “slow-wave activity saturation” constitutes a marker of loss of brain responsiveness induced by propofol (Ní Mhuircheartaigh et al., 2013), a promising avenue for future work will be to investigate whether a corresponding “saturation of low-frequency connectome harmonics” can be identified in the fMRI signal. Likewise, establishing the susceptibility of connectome harmonics to intervention via Transcranial Magnetic Stimulation will shed light on their suitability as a target for treatment, and enable a potential convergence with the “Perturbational Complexity Index”: one of the most sensitive indicators of consciousness available to date, which is based on how the brain’s response to TMS pulses spread (Casali et al., 2013; Casarotto et al., 2016; Comolatti et al., 2019).

## Conclusion

Overall, the energy spectrum of connectome harmonics and the diversity of their repertoire provide distinct and synergistic insights, to identify meaningful relationships between brain function, its network structure, and conscious experience. Having demonstrated the generalisability of connectome harmonic decomposition across datasets and states of consciousness, our results lay the groundwork for future harmonic-based quantitative comparison of different mental states in health and disease.

## Data Availability

The raw data analysed during the current study are available on request from the following authors. Propofol anaesthesia, Disorders of Consciousness and test-retest datasets: Dr. Emmanuel A. Stamatakis (University of Cambridge, Division of Anaesthesia). LSD dataset: Dr. Robin L. Carhart-Harris (Imperial College London/University of California – San Francisco). Ketamine dataset: Dr. Ram Adapa (University of Cambridge, Division of Anaesthesia). The Human Connectome Project datasets are freely available from http://www.humanconnectome.org/.

## Code Availability

The CONN toolbox is freely available online (http://www.nitrc.org/projects/conn). The Java Information Dynamics Toolbox is freely available online: https://github.com/jlizier/jidt. DSI Studio is freely available online: http://dsi-studio.labsolver.org. Lead-DBS is freely available online: http://www.lead-dbs.org. The Brain Connectivity Toolbox is freely available online: https://sites.google.com/site/bctnet/. Code for the spherical rotations is freely available online: github.com/spin-test/spin-test.

## Author Contributions

AIL, JV, EAS, SA and MLK conceived the study. AIL, JV, PAMM, SA, MLK and EAS designed the methodology and the analysis. AIL analysed the data. JV, PAMM, MMC, IP, ARDP contributed to data analysis. PF, GBW, JA, JDP, RA, DKM, EAS, LR, RLCH, BJS were involved in designing the original studies for which the present data were collected. PF, MMC, GBW, JA, EAS, RA, AEM, RLCH, LR all participated in data collection. AIL, EAS and SA wrote the manuscript with feedback from all co-authors.

## Acknowledgments

AIL, JV and PAMM would like to thank Lena Dorfschmidt for co-organising OxBridge BrainHack, where this work began. We also thank all volunteers and patients who provided data. This work was supported by grants from The Wellcome Trust Research Training Fellowship (grant no. 083660/Z/07/Z), Raymond and Beverly Sackler Studentship, and the Cambridge Commonwealth Trust [RA]; the UK Medical Research Council (U.1055.01.002.00001.01) [JDP]; The James S. McDonnell Foundation [JDP]; the Canadian Institute for Advanced Research (CIFAR; grant RCZB/072 RG93193) [to DKM and EAS]; The National Institute for Health Research (NIHR, UK), Cambridge Biomedical Research Centre and NIHR Senior Investigator Awards [JDP and DKM]; The British Oxygen Professorship of the Royal College of Anaesthetists [DKM]; The Stephen Erskine Fellowship, Queens’ College, University of Cambridge [EAS]; The Evelyn Trust, Cambridge and the EoE CLAHRC fellowship [JA]; The Gates Cambridge Trust [AIL]; The Cambridge International Trust and the Howard Sidney Sussex Studentship [MMC]; The Oon Khye Beng Ch’Hia Tsio Studentship for Research in Preventive Medicine, Downing College, University of Cambridge [IP]; The Wellcome Trust (grant no. 210920/Z/18/Z) [PAMM]; The European Research Council Consolidator Grant CAREGIVING (615539) [MLK and SA]; The Center for Music in the Brain, funded by the Danish National Research Foundation (DNRF117) [MLK, SA and JV]; The Centre for Eudaimonia and Human Flourishing, funded by the Pettit and Carlsberg Foundations [MLK]; The Imperial College President’s Scholarship [LR]; The Alex Mosley Charitable Trust [RLCH]; The ketamine study was funded by the Bernard Wolfe Health Neuroscience Fund and the Wellcome Trust. The original LSD study received support from a Crowd Funding Campaign and the Beckley Foundation, as part of the Beckley-Imperial Research Programme. The research was also supported by the NIHR Brain Injury Healthcare Technology Co-operative based at Cambridge University Hospitals NHS Foundation Trust and University of Cambridge. Data used to obtain the human connectome were provided by the Human Connectome Project, WU-Minn Consortium (Principal Investigators: David Van Essen and Kamil Ugurbil; 1U54MH091657) funded by the 16 NIH Institutes and Centers that support the NIH Blueprint for Neuroscience Research; and by the McDonnell Center for Systems Neuroscience at Washington University.

## Footnotes

1 A fundamentally different way of relating structure and function – including in different states of consciousness – is by means of whole-brain computational models (Cofré et al., 2020; Kringelbach and Deco, 2020; Luppi et al., 2021d)(Cofré et al., 2020; Kringelbach and Deco, 2020; Luppi et al., 2021d). Rather than focusing on the direct quantification of structure-function correspondence in empirical data, these approaches aim to simulate brain activity from structural connectivity.

