## Supplementary Information for "Distributed harmonic patterns of structure-function dependence orchestrate human consciousness"

### Supplementary Figures

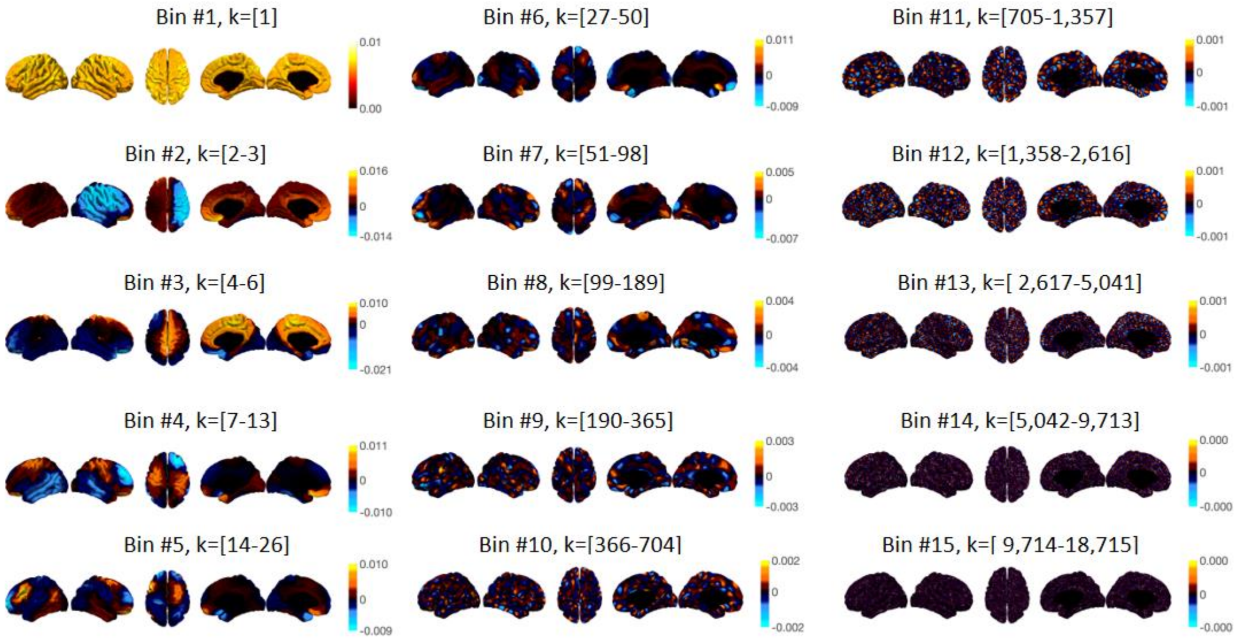


**Figure S1**. **Binned connectome harmonics**. Surface projections of connectome harmonics averaged over each of 15 logarithmically spaced bins (with corresponding wavenumbers *k* indicated in braces), showing the progressive increase in complexity and granularity of the connectome harmonic patterns, with increasing spatial frequency.

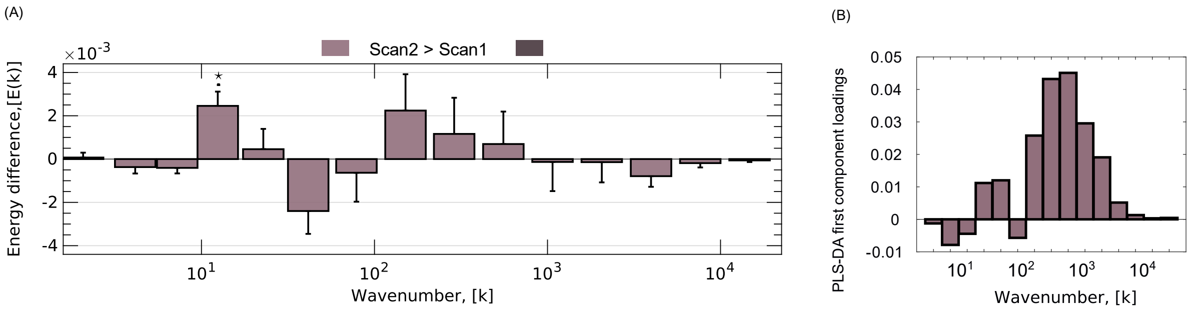


**Figure S2**. **Stable energy levels across two scans of the same individuals**. (A) Differences in frequency-specific energy of connectome harmonics for two different scans of the same individuals. (B) Multivariate energy signature (MVS) for discriminating between the first and second scan. * $p < 0.05$ (FDR-corrected across 15 frequency bins, for the frequency-specific analysis).

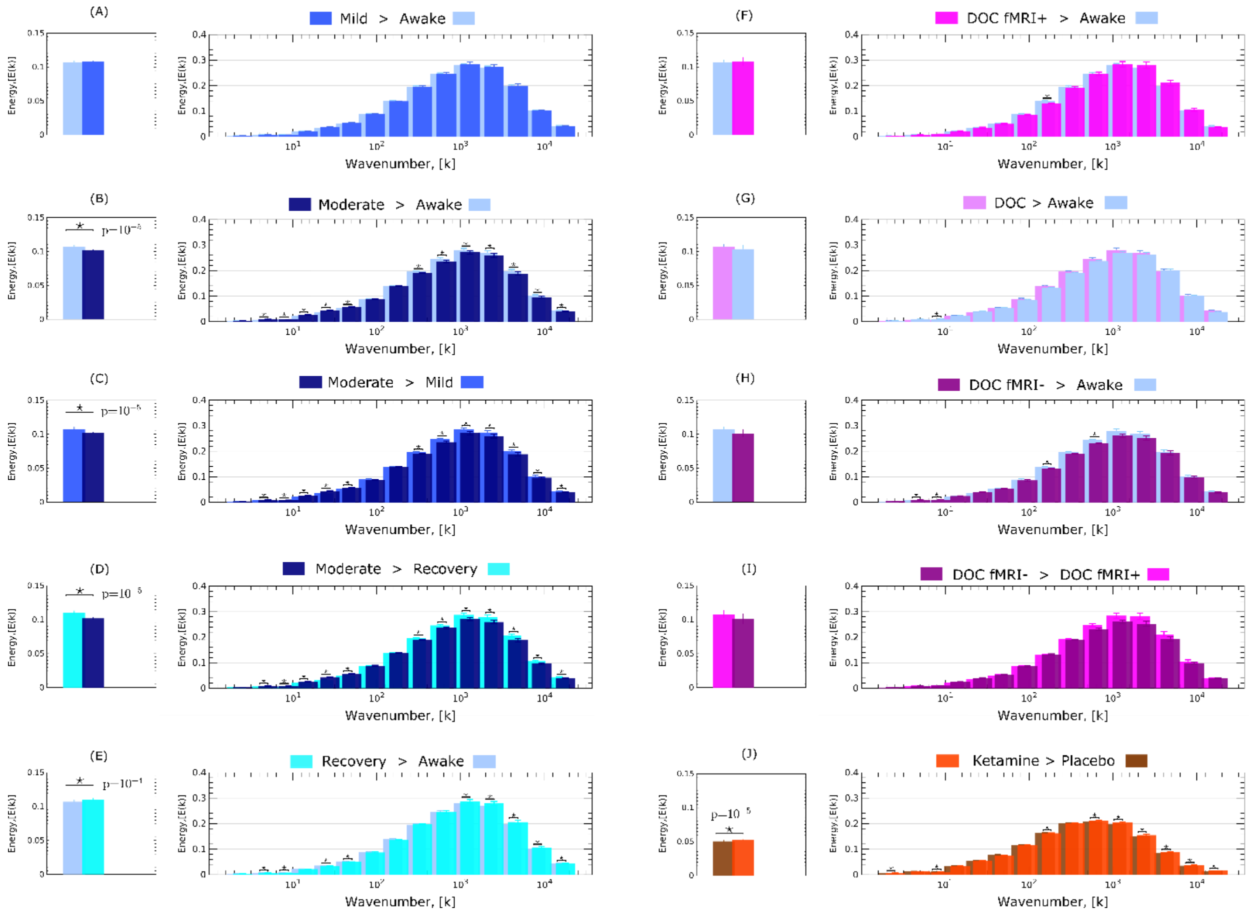


**Figure S3**. **Energy levels across states of consciousness**. (A) Total energy (left) and frequency-specific energy of connectome harmonics (right) for Mild propofol sedation vs wakefulness. (B) Total energy (left) and frequency-specific energy of connectome harmonics (right) for Moderate anaesthesia vs wakefulness. (C) Total energy (left) and frequency-specific energy of connectome harmonics (right) for Moderate anaesthesia vs mild sedation. (D) Total energy (left) and frequency-specific energy of connectome harmonics (right) for Moderate anaesthesia vs post-anaesthetic recovery. (E) Total energy (left) and frequency-specific energy of connectome harmonics (right) for Recovery vs wakefulness. (F) Total energy (left) and frequency-specific energy of connectome harmonics (right) for DOC patients vs awake healthy controls. (G) Total energy (left) and frequency-specific energy of connectome harmonics (right) for DOC fMRI+ patients vs awake healthy controls. (H) Total energy (left) and frequency-specific energy of connectome harmonics (right) for DOC fMRI- patients vs awake healthy controls. (I) Total energy (left) and frequency-specific energy of connectome harmonics (right) for fMRI- vs fMRI+ DOC patients. (J) Ketamine > placebo. * $p < 0.05$ (FDR-corrected across 15 frequency bins, for the frequency-specific analysis).

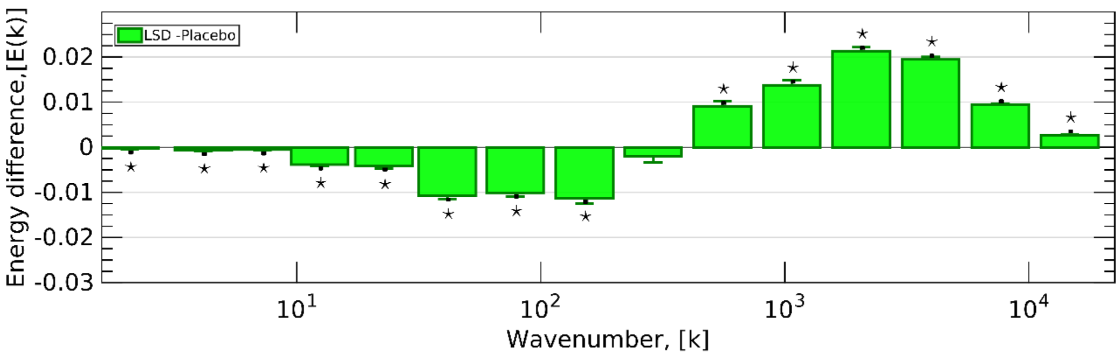


**Figure S4**. **Connectome harmonic energy signature of LSD.** Re-derived from the same data used by Atasoy and colleagues (2017). * $p < 0.05$, FDR-corrected across 15 frequency bins.

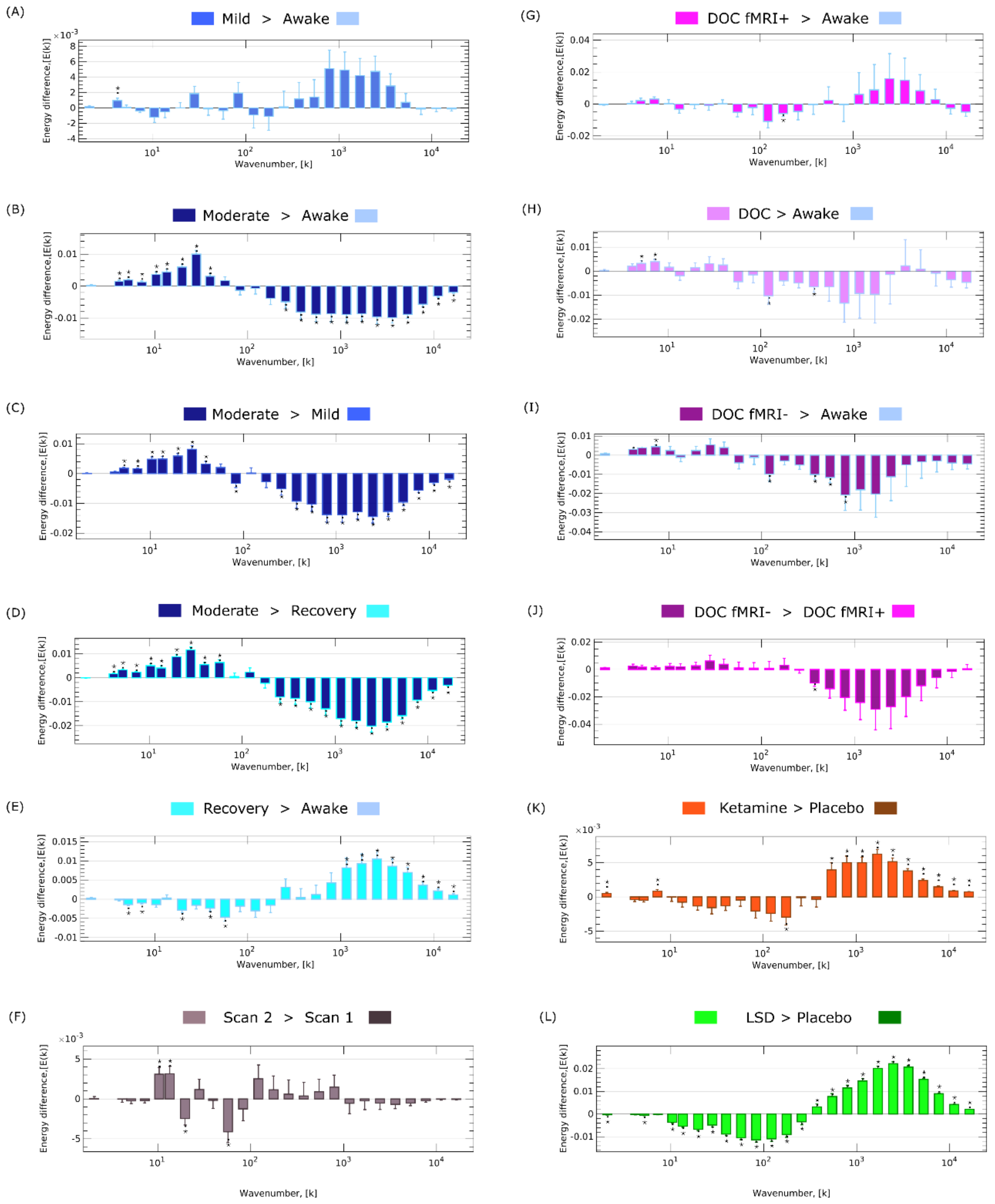


**Figure S5**. **Replication of frequency-specific changes of connectome harmonic energy across states of consciousness when using 25 bins.** (A) Mild propofol sedation > wakefulness. (B) Moderate anaesthesia > wakefulness. (C) Moderate anaesthesia > mild sedation. (D) Moderate anaesthesia > post-anaesthetic recovery. (E) Recovery > wakefulness. (F) Test-retest scan2 > scan 1. (G) DOC patients > awake healthy controls. (H) DOC fMRI+ patients > awake healthy controls. (I) DOC fMRI- patients > awake healthy controls. (J) fMRI- > fMRI+ DOC patients. (K) Ketamine > placebo. (L) LSD > placebo. * $p < 0.05$, FDR-corrected across 25 logarithmically spaced frequency bins.

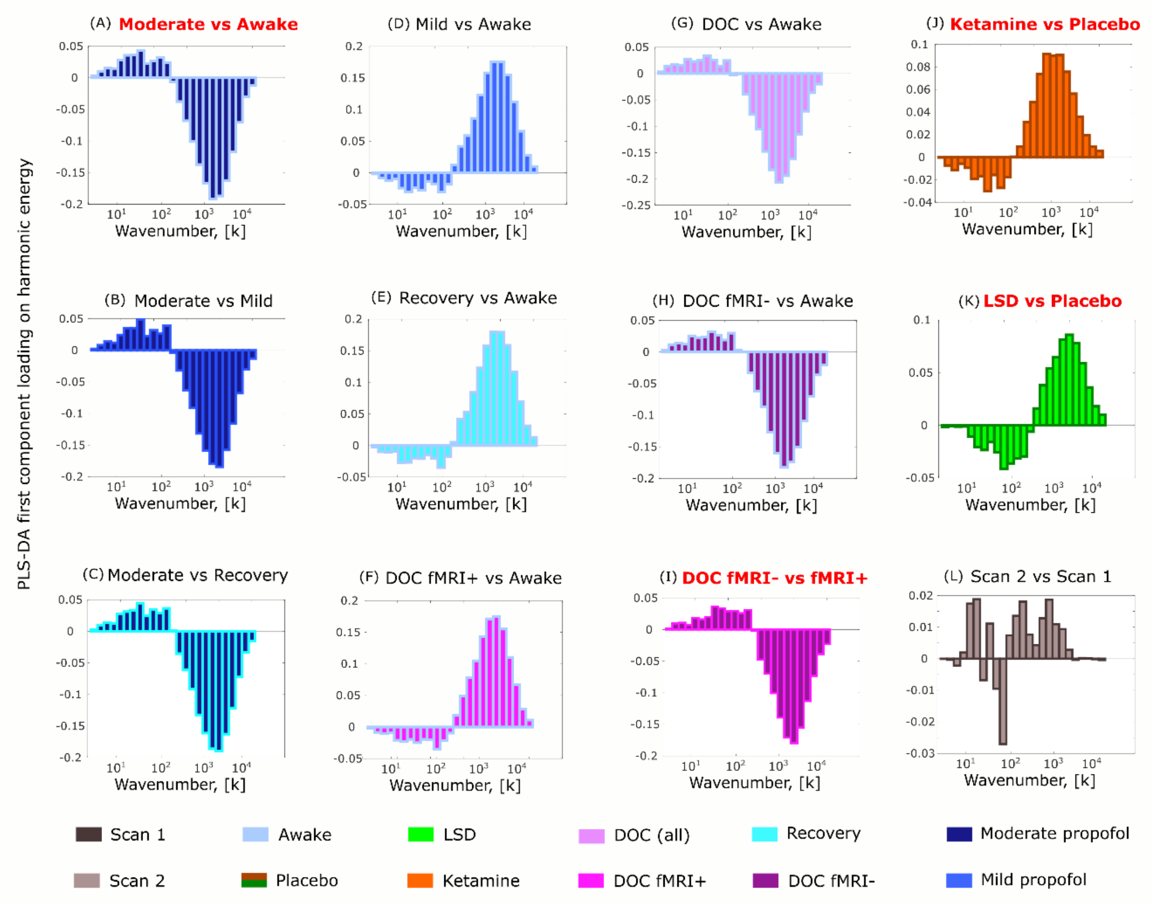


**Figure S6. Replication of multivariate signatures of connectome harmonic energy for different states of consciousness, when using 25 bins.** (A) Moderate anaesthesia > wakefulness. (B) Moderate anaesthesia > mild sedation. (C) Moderate anaesthesia > post-anaesthetic recovery. (D) Mild sedation > wakefulness. (E) Post-anaesthetic recovery > wakefulness. (F) DOC fMRI+ patients > awake healthy controls. (G) DOC patients > awake healthy controls. (H) DOC fMRI- patients > awake healthy controls. (I) fMRI- > fMRI+ DOC patients. (J) Ketamine > placebo. (K) LSD > placebo. (L) Test-retest scan2 > scan 1. Bar colour indicates the target state; contours indicate the reference state.

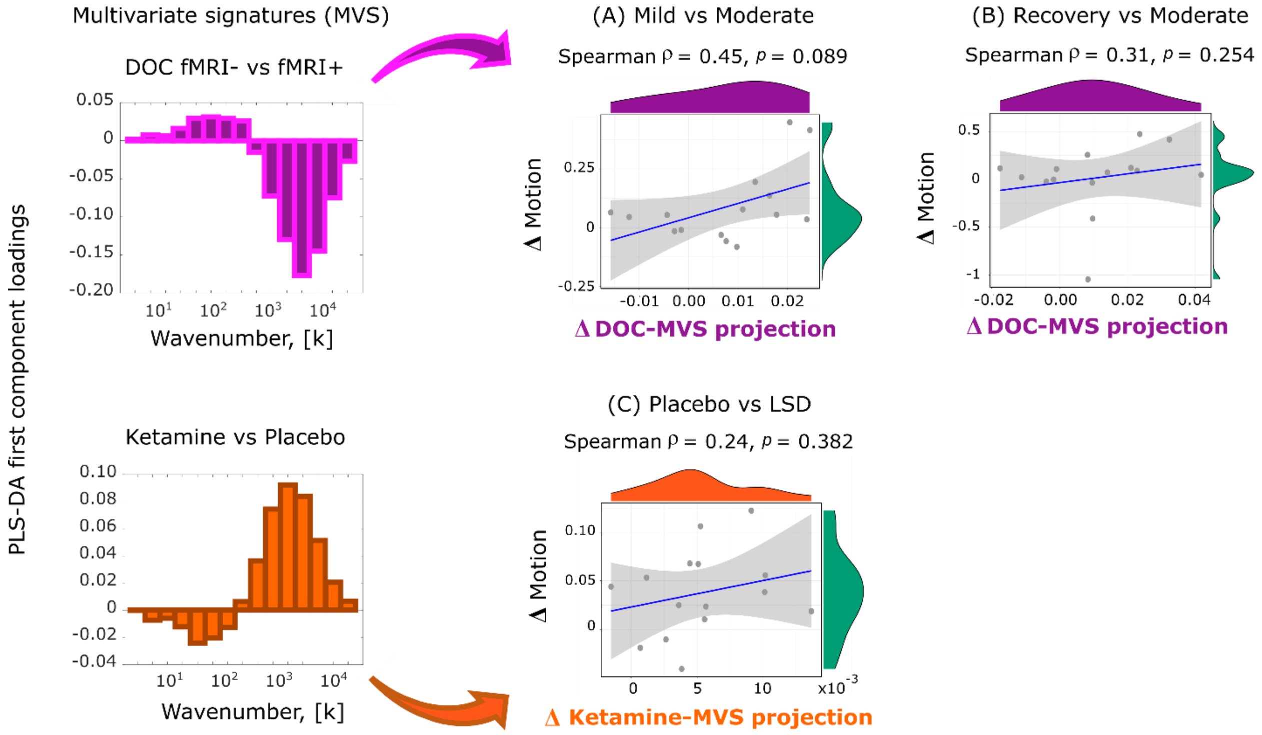


**Figure S7. Change in projection onto cross-dataset multivariate energy signature (MVS) does not significantly correlate with differences in subject motion in the scanner.** (A) Scatterplot of delta in connectome harmonic energy projection onto the MVS derived from the DOC dataset, versus the delta in head motion (moderate anaesthesia minus mild anaesthesia). (B) Scatterplot of delta in connectome harmonic energy projection onto the MVS derived from the DOC dataset, versus the delta in head motion (moderate anaesthesia minus recovery). (C) Scatterplot of delta in connectome harmonic energy projection onto the MVS derived from the ketamine dataset, versus the delta in head motion (LSD minus placebo).

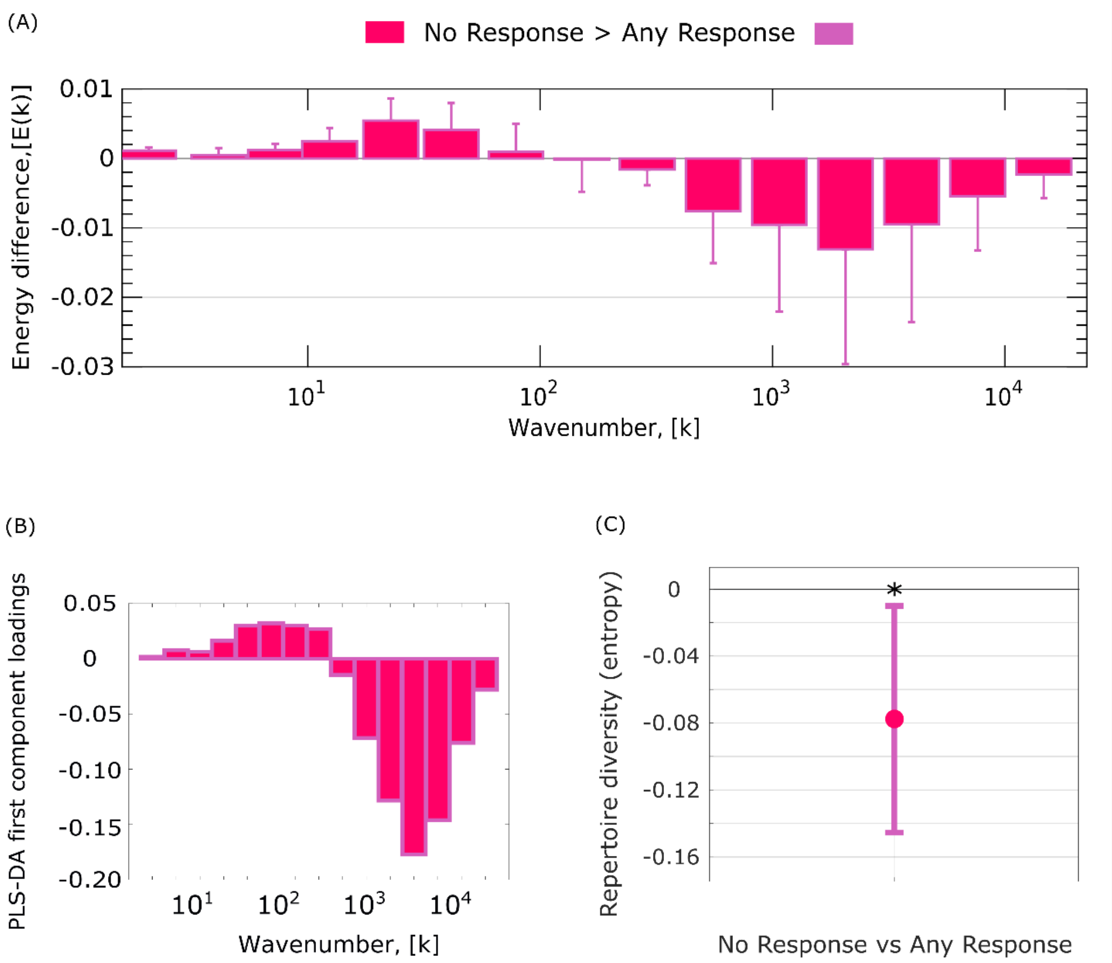


**Figure S8**. **Connectome harmonic differences between DOC patients based on combined clinical and fMRI classification.** (A) Differences in frequency-specific energy of connectome harmonics between patients who are both UWS and fMRI- (“No response”, N=8) and all other patients (“Any response”, N=14). (B) Multivariate energy signature (MVS) for best discriminating between the two groups of patients. (C) Diversity (entropy) of the full connectome harmonic repertoire is significantly diminished in fMRI- UWS patients compared with the other DOC patients. * $p < 0.05$.

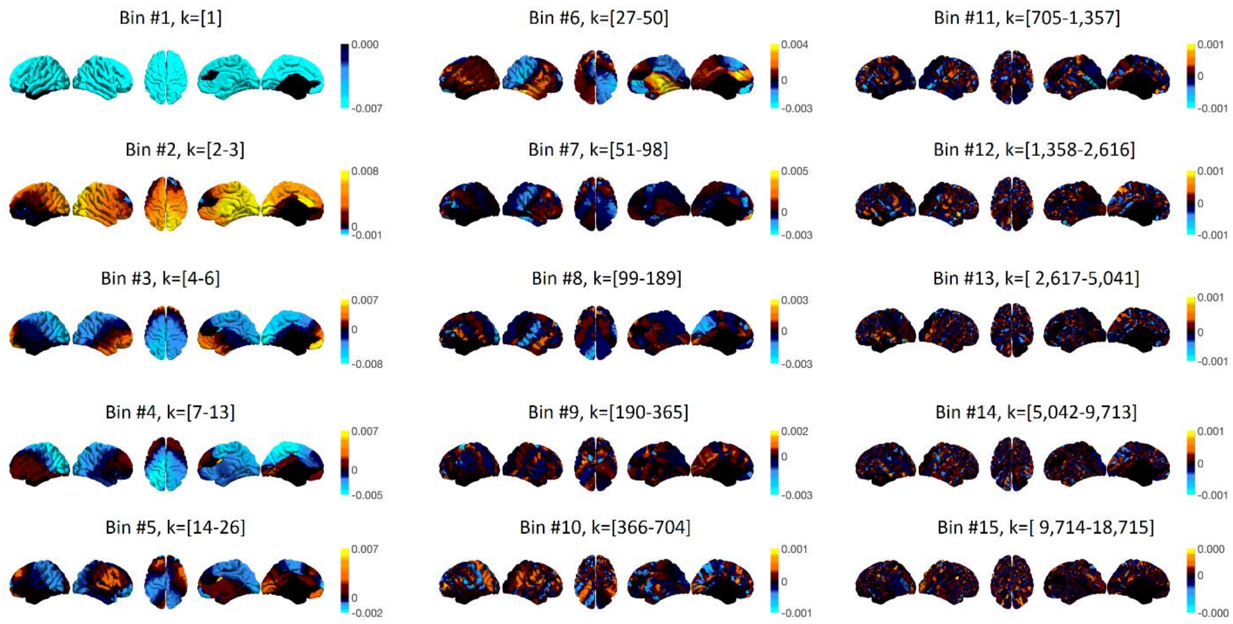


**Figure S9**. **Rotated connectome harmonics**. Surface projections of connectome harmonics averaged over each of 15 logarithmically spaced bins (with corresponding wavenumbers *k* indicated in braces), showing the progressive increase in complexity and granularity of the connectome harmonic patterns, with increasing spatial frequency. Each harmonic was projected on a sphere and the sphere was then randomly rotated before back-projecting each harmonic on the brain surface (Methods).

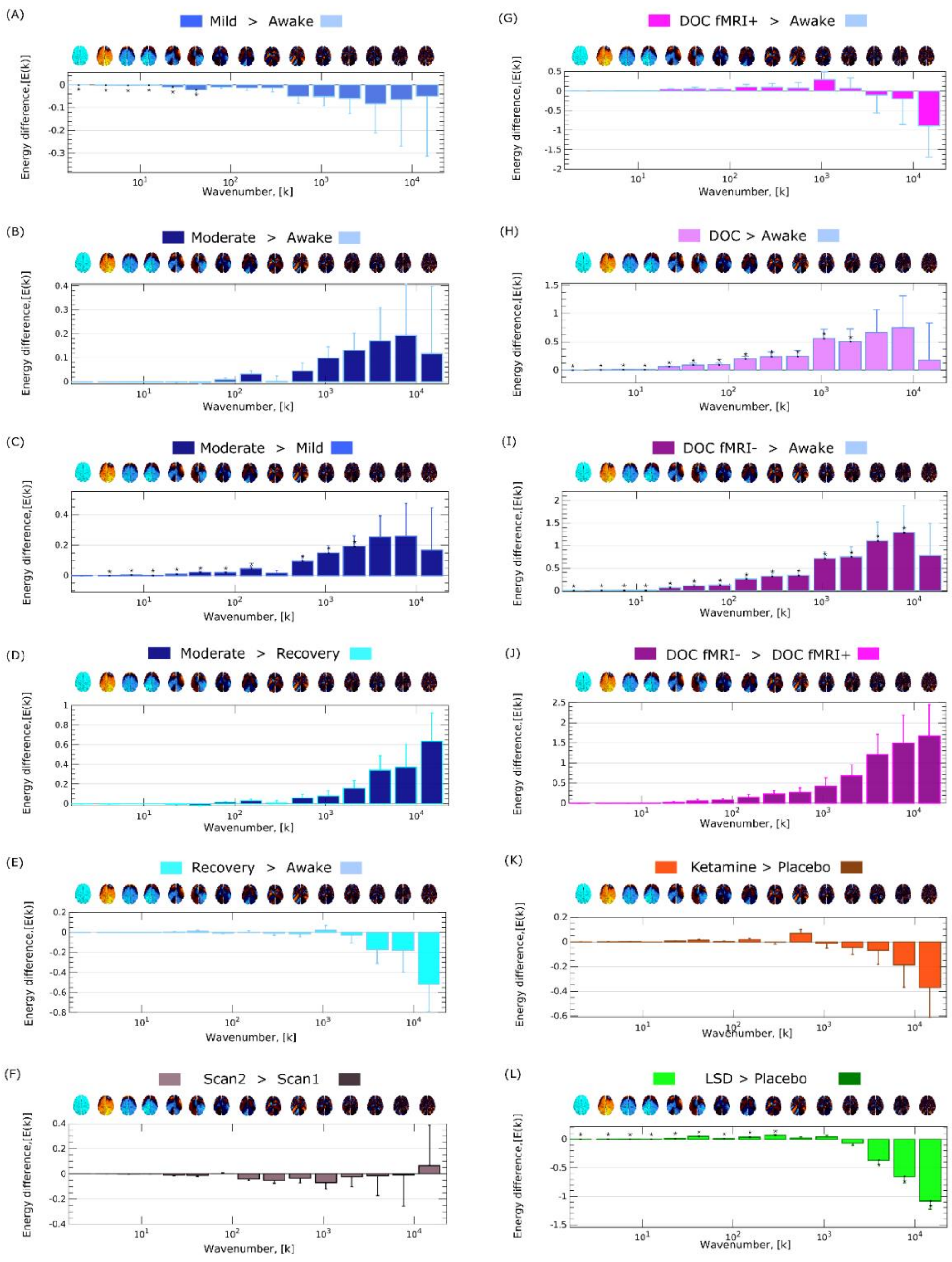


**Figure S10**. **No consistent frequency-specific energy changes are observed across states of consciousness, for the rotated harmonics.** (A) Mild propofol sedation > wakefulness. (B) Moderate anaesthesia > wakefulness. (C) Moderate anaesthesia > mild sedation. (D) Moderate anaesthesia > post-anaesthetic recovery. (E) Recovery > wakefulness. (F) Test-retest scan2 > scan 1. (G) DOC patients > awake healthy controls. (H) DOC fMRI+ patients > awake healthy controls. (I) DOC fMRI- patients > awake healthy controls. (J) fMRI- > fMRI+ DOC patients. (K) Ketamine > placebo. (L) LSD > placebo. * $p < 0.05$, FDR-corrected across 15 frequency bins. A brain surface projection of the connectome harmonic pattern corresponding to each frequency bin, averaged over the constituent spatial frequencies, is shown above each bin.

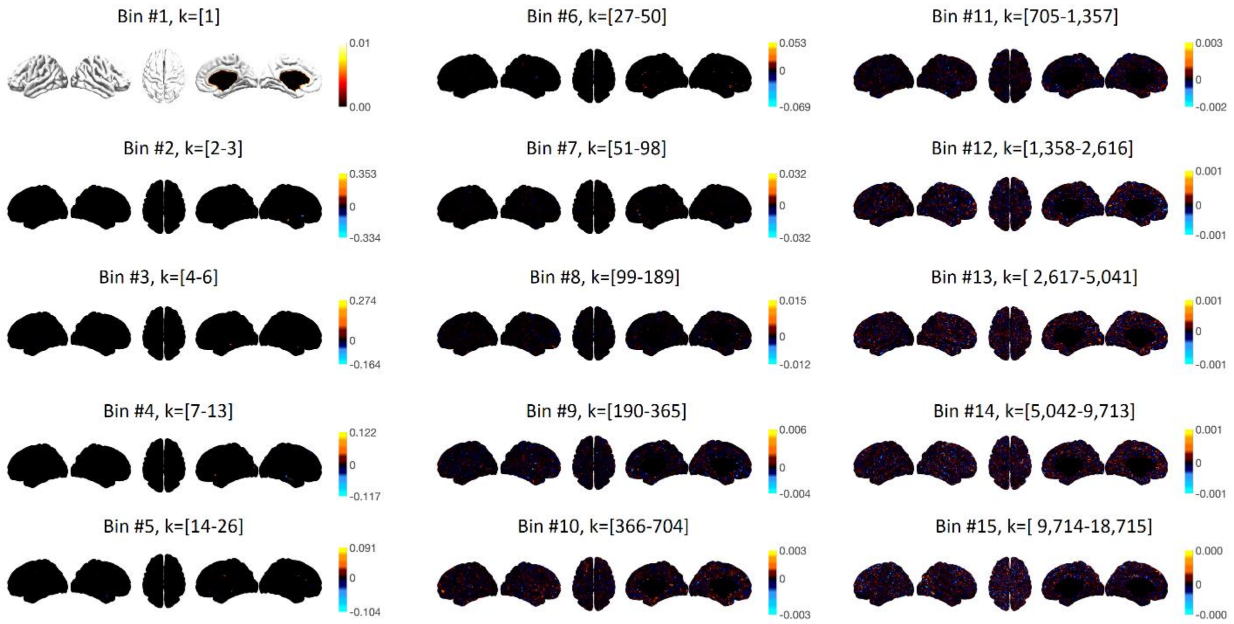


**Figure S11**. **Binned connectome harmonics from a randomised connectome**. Surface projections of connectome harmonics averaged over each of 15 logarithmically spaced bins (with corresponding wavenumbers *k* indicated in braces; note that only the first 14 bins are used for analysis). Note how the colour is largely uniform as the harmonics obtained from the randomised connectome do not display clear spatial patterns.

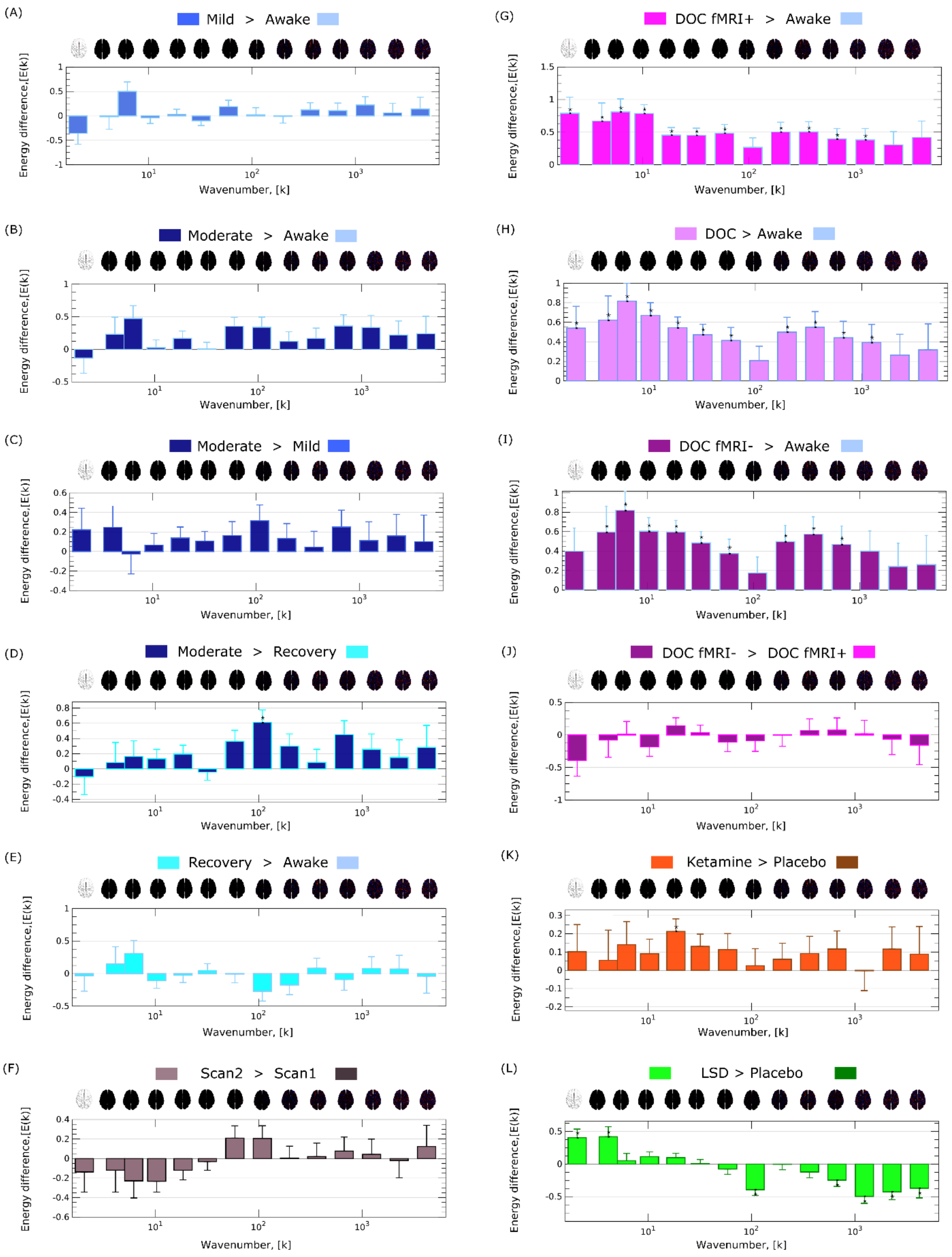


**Figure S12. No consistent frequency-specific energy changes are observed across states of consciousness, for the randomised connectome.** (A) Mild propofol sedation > wakefulness. (B) Moderate anaesthesia > wakefulness. (C) Moderate anaesthesia > mild sedation. (D) Moderate anaesthesia > post-anaesthetic recovery. (E) Recovery > wakefulness. (F) Test-retest scan2 > scan 1. (G) DOC patients > awake healthy controls. (H) DOC fMRI+ patients > awake healthy controls. (I) DOC fMRI- patients > awake healthy controls. (J) fMRI- > fMRI+ DOC patients. (K) Ketamine > placebo. (L) LSD > placebo. * $p < 0.05$, FDR-corrected across 14 frequency bins. A brain surface projection of the connectome harmonic pattern corresponding to each frequency bin, averaged over the constituent spatial frequencies, is shown above each bin.

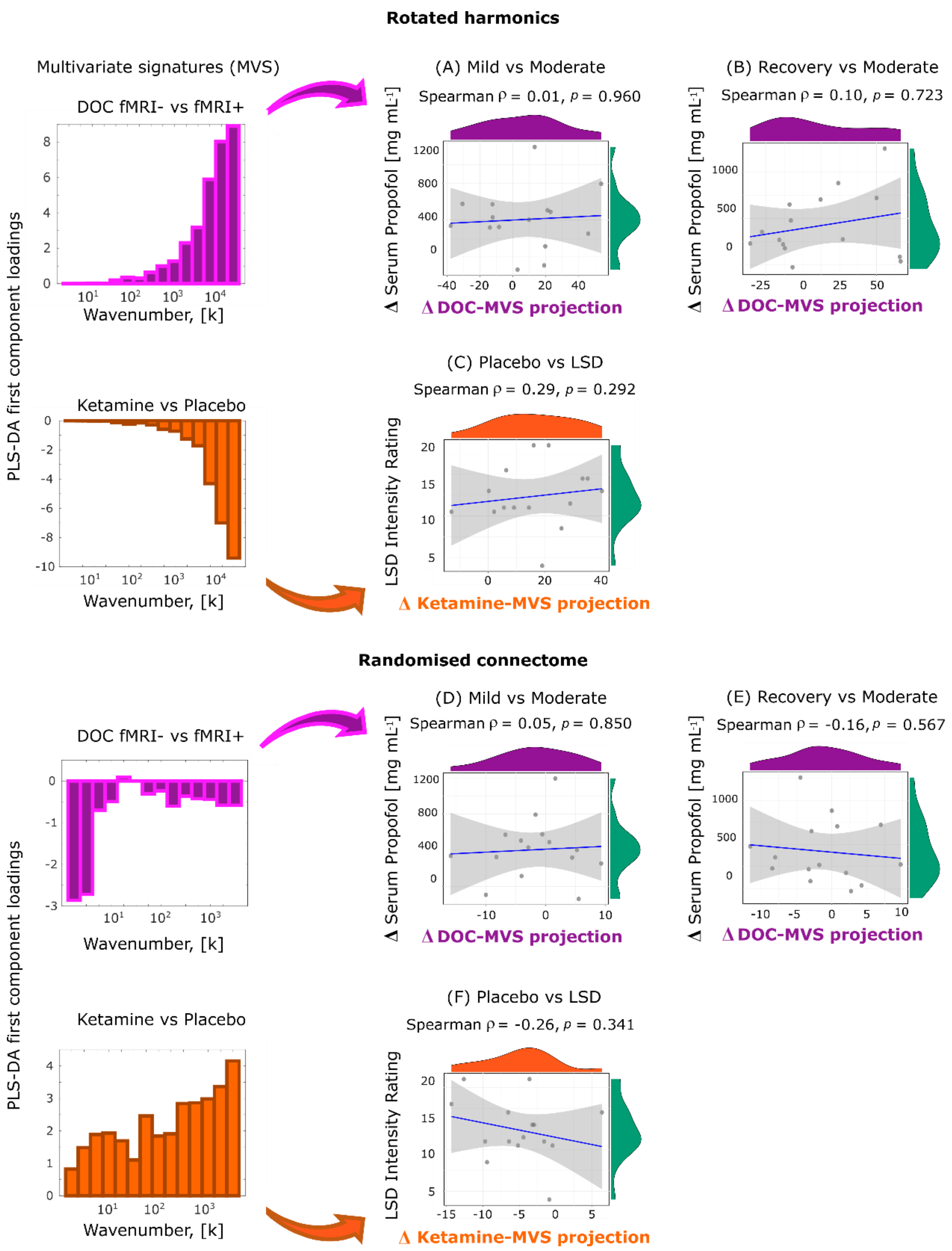


**Figure S13. No generalisation of connectome harmonic signatures is observed when using rotated harmonics or a randomised connectome.** For (A-C), the connectome harmonic energy spectrum is obtained from rotated harmonics. For (D-E), it is obtained from the harmonics of a randomised connectome.

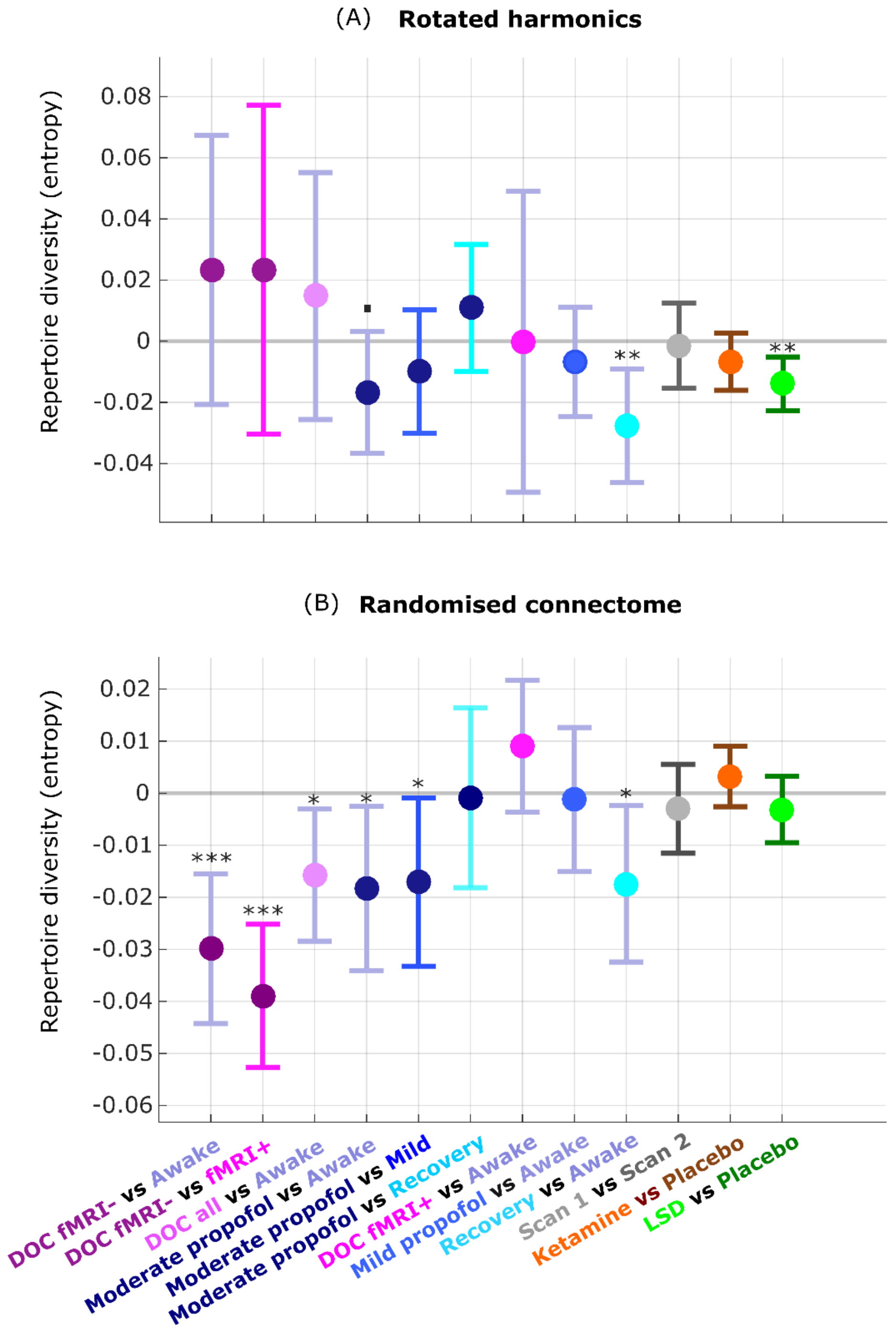


**Figure S14. Repertoire diversity does not track level of consciousness across datasets, when obtained from rotated harmonics (A) or from a randomised connectome (B).** (A) Diversity of connectome harmonic repertoire obtained from rotated harmonics. (B) Diversity of connectome harmonic repertoire obtained from a randomised connectome. *** $p < 0.001$; ** $p < 0.01$; * $p < 0.05$; . $p < 0.10$.

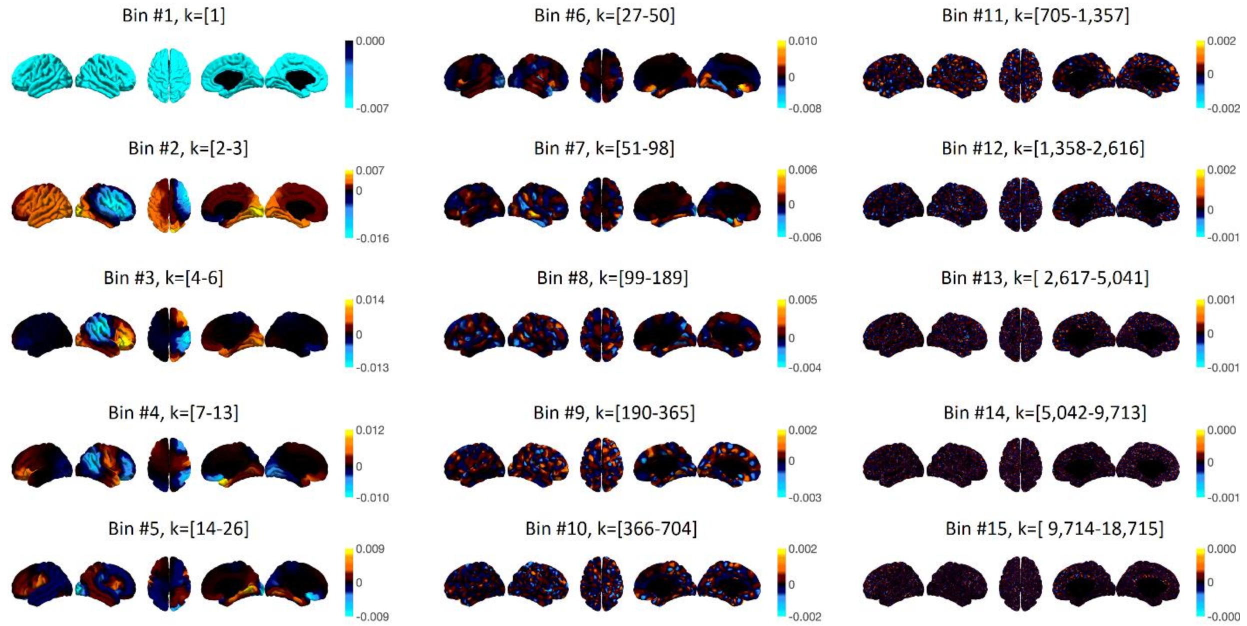


**Figure S15**. **Binned connectome harmonics for the HCP-985 connectome**. Surface projections of connectome harmonics averaged over each of 15 logarithmically spaced bins (with corresponding wavenumbers *k* indicated in braces), showing the progressive increase in complexity and granularity of the connectome harmonic patterns, with increasing spatial frequency. Note that only the first 14 bins are used for analysis.

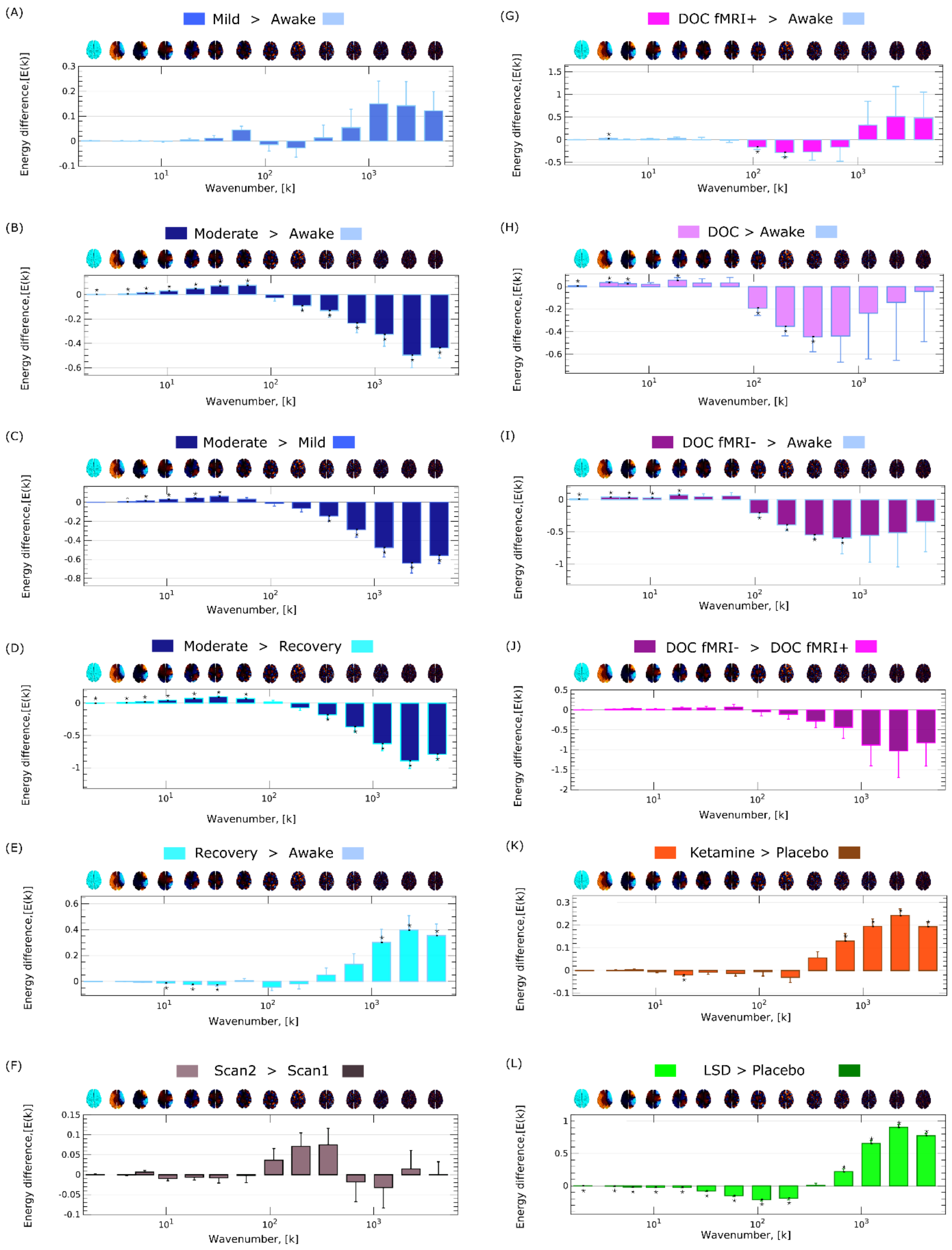


**Figure S16**. **Frequency-specific energy changes across states of consciousness, for the HCP-985 connectome.** (A) Mild propofol sedation > wakefulness. (B) Moderate anaesthesia > wakefulness. (C) Moderate anaesthesia > mild sedation. (D) Moderate anaesthesia > post-anaesthetic recovery. (E) Recovery > wakefulness. (F) Test-retest scan2 > scan 1. (G) DOC patients > awake healthy controls. (H) DOC fMRI+ patients > awake healthy controls. (I) DOC fMRI- patients > awake healthy controls. (J) fMRI- > fMRI+ DOC patients. (K) Ketamine > placebo. (L) LSD > placebo. * $p < 0.05$, FDR-corrected across 14 frequency bins. A brain surface projection of the connectome harmonic pattern corresponding to each frequency bin, averaged over the constituent spatial frequencies, is shown above each bin.

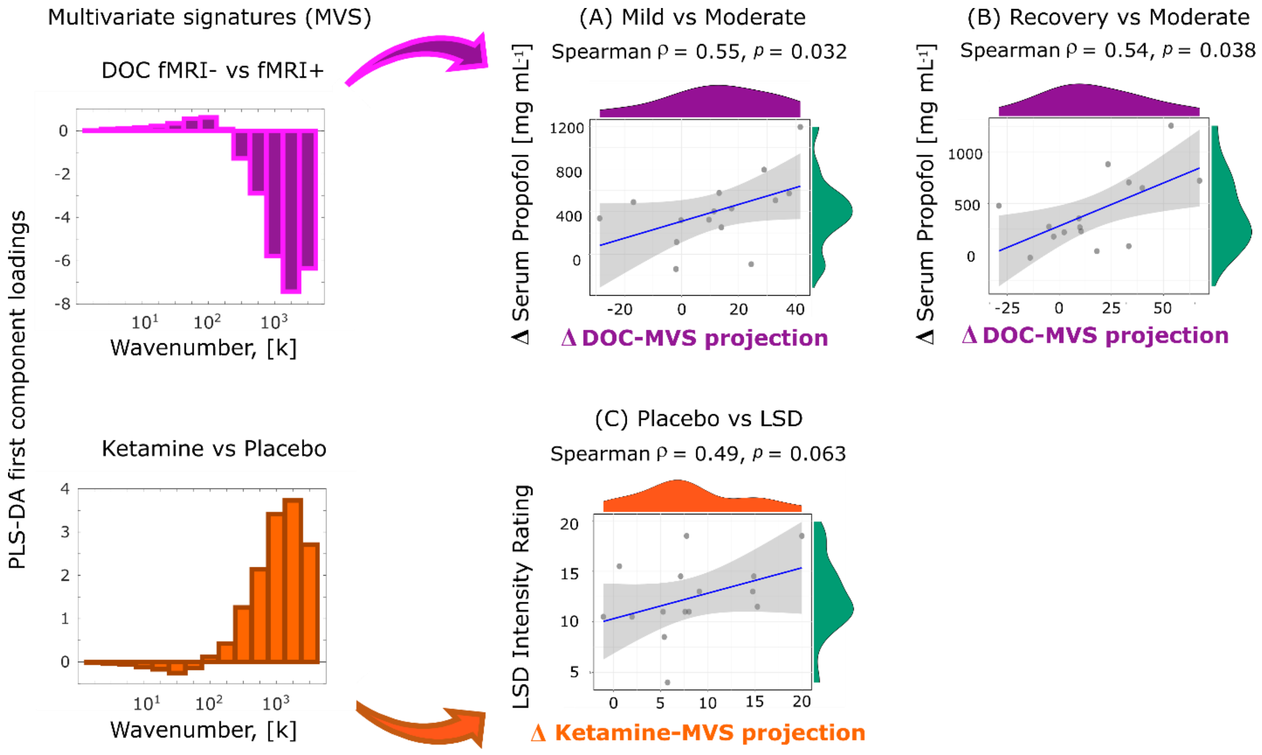


**Figure S17. Correlations between multivariate energy signatures (MVS) and propofol levels and LSD intensity scores, are replicated with the HCP-985 connectome.** (A) Scatterplot of the change (moderate anaesthesia minus mild) in connectome harmonic energy projection onto the multivariate energy signature (MVS) derived from the DOC dataset, versus the change in propofol levels in volunteers' blood serum, between mild and moderate propofol anaesthesia. (B) Scatterplot of the change (moderate minus recovery) in connectome harmonic energy projection onto the multivariate signature derived from the DOC dataset, versus the change in propofol levels in volunteers' blood serum, between moderate anaesthesia and recovery. (C) Scatterplot of the change (LSD minus placebo) in connectome harmonic energy projection onto the multivariate signature derived from the ketamine dataset, versus the subjective intensity of the psychedelic experience induced by LSD.

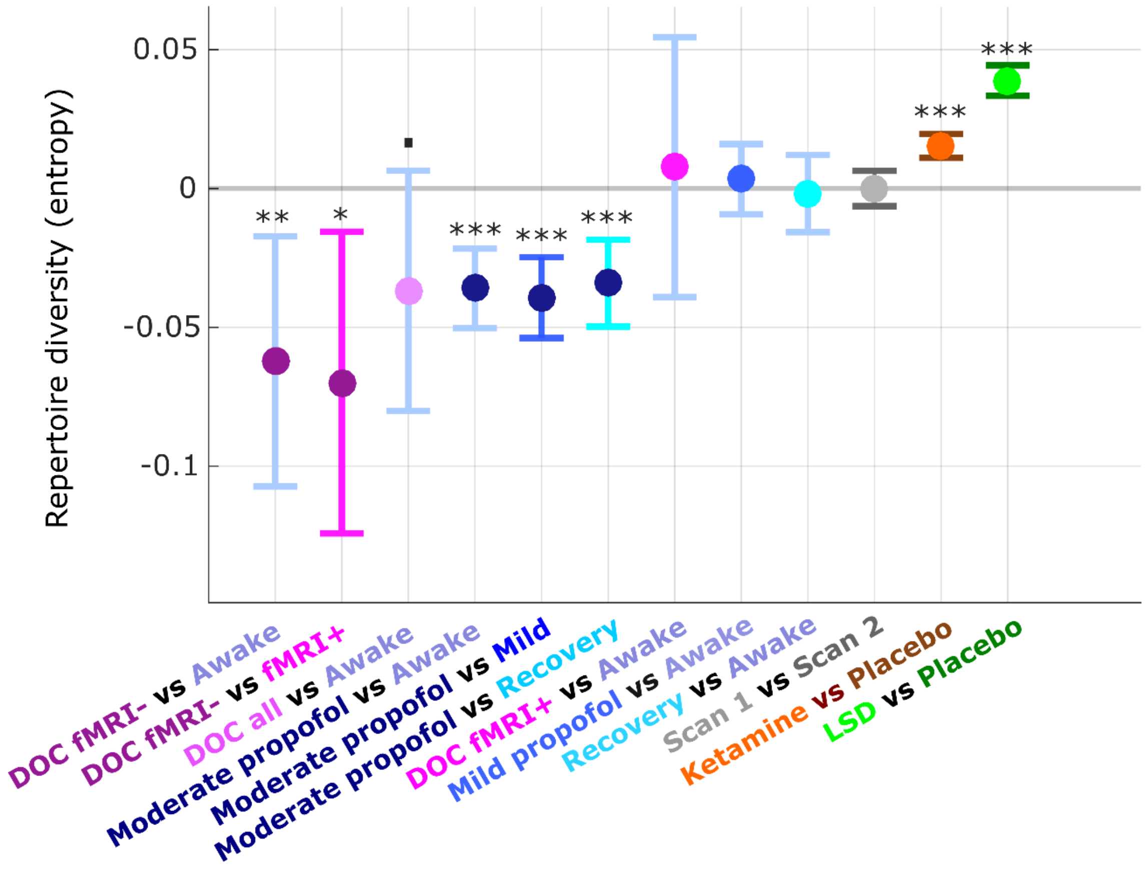


**Figure S18. Replication of repertoire diversity results for the HCP-985 connectome.** Fixed effects (and 95% CI) of the comparison in repertoire diversity (entropy of connectome harmonic power distribution) between pairs of conditions (states of consciousness), treating condition as a fixed effect, and subjects as random effects. Timepoints were also included as random effects, nested within subjects. *** $p < 0.001$; ** $p < 0.01$; * $p < 0.05$.

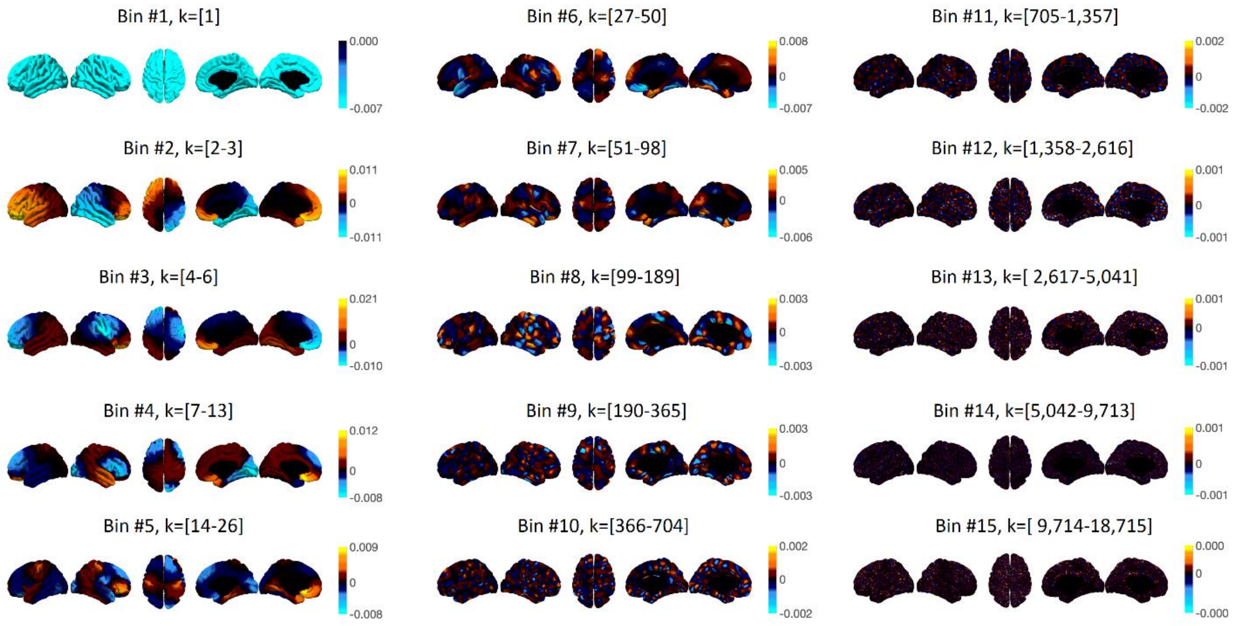


**Figure S19**. **Binned connectome harmonics for the MGH-32 connectome**. Surface projections of connectome harmonics averaged over each of 15 logarithmically spaced bins (with corresponding wavenumbers *k* indicated in braces), showing the progressive increase in complexity and granularity of the connectome harmonic patterns, with increasing spatial frequency. Note that only the first 14 bins are used for analysis.

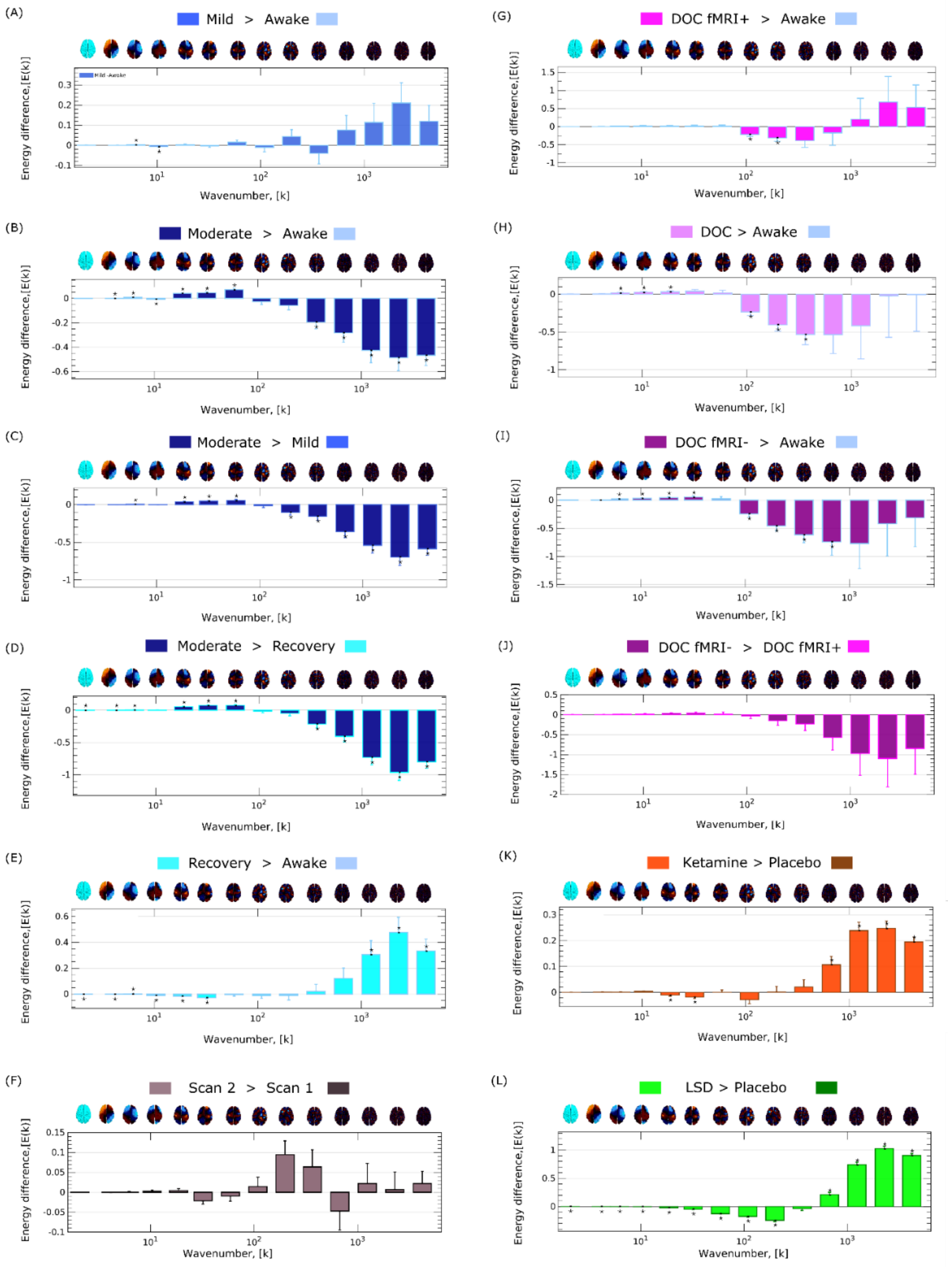


**Figure S20**. **Frequency-specific energy changes across states of consciousness, for the MGH-32 connectome.** (A) Mild propofol sedation > wakefulness. (B) Moderate anaesthesia > wakefulness. (C) Moderate anaesthesia > mild sedation. (D) Moderate anaesthesia > post-anaesthetic recovery. (E) Recovery > wakefulness. (F) Test-retest scan2 > scan 1. (G) DOC patients > awake healthy controls. (H) DOC fMRI+ patients > awake healthy controls. (I) DOC fMRI- patients > awake healthy controls. (J) fMRI- > fMRI+ DOC patients. (K) Ketamine > placebo. (L) LSD > placebo. * $p < 0.05$, FDR-corrected across 14 frequency bins. A brain surface projection of the connectome harmonic pattern corresponding to each frequency bin, averaged over the constituent spatial frequencies, is shown above each bin.

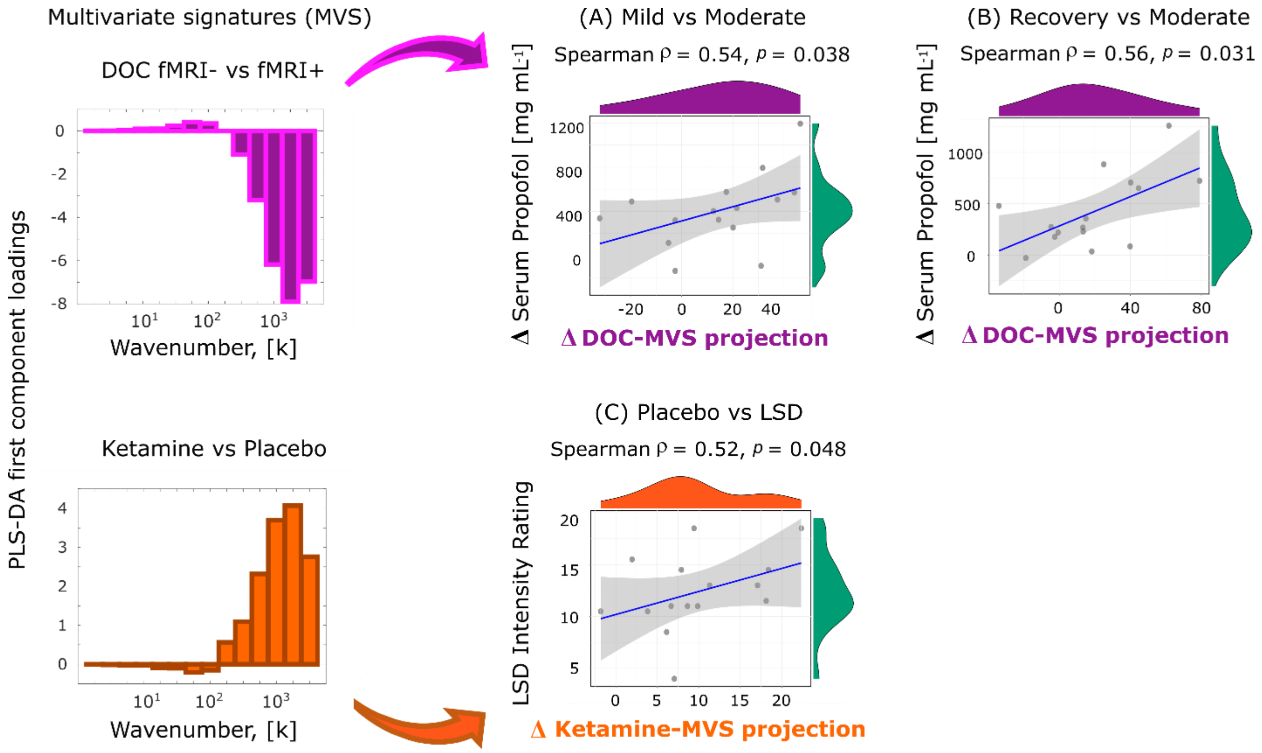


**Figure S21. Correlations between multivariate energy signatures (MVS) and propofol levels and LSD intensity scores, are replicated with the MGH-32 connectome.** (A) Scatterplot of the change (moderate anaesthesia minus mild) in connectome harmonic energy projection onto the multivariate energy signature (MVS) derived from the DOC dataset, versus the change in propofol levels in volunteers' blood serum, between mild and moderate propofol anaesthesia. (B) Scatterplot of the change (moderate minus recovery) in connectome harmonic energy projection onto the multivariate signature derived from the DOC dataset, versus the change in propofol levels in volunteers' blood serum, between moderate anaesthesia and recovery. (C) Scatterplot of the change (LSD minus placebo) in connectome harmonic energy projection onto the multivariate signature derived from the ketamine dataset, versus the subjective intensity of the psychedelic experience induced by LSD.

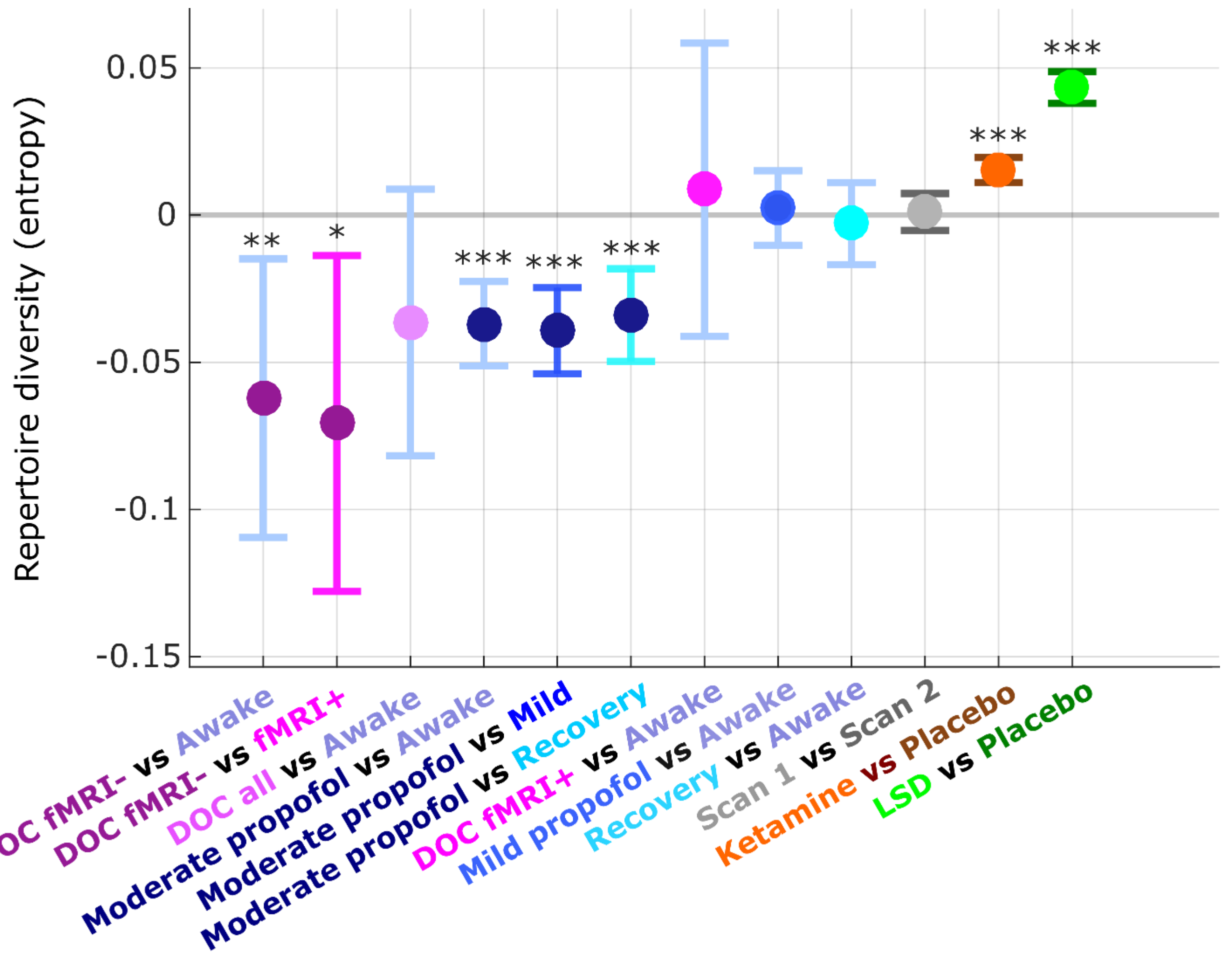


**Figure S22. Replication of repertoire diversity results for the MGH-32 connectome.** Fixed effects (and 95% CI) of the comparison in repertoire diversity (entropy of connectome harmonic power distribution) between pairs of conditions (states of consciousness), treating condition as a fixed effect, and subjects as random effects. Timepoints were also included as random effects, nested within subjects. *** $p < 0.001$; ** $p < 0.01$; * $p < 0.05$.

### Supplementary Tables

| Contrast | Fixed Effect | 95% CI Lower | 95% CI Upper | p-value | Significance |
|---|---|---|---|---|---|
| **Awake vs fMRI-** | -5.598 | -15.603 | 4.406 | 0.273 | n.s. |
| **fMRI+ vs fMRI-** | -6.835 | -19.879 | 6.209 | 0.304 | n.s. |
| **Awake vs DOC** | -3.113 | -12.533 | 6.307 | 0.517 | n.s. |
| **Awake vs Moderate Propofol** | -4.943 | -6.593 | -3.293 | p < 0.001 | *** |
| **Mild vs Moderate Propofol** | -5.724 | -7.394 | -4.055 | p < 0.001 | *** |
| **Recovery vs Moderate Propofol** | -8.665 | -10.597 | -6.733 | p < 0.001 | *** |
| **Awake vs fMRI+** | 1.236 | -10.359 | 12.832 | 0.834 | n.s. |
| **Awake vs Mild Propofol** | 0.782 | -0.758 | 2.321 | 0.320 | n.s. |
| **Awake vs Recovery** | 3.722 | 1.904 | 5.541 | p < 0.001 | *** |
| **Placebo vs Ketamine** | 1.798 | 1.425 | 2.172 | p < 0.001 | *** |
| **Placebo vs LSD** | 8.101 | 7.472 | 8.730 | p < 0.001 | *** |

**Table S1.** LME results for total connectome harmonic energy (units are x$10^{-3}$). * $p < 0.05$; ** $p < 0.01$; *** $p < 0.001$;  . $p < 0.10$; n.s. not significant.

| Contrast | Spearman rho | p-value | Significance |
|---|---|---|---|
| **Awake vs fMRI-** | -0.08 | 0.768 | n.s. |
| **fMRI+ vs fMRI-** | 0.12 | 0.662 | n.s. |
| **Awake vs DOC** | -0.24 | 0.384 | n.s. |
| **Awake vs Moderate Propofol** | 0.03 | 0.899 | n.s. |
| **Mild vs Moderate Propofol** | 0.11 | 0.691 | n.s. |
| **Recovery vs Moderate Propofol** | 0.07 | 0.786 | n.s. |
| **Awake vs fMRI+** | -0.33 | 0.230 | n.s. |
| **Awake vs Mild Propofol** | -0.31 | 0.258 | n.s. |
| **Awake vs Recovery** | -0.12 | 0.660 | n.s. |
| **Placebo vs Ketamine** | -0.19 | 0.485 | n.s. |
| **Placebo vs LSD** | -0.15 | 0.590 | n.s. |

**Table S2.** Spearman correlation between the connectome harmonic energy signatures obtained from different states of consciousness (Figures 3 and S4), and the signature obtained from comparing test and retest scans from the same awake volunteers (Figure S2). n.s. not significant.

| Dataset | Source | Fixed Effect | 95% CI Lower | 95% CI Upper | p-value | Significance |
|---|---|---|---|---|---|---|
| **Placebo vs LSD** | propofol | -11.71 | -12.84 | -10.59 | p < 0.001 | *** |
| | DOC | -10.88 | -11.75 | -10.01 | p < 0.001 | *** |
| | ketamine | 5.30 | 4.67 | 5.93 | p < 0.001 | *** |
| | LSD | 6.08 | 5.74 | 6.42 | p < 0.001 | *** |
| **Placebo vs Ketamine** | propofol | -3.00 | -3.74 | -2.26 | p < 0.001 | *** |
| | DOC | -2.70 | -3.24 | -2.16 | p < 0.001 | *** |
| | ketamine | 1.44 | 1.01 | 1.88 | p < 0.001 | *** |
| | LSD | 1.42 | 1.23 | 1.61 | p < 0.001 | *** |
| **Propofol Awake vs Mild** | propofol | -2.16 | -4.62 | 0.31 | 0.087 | . |
| | DOC | -1.88 | -3.88 | 0.13 | 0.067 | . |
| | ketamine | 1.06 | -0.26 | 2.37 | 0.115 | |
| | LSD | 0.89 | 0.04 | 1.75 | 0.040 | * |
| **Propofol Awake vs Recovery** | propofol | -5.08 | -7.92 | -2.25 | p < 0.001 | *** |
| | DOC | -4.62 | -6.94 | -2.30 | p < 0.001 | *** |
| | ketamine | 2.30 | 0.80 | 3.79 | 0.003 | ** |
| | LSD | 2.38 | 1.38 | 3.38 | p < 0.001 | *** |
| **Propofol Awake vs Moderate** | propofol | 6.85 | 4.23 | 9.48 | p < 0.001 | *** |
| | DOC | 5.79 | 3.66 | 7.93 | p < 0.001 | *** |
| | ketamine | -3.45 | -4.84 | -2.06 | p < 0.001 | *** |
| | LSD | -2.88 | -3.79 | -1.97 | p < 0.001 | *** |
| **Propofol Mild vs Moderate** | propofol | 9.01 | 6.33 | 11.69 | p < 0.001 | *** |
| | DOC | 7.67 | 5.49 | 9.85 | p < 0.001 | *** |

| | | | | | | |
|---|---|---|---|---|---|---|
| | ketamine | -4.51 | -5.93 | -3.08 | p < 0.001 | *** |
| | LSD | -3.78 | -4.70 | -2.85 | p < 0.001 | *** |
| **Propofol Recovery vs Moderate** | propofol | 11.94 | 8.90 | 14.97 | p < 0.001 | *** |
| | DOC | 10.41 | 7.93 | 12.90 | p < 0.001 | *** |
| | ketamine | -5.75 | -7.34 | -4.15 | p < 0.001 | *** |
| | LSD | -5.26 | -6.33 | -4.19 | p < 0.001 | *** |
| **DOC fMRI+ vs fMRI-** | propofol | 14.43 | -1.97 | 30.83 | 0.085 | . |
| | DOC | 12.65 | -2.18 | 27.48 | 0.094 | . |
| | ketamine | -7.04 | -14.63 | 0.55 | 0.069 | . |
| | LSD | -6.32 | -13.97 | 1.33 | 0.105 | |
| **Awake vs DOC** | propofol | 3.87 | -9.13 | 16.86 | 0.098 | . |
| | DOC | 2.90 | -8.67 | 14.46 | 0.101 | n.s. |
| | ketamine | -2.38 | -8.52 | 3.77 | 0.105 | n.s. |
| | LSD | -1.13 | -7.00 | 4.74 | 0.109 | n.s. |
| **Awake vs DOC fMRI+** | propofol | -5.31 | -22.34 | 11.72 | 0.112 | n.s. |
| | DOC | -5.16 | -20.17 | 9.85 | 0.116 | n.s. |
| | ketamine | 2.11 | -6.02 | 10.23 | 0.120 | n.s. |
| | LSD | 2.89 | -4.64 | 10.42 | 0.123 | n.s. |
| **Awake vs DOC fMRI-** | propofol | 9.11 | -4.20 | 22.43 | 0.127 | n.s. |
| | DOC | 7.50 | -4.37 | 19.36 | 0.131 | n.s. |
| | ketamine | -4.94 | -11.21 | 1.34 | 0.134 | n.s. |
| | LSD | -3.43 | -9.46 | 2.60 | 0.138 | n.s. |

**Table S3**. LME results for the projection onto the PLS-DA first component of different states of consciousness (units are x$10^{-3}$). * $p < 0.05$; ** $p < 0.01$; *** $p < 0.001$; . $p < 0.10$; n.s. not significant.

| Contrast | Fixed Effect | 95% CI Lower | 95% CI Upper | p-value | Significance |
|---|---|---|---|---|---|
| **Awake vs fMRI-** | -0.074 | -0.129 | -0.018 | 0.009 | ** |
| **fMRI+ vs fMRI-** | -0.063 | -0.133 | 0.007 | 0.079 | . |
| **Awake vs DOC** | -0.051 | -0.102 | 0.000 | 0.051 | . |
| **Awake vs Moderate Propofol** | -0.039 | -0.052 | -0.026 | p<0.001 | *** |
| **Mild vs Moderate Propofol** | -0.040 | -0.054 | -0.027 | p<0.001 | *** |
| **Recovery vs Moderate Propofol** | -0.040 | -0.054 | -0.025 | p<0.001 | *** |
| **Awake vs fMRI+** | -0.011 | -0.064 | 0.042 | 0.690 | n.s. |
| **Awake vs Mild Propofol** | 0.001 | -0.011 | 0.013 | 0.875 | n.s. |
| **Scan 1 vs Scan 2** | 0.001 | -0.013 | 0.014 | 0.925 | n.s. |
| **Awake vs Recovery** | -0.003 | -0.008 | 0.003 | 0.362 | n.s. |
| **Placebo vs Ketamine** | 0.020 | 0.016 | 0.024 | p<0.001 | *** |
| **Placebo vs LSD** | 0.054 | 0.049 | 0.059 | p<0.001 | *** |
| **DOC Any response vs No Response** | -0.08 | -0.15 | -0.01 | 0.024 | * |

**Table S4.** LME results for repertoire diversity of the connectome harmonics. * $p < 0.05$; ** $p < 0.01$; *** $p < 0.001$;  ** $p < 0.01$;  . $p < 0.10$; n.s. not significant.

| Contrast | Fixed Effect | 95% CI Lower | 95% CI Upper | p-value | Significance |
|---|---|---|---|---|---|
| **Awake vs fMRI-** | 0.023 | -0.021 | 0.067 | 0.299 | n.s. |
| **fMRI+ vs fMRI-** | 0.023 | -0.030 | 0.077 | 0.394 | n.s. |
| **Awake vs DOC** | 0.015 | -0.026 | 0.055 | 0.473 | n.s. |
| **Awake vs Moderate Propofol** | -0.017 | -0.037 | 0.003 | 0.099 | . |
| **Mild vs Moderate Propofol** | -0.010 | -0.030 | 0.010 | 0.334 | n.s. |
| **Recovery vs Moderate Propofol** | 0.011 | -0.010 | 0.032 | 0.303 | n.s. |
| **Awake vs fMRI+** | 0.000 | -0.049 | 0.049 | 0.996 | n.s. |
| **Awake vs Mild Propofol** | -0.007 | -0.025 | 0.011 | 0.454 | n.s. |
| **Scan 1 vs Scan 2** | -0.028 | -0.046 | -0.009 | 0.003 | ** |
| **Awake vs Recovery** | -0.001 | -0.015 | 0.012 | 0.836 | n.s. |
| **Placebo vs Ketamine** | -0.007 | -0.016 | 0.003 | 0.164 | n.s. |
| **Placebo vs LSD** | -0.014 | -0.023 | -0.005 | 0.002 | ** |

**Table S5.** LME results for repertoire diversity of the connectome harmonics, from rotated harmonics. * $p < 0.05$; ** $p < 0.01$; *** $p < 0.001$; ** $p < 0.01$; . $p < 0.10$; n.s. not significant.

| Contrast | Fixed Effect | 95% CI Lower | 95% CI Upper | p-value | Significance |
|---|---|---|---|---|---|
| **Awake vs fMRI-** | -0.030 | -0.044 | -0.015 | p<0.001 | *** |
| **fMRI+ vs fMRI-** | -0.039 | -0.053 | -0.025 | p<0.001 | *** |
| **Awake vs DOC** | -0.016 | -0.028 | -0.003 | 0.015 | * |
| **Awake vs Moderate Propofol** | -0.018 | -0.034 | -0.003 | 0.023 | * |
| **Mild vs Moderate Propofol** | -0.017 | -0.033 | -0.001 | 0.039 | * |
| **Recovery vs Moderate Propofol** | -0.001 | -0.018 | 0.016 | 0.921 | n.s. |
| **Awake vs fMRI+** | 0.009 | -0.004 | 0.022 | 0.163 | n.s. |
| **Awake vs Mild Propofol** | -0.001 | -0.015 | 0.013 | 0.862 | n.s. |
| **Scan 1 vs Scan 2** | -0.017 | -0.032 | -0.002 | 0.023 | * |
| **Awake vs Recovery** | -0.003 | -0.012 | 0.006 | 0.493 | n.s. |
| **Placebo vs Ketamine** | 0.003 | -0.003 | 0.009 | 0.279 | n.s. |
| **Placebo vs LSD** | -0.003 | -0.009 | 0.003 | 0.335 | n.s. |

**Table S6.** LME results for repertoire diversity of the connectome harmonics, from a randomised connectome. * $p < 0.05$; ** $p < 0.01$; *** $p < 0.001$;  ** $p < 0.01$;  . $p < 0.10$; n.s. not significant.

| Contrast | Fixed Effect | 95% CI Lower | 95% CI Upper | p-value | Significance |
|---|---|---|---|---|---|
| **Awake vs fMRI-** | -0.062 | -0.107 | -0.017 | 0.007 | ** |
| **fMRI+ vs fMRI-** | -0.070 | -0.124 | -0.016 | 0.012 | * |
| **Awake vs DOC** | -0.037 | -0.080 | 0.006 | 0.095 | . |
| **Awake vs Moderate Propofol** | -0.036 | -0.050 | -0.022 | p<0.001 | *** |
| **Mild vs Moderate Propofol** | -0.039 | -0.054 | -0.025 | p<0.001 | *** |
| **Recovery vs Moderate Propofol** | -0.034 | -0.050 | -0.018 | p<0.001 | *** |
| **Awake vs fMRI+** | 0.008 | -0.039 | 0.054 | 0.749 | n.s. |
| **Awake vs Mild Propofol** | 0.003 | -0.009 | 0.016 | 0.605 | n.s. |
| **Scan 1 vs Scan 2** | -0.002 | -0.016 | 0.012 | 0.793 | n.s. |
| **Awake vs Recovery** | 0.000 | -0.006 | 0.006 | 0.996 | n.s. |
| **Placebo vs Ketamine** | 0.015 | 0.011 | 0.020 | p<0.001 | *** |
| **Placebo vs LSD** | 0.039 | 0.033 | 0.044 | p<0.001 | *** |

**Table S7.** LME results for repertoire diversity of the connectome harmonics, obtained from the HCP-985 connectome. * $p < 0.05$; ** $p < 0.01$; *** $p < 0.001$; ** $p < 0.01$; . $p < 0.10$; n.s. not significant.

| Contrast | Fixed Effect | 95% CI Lower | 95% CI Upper | p-value | Significance |
|---|---|---|---|---|---|
| **Awake vs fMRI-** | -0.062 | -0.109 | -0.015 | 0.010 | ** |
| **fMRI+ vs fMRI-** | -0.071 | -0.128 | -0.014 | 0.015 | * |
| **Awake vs DOC** | -0.036 | -0.082 | 0.009 | 0.114 | n.s. |
| **Awake vs Moderate Propofol** | -0.037 | -0.051 | -0.023 | p<0.001 | *** |
| **Mild vs Moderate Propofol** | -0.039 | -0.054 | -0.025 | p<0.001 | *** |
| **Recovery vs Moderate Propofol** | -0.034 | -0.050 | -0.018 | p<0.001 | *** |
| **Awake vs fMRI+** | 0.009 | -0.041 | 0.058 | 0.735 | n.s. |
| **Awake vs Mild Propofol** | 0.002 | -0.010 | 0.015 | 0.709 | n.s. |
| **Scan 1 vs Scan 2** | -0.003 | -0.017 | 0.011 | 0.681 | n.s. |
| **Awake vs Recovery** | 0.001 | -0.005 | 0.007 | 0.738 | n.s. |
| **Placebo vs Ketamine** | 0.015 | 0.011 | 0.020 | p<0.001 | *** |
| **Placebo vs LSD** | 0.043 | 0.038 | 0.049 | p<0.001 | *** |

**Table S8.** LME results for repertoire diversity of the connectome harmonics, obtained from the MGH-32connectome. * $p < 0.05$; ** $p < 0.01$; *** $p < 0.001$;  . $p < 0.10$;  n.s. not significant.
